# Pig herd management and infection transmission dynamics: a challenge for modellers

**DOI:** 10.1101/2023.05.17.541128

**Authors:** Vianney Sicard, Sébastien Picault, Mathieu Andraud

## Abstract

The control of epidemics requires a thorough understanding of the complex interactions between pathogen transmission, disease impact, and population dynamics and management. Mechanistic epidemiological modelling is an effective way to address this issue, but handling highly structured and dynamic systems, remains challenging. We therefore developed a novel approach that combines Multi-Level Agent-Based Systems (MLABS) with spatial and temporal organization, allowing for a tuned representation of the transmission processes amongst the host population. We applied this method to model the spread of a PRRSv-like virus in pig farms, integrating the clinical consequences (conception and reproduction failures), in terms of animal husbandry practices. Results highlighted the importance to account for spatial and temporal structuring and herd management policies in epidemiological models. Indeed, disease-related abortions, inducing reassignments of sows in different batches, was shown to enhance the transmission process, favouring the persistence of the virus at the herd level. Supported by a declarative Domain-Specific Language (DSL), our approach provides flexible and powerful solutions to address the issues of on-farm epidemics and broader public health concerns. The present application, based on a simple Susceptible-Exposed-Infected-Recovered (SEIR) model, opens the way to the representation of more complex epidemiological systems, including more specific features such as maternally derived antibodies, vaccination, or dual infections, along with their respective clinical consequences on the management practices.

## 1 INTRODUCTION

Understanding the mechanisms of pathogen spread in a highly structured population is a key element for epidemic control. This requires capturing the spatial and social structures, defining the contact rates between individuals and groups of hosts in the population. Livestock management clearly illustrates that problematic, obeying predefined rules ensuring a balance between animal welfare, good sanitary conditions, and productivity. The spread of pathogens in a pig farm, managed with batch-rearing procedures, therefore represents an ideal application for the integration of an innovative organizational pattern within a multi-level agent-based modelling framework dedicated to epidemiological modelling (Sicard et al., 2021b).

To further understand the impact of batch management, housing, and possible deviations in herd management practices on the spread of pathogens at different scales, and to identify realistic levers, new modelling approaches had to be developed. In the era of open and reproducible science, ensuring legibility, revisability, and flexibility of models is pivotal. The response provided by the EMULSION framework, based on AI methods and complemented with an organizational pattern, offers solutions to epidemiological modelling issues (Picault et al., 2019).

In a previous study, we first developed a model of swine influenza A in pig farms, highlighting the impact of the spatio-temporal structure of the herd on the transmission dynamics and its impact on virus spread and control based on EMULSION extended with an organizational pattern (Sicard et al., 2021a). However, in this study, influenza infections were not considered to have any consequence in terms of animal management. Therefore, the present study aims to account for the interplay between infectious dynamics, clinical consequences, and management practices. For this purpose, we developed an epidemiological model representing the spread of a disease transmitted by direct contact between animals (e.g. porcine reproductive and respiratory syndrome virus or porcine circovirus of type 2), and assuming clinical reproductive consequences in sows leading to modifications of the management rules. This assumption would reflect the situation early after introduction of Porcine Reproductive and Respiratory Syndrome (PRRSv) virus in a farm.

To ensure realistic epidemiological modelling regarding field situations, including the contact structure between animals and different observation levels (e.g. farm, herd, individual), agent-based simulation (ABS) has proven its value (Roche et al., 2008). ABS provides methods for handling behaviours, interactions, and tracking of individuals. For further explicit representation of complex systems, multi-level agent-based simulation (MLABS) allows several scales (individuals, groups, batches, populations, etc.) to be associated with agents endowed with their own behaviours (Mathieu et al., 2018). Besides, MLABS makes it possible to separate procedural knowledge (calculations and processes involved in stochastic epidemiological models, according to the modelling paradigm and scale: e.g. compartmental models, individual-based, metapopulations), from declarative knowledge (model structure, assumptions, description of groups and processes, parameter values, initial conditions), as set out in symbolic AI (Weyns, 2005; Mathieu et al., 2015, 2018). This separation of concerns provides the ability to make models modular and easy to use by defining independent processes (e.g. infection, population dynamics, trade movements, detection) which can be coupled, rather than a representation with a single, huge and tangled flow diagram.

We propose a prototype model architecture accounting for the complex interplay between pathogen transmission dynamics and consequences of clinical cases on herd management, by coupling a mechanistic multi-level agent-based modelling approach (EMULSION framework) with specific organizational considerations, including exceptions in management practices related with clinical consequences of infections in reproductive sows. The EMULSION modelling framework enables the specification of different scenarios by varying the population dynamics in the breeding sectors (e.g. batch management, exceptions). These scenarios were used to assess the impact of clinical outcomes of infectious diseases on population and transmission dynamics at the herd level. We illustrate this approach through a PRRSv-like disease spreading in a fine-grained realistic pig farm model.

## 2 MATERIALS AND METHODS

### 2.1 Model overview

The management of the involved batches aligned with the procedures described in (Sicard et al., 2022), such as sector allocation according to physiological state durations. To represent the clinical reproductive consequences of infections in sows, a probability of insemination failure was considered, leading to potential batch downgrading for infected sows. Infection consequences were modulated upon different periods of gestation with specific impacts on the health status of piglets at birth, including abortions, vertical transmission, and maternally derived antibodies delivery. Furthermore, the model was able to represent batch management at a fine-grained level, encompassing both litter and pen levels (Fig. 1), thus providing the ability to represent zootechnical practices such as adoptions, pig gathering procedures, and sow renewal process.

**Figure 1:**
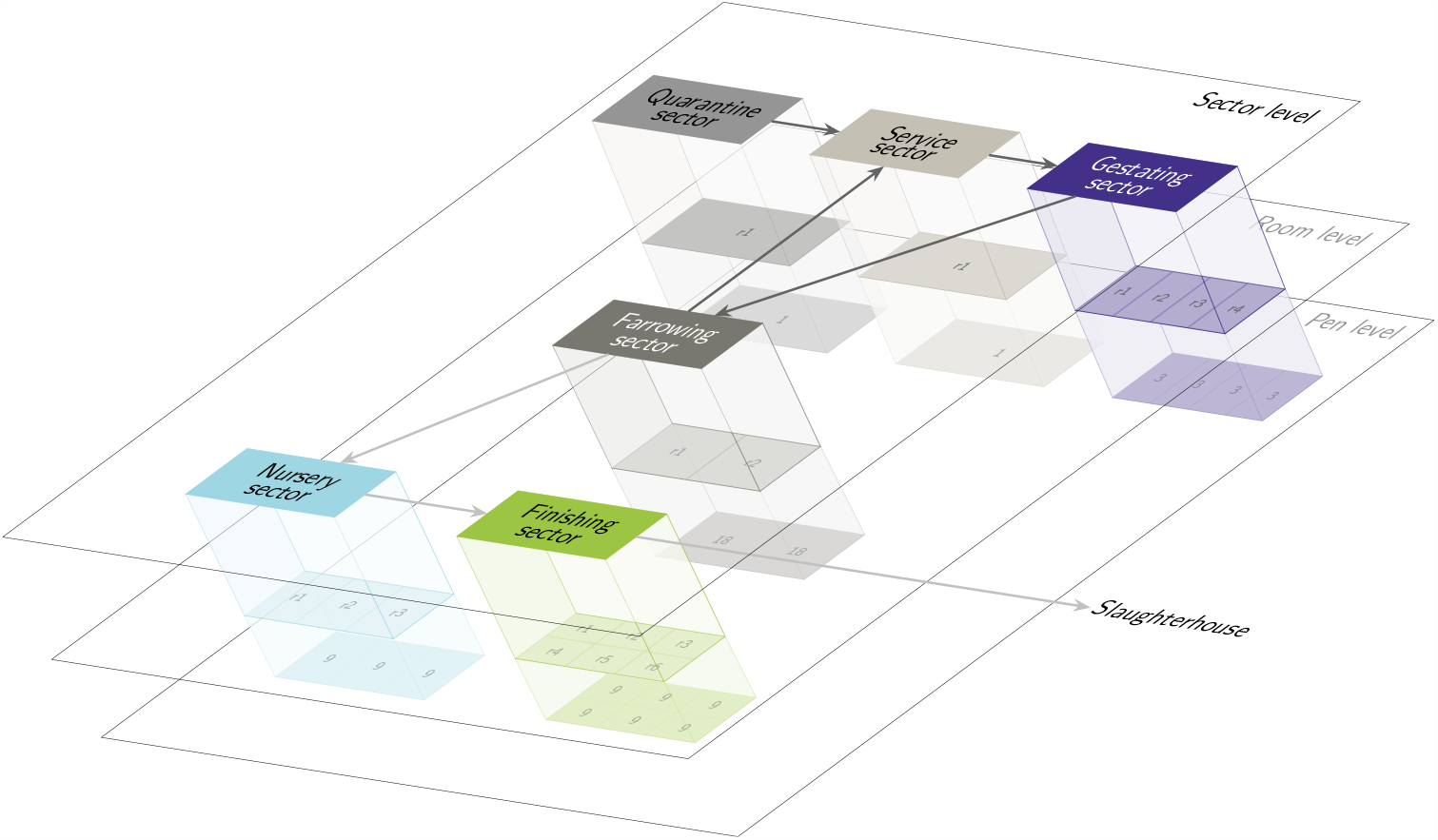
Representation of the three spatial levels: Sector, Room, and Pen levels. Each sector (Sector level) is further divided into rooms (*r*_*n*_ at the Room level), and each room is further subdivided into pens (Pen level, where the numbers correspond to the number of pens per room).

### 2.2 EMULSION Framework

The model was developed using the EMULSION framework (Picault et al., 2019), which is dedicated to stochastic mechanistic epidemiological modelling. An essential concept of the framework is the separation between knowledge representation (the model as a structured text), and the simulation processes (provided by a generic engine which reads and executes the model description).

The EMULSION framework, extended with its organizational component (Sicard et al., 2021b), was used to account for the complex herd structure in both space and time, including multilevel aspects. An organization is an entity made up of groups to which individuals belong. Organizations and groups encapsulate environments that correspond to spaces where agents are located (either groups or individuals). The organizational pattern systematically describes three levels: organization, group, and individual.

A particular feature of the pattern is its ability to be used recursively. In other words, a group can itself be an organization (thus becoming a sub-organization) (Figure 2), thereby describing multi-level dynamics and relationships between levels.

**Figure 2:**
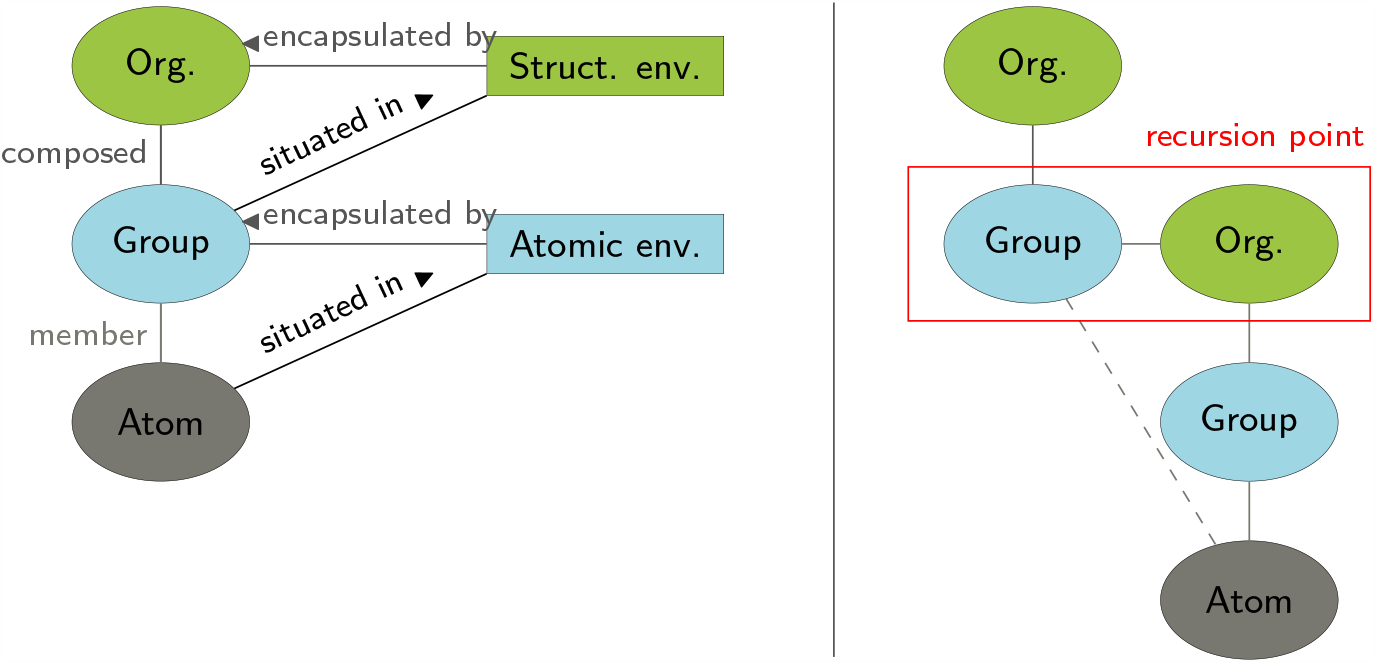
Structure of the organizational pattern. The agent *Organization* encapsulates an environment where agents *Group* are situated. Each agent *Group* encapsulates an environment where the atomic agents (atom) are situated. The pattern can be used recursively, i.e. an agent *Group* can itself be an organization. The environment in the pattern can be either spatial or social. (Sicard et al., 2021b)

### 2.3 Population dynamics

The pig production herd was raised according to a seven-batch-rearing system with a 21-day between-batch interval, and all-in-all-out procedures, the main management practice held in France, which was fully described in Sicard et al. (2022). Two subpopulations, breeding sows and growing pigs, were represented, structured and managed according to husbandry constraints (d’Agriculture de Bretagne, 2010). The batches remained consistent, i.e. all animals remain within the same batch throughout their life cycle, being in the same physiological state at the same time and obeying an all-in-all-out procedure.

Sows evolved through three physiological states over 147 days: insemination (34 days), gestating (85 days) and farrowing/lactating (28 days). Growing pigs evolved through three stages over 182 days before being sent to the slaughterhouse: farrowing/suckling (28 days), post-weaning (40 days) and finishing (114 days) (Figure 3). Each physiological stage corresponded to a specific physical sector of the farm. In the field, the duration spent in different states may vary, especially due to variations in parturition timing. These variations can occur within a time window of two days, either before or after the predicted date. However, the practice of all-in-all-out remains observed on farms. This means that the timing of animal movement can be deterministically scheduled in the model.

**Figure 3:**
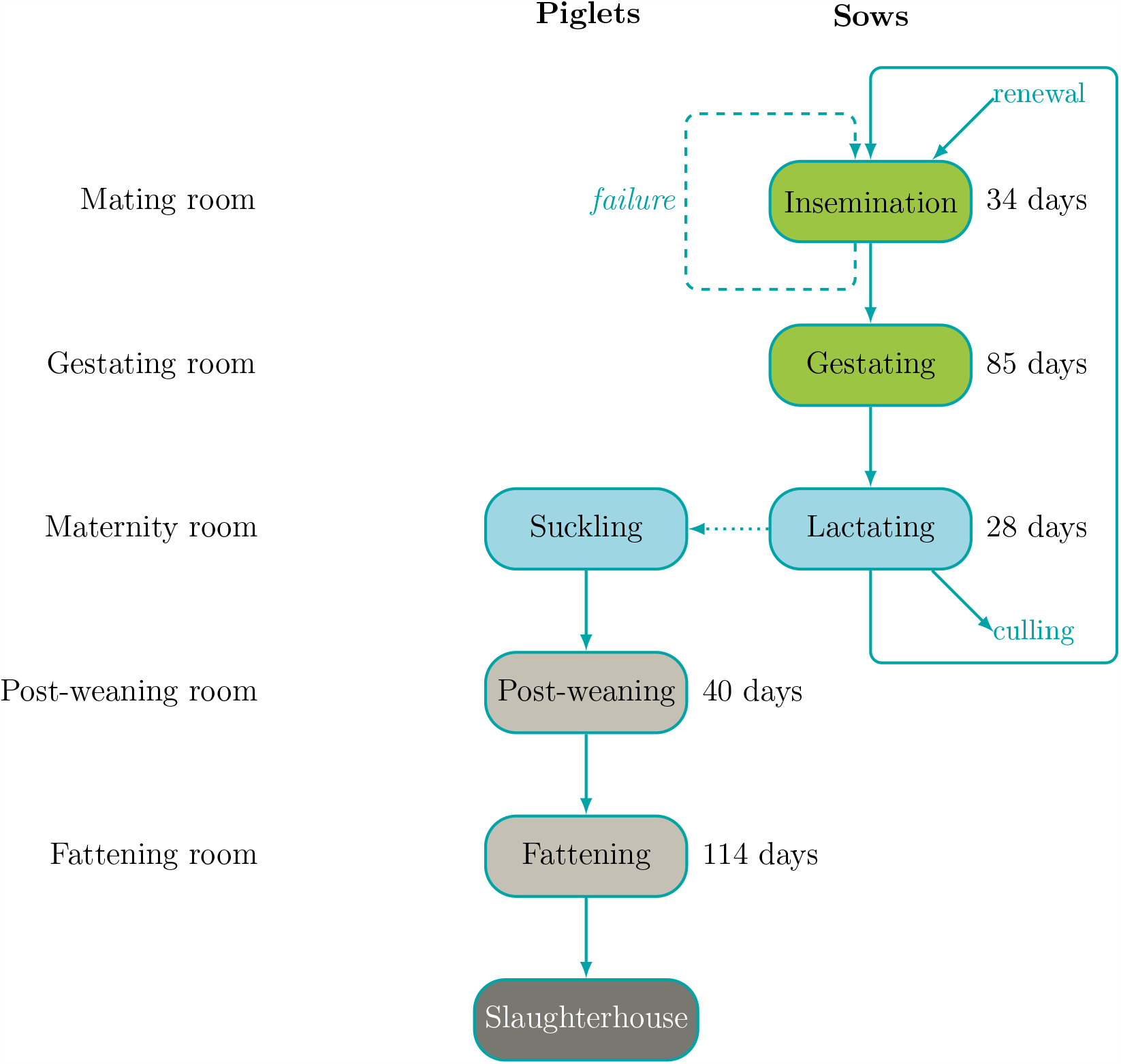
Illustration of the duration of physiological stages in relation to housing. The dotted arrow represents piglet production from sows and the dashed one represents sows that are retrograded to the previous batch in case of insemination failure.

The *housing* organization ensures that the spatial sectorial structure of the housing is adequate for the physiological needs of the individuals. The sectorial organizations are divided into downstream levels corresponding to the rooms in the herd. Each level was considered an organization, which could be further subdivided into downstream sublevels. The rooms are subdivided into pens to represent the direct and indirect contact of animals within and between pens (Fig. 4).

**Figure 4:**
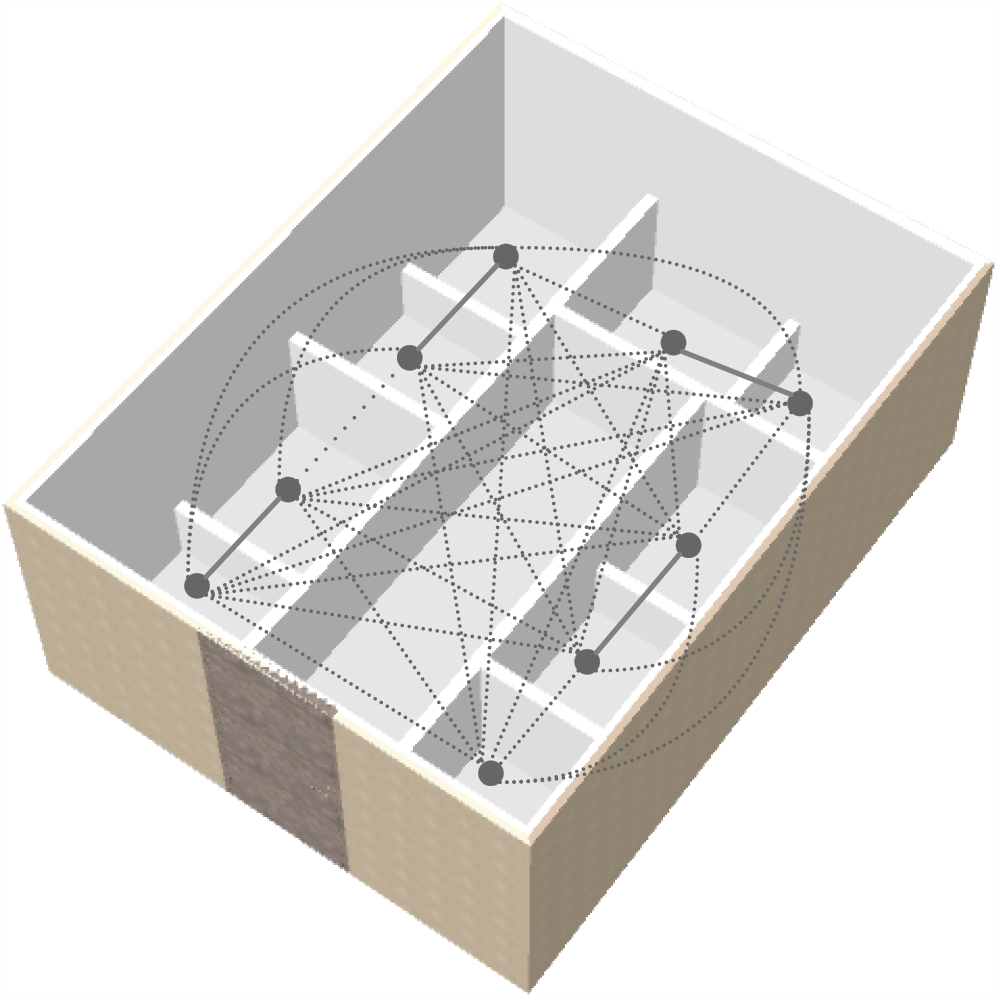
Representation of a room in the finishing sector: physical division into pens, including a network of contacts. It illustrates the model’s ability to represent fine-grained contact networks, both direct contacts between adjacent pens, indicated by solid lines, and indirect contacts with other pens, shown as dotted lines.

Batches were managed by several organizations: *Batches* for main herd management, *Litter* for making the link between breeding sows and their piglets, *Litter_group* for gathering purpose in nursery pens, and *Housing* specifying the exact location of each animal (or group of animals) in corresponding sectors, rooms, and pens.

At initialization, 98 (14 sow per batch) sows were homogeneously distributed among the 7-batch organization. At farrowing, newborn piglets were assigned to the same batch and same pen as their mother. The *litter* organization was designed as the dams and their relative piglets, located in the same space (pens) in farrowing rooms. The pen organization level included determining the number of pens required to accommodate all sows in case of overcrowding due to insemination failures or gestation-related abortions in other batches (Figure 1). Sows were alternatively placed in designated organizational spaces, while piglets were housed in the same pen as their respective mothers during farrowing.

*litter_group* was designed to represent pig gathering policy in nursery rooms after weaning. Indeed, pig litters are frequently gathered into larger groups within pens when entering the nursery sector. In our model, it was assumed that each pen in the nursery rooms housed two litters from the farrowing rooms. The location of the litters within the pens was managed by one of the *housing* sub-organisations.

Herd renewal might be a factor influencing disease spread by introducing susceptible animals into the system. In commercial pig production, sows are selected for replacement based on productivity criteria. For example, old sows are known to have smaller litters, and farmers cannot afford to keep sows that experience multiple gestation failures. In the present model, for the sake of illustration, the decision to cull based on their condition (*condition_to_cull*) was made :

- sows with parity higher than 5, or
- sows with more than 5 gestation failures.

The maximum number of sows that could be replaced at each batch-cycle was set to 2 and adjusted to ensure a stable population in each batch, ranging between the initial number of sows in the herd (14 sows per batch) and the total number of pens in farrowing room (18 pens). Sow replacement was managed as follows: a sow with parity rank higher than 5, or after two unsuccessful inseminations, could be culled. The adjustment of the number of sows was evaluated regardless of the reason for replacement, due to culling. The renewal was managed by the *culling* state machine accounting for three states: *to_keep* corresponding to sows that will be not renewed, *to_cull* corresponding to sows that will be renewed, and *culled* corresponding to sows that leave the system. The decision on the status of the sows is made when entering in farrowing room; after the lactation period, sows are culled and replaced by renewal susceptible gilts for the next reproduction cycle.

Insemination failure was represented by the state machine *inseminationStatus*. On entering the insemination sector (corresponding to the gestating state of the *physiologicalStep* state machine), sows were inseminated (inseminate status of the statemachine *inseminationStatus*). 42 days (i.e., two batch intervals) after insemination, sows are checked for gestation, with a failure probability defined as *proba_failure_ins*. Sows that were successfully inseminated were moved to *SuccessInf* state. Sows that had failed insemination were retrograded from two previous batches. Owing to the 7 batch-farrowing system, an insemination failure for a sow in batch *N* led to a transfer into batch *N −* 2, for new insemination attempt. Such event will be next mentioned as a batch reassignment event.

### 2.4 Health states

Four health states were considered: animals protected by maternal immunity (M), susceptible (S), infectious (I), and recovered (R). These health states were managed by the state machine *health_state* (Figure 5). The state *M*, corresponding to a piglet protected by maternal immunity, was managed separately by the state machine *maternal_immunity*. The epidemiological parameters were based on a PRRSv-like disease, and are reported in the Table 1. Parameter *β*_*ind*_ represented the rate of transmission resulting from occasional contacts between individuals from adjacent pens. Parameter *β* represented the direct transmission between individuals. Parameter *γ* corresponded to the recovery rate, i.e., the rate at which individuals become recovered (R). The duration in state M was distributed according to a gamma distribution, after which pigs became fully susceptible.

**Table 1:**
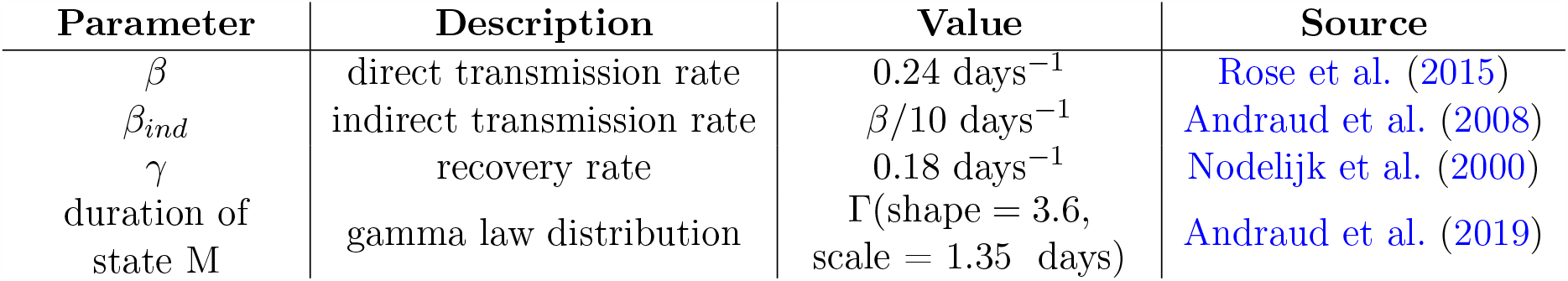
Table of epidemiological parameter values.

| Parameter | Description | Value | Source |
| --- | --- | --- | --- |
| $\beta$ | direct transmission rate | 0.24 days<sup>-1</sup> | Rose et al. (2015) |
| $\beta_{ind}$ | indirect transmission rate | $\beta/10$ days<sup>-1</sup> | Andraud et al. (2008) |
| $\gamma$ | recovery rate | 0.18 days<sup>-1</sup> | Nodelijk et al. (2000) |
| duration of state M | gamma law distribution | $\Gamma(\text{shape} = 3.6, \text{scale} = 1.35 \text{ days})$ | Andraud et al. (2019) |

**Figure 5:**
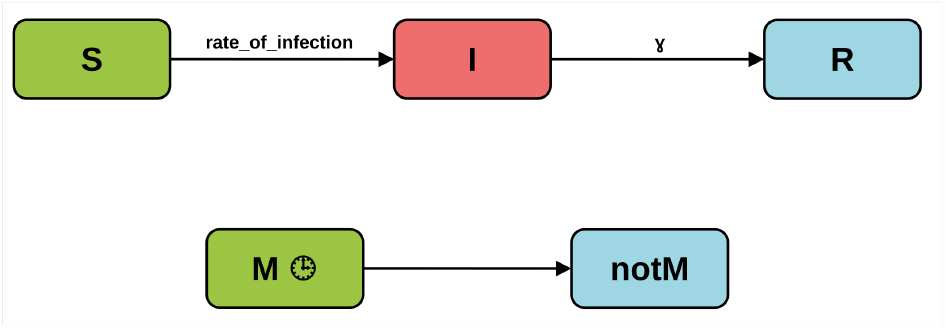
Representation of the two state machines: *health_state* (on the top) and *maternal_- immunity* (on the bottom). A susceptible individual (S) becomes exposed (I) with a rate (rate_- of_infection), and then recovers (R) with a rate *γ*). Piglets with maternal immunity (M) stay in the state M for a duration described in Table 1 and then lose their immunity. Maternal immunity provides protection against infection.

To account for the complex spatial structure of the animal housing, calculating the force of infection, including the direct pairwise contact in the pens, was a cumbersome task. To facilitate the representation of transmission routes and increase the flexibility of the contact structure, a graphic-based approach was adopted. Specifically, the housing system was represented as a graph, where the nodes corresponded to the physical environmental spaces such as pens, corridors, and rooms, and the links represented the interactions between these spaces. The links were weighted by the epidemiological information of each space. This approach provided a comprehensive and flexible representation of the transmission dynamics within the housing system (Fig. 4).

Specifically, each space in the housing system was associated with the number of infectious animals it contained, and this information was propagated through the graph. The cumulative information obtained from the graph was then used to compute the force of infection:

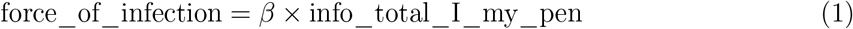

where *info_total_I_my_pen* corresponded to a function that the EMULSION framework automatically generates. This function retrieves the value of the total number of infectious individuals (*total_I*) of the current space (*my_pen*).

Piglets’ epidemiological status at birth was assumed dependent on the period of gestation the sows were infected. Sow infected in the first third of their gestation produce susceptible piglets; Sow infected in the second third of their gestation produce piglets infected through transplacental transmission; finally, piglets born to sows infected in the last phase of the gestation period acquire maternally derived antibodies in the “M” state.

### 2.5 Scenarios

Four scenarios were established to represent several types of management, taking into account the exchange of sows between batches in case of insemination failure or abortion, the type of grouping in gestating, and the rate of occasional contact (Table 2). Each scenario was run with 100 stochastic replicates over a period of 1500 days, including a burning period of two reproductive cycles (2 *×* 147days). The virus was introduced through an infected sow in batch 1, at the beginning of the third reproductive cycle of the batch, assuming the introduction of an infected gilt into the system (294 days). In terms of the allocation process, the gilt was assigned to litter 1, indicating its location in room 1, pen 1 within the gestating sector.

**Table 2:** Description of the four scenarios. Each scenario corresponds to a combination of grouping type in the gestation sector (grouped by 6 or All-in-One), and batch management policies (batch reassignment due to insemination failure or batch stability)

| Scenario | Description |
| --- | --- |
| Scenario 1 | Sows grouped by 6 without batch reassignment |
| Scenario 2 | Sows grouped by 6 with batch reassignment |
| Scenario 3 | All-in-One grouped sows without batch reassignment |
| Scenario 4 | All-in-One grouped sows with batch reassignment |

## 3 RESULTS

The results provided a measure of the impact of management changes (grouping) of breeding sows on the infectious dynamics. Scenarios with no grouping in the gestation sector showed a weak impact of batch changes due to insemination failure. For the scenarios with *β*_*ind*_ of 0 and *β/*10, the dynamic profiles were similar (Figure 6).

**Figure 6:**
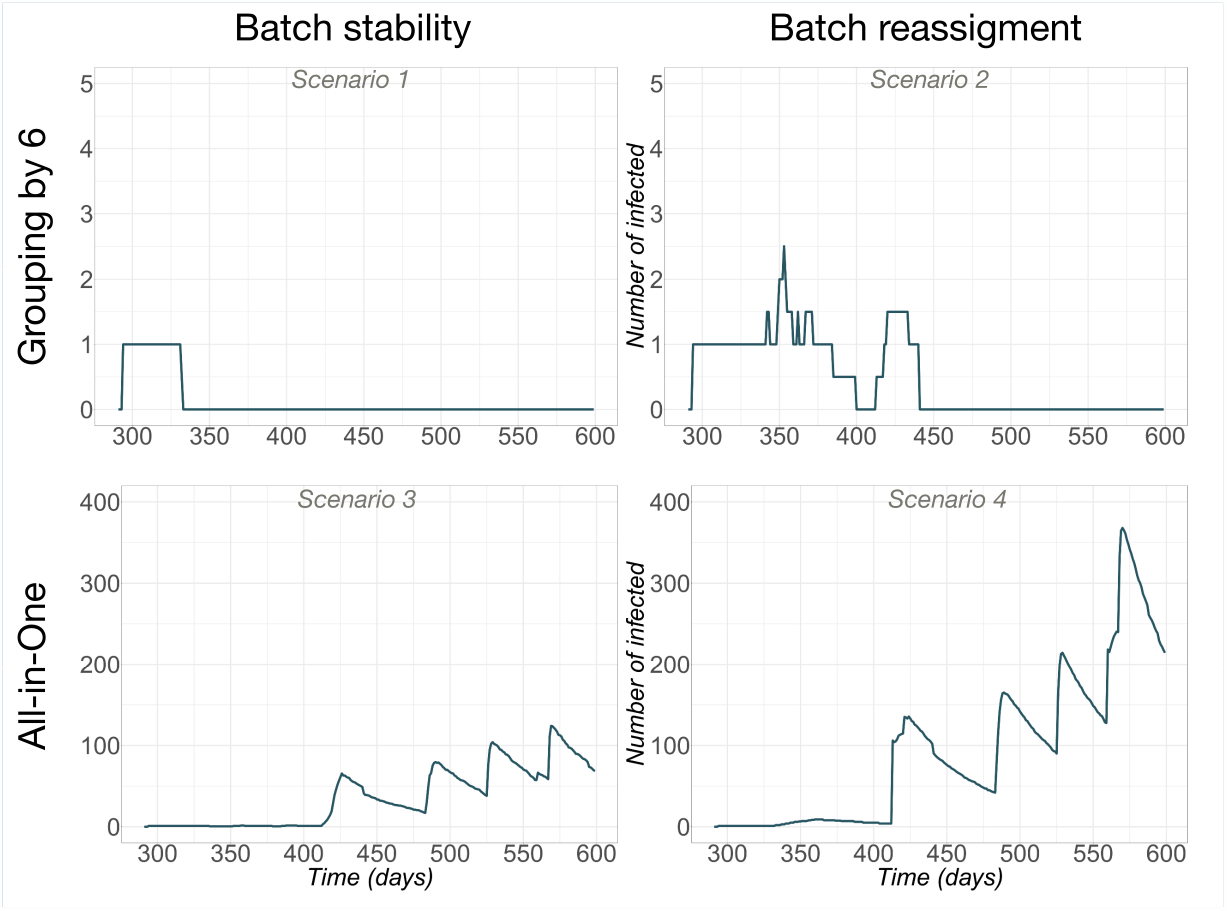
Infection dynamics for scenarios with grouping in gestating sector. This shows the median number of infected sows over time across the whole farm after the introduction of an infected sow in batch 1 at 294 days. The time scale started at 294 days, which corresponds to the date of introducing an infectious gilt into the system, following the burning period.

In the scenarios with *β*_*ind*_ of 0 (scenarios 3 and 4), the first peak occurred approximately 34 days after the virus was introduced and corresponded to the entry into the gestation sector. As sows in the same batch were mixed, the virus spread rapidly within the batch. The second peak, about 85 days after the first, corresponded to farrowing sector. The following peak, about 28 days later, corresponded to piglets entering the post-weaning sector. In the farrowing sector, each pen contained one litter consisting of sow and her piglets, and the pens were assumed independent, with no possible contact with neighbouring pens. With a *β*_*ind*_ value of 0, only within-pen transmission occurred. Conversely, in the post-weaning sector, where pens were not independent, litters were grouped in pairs in pens, and transmission occurred between adjacent pens with *β*_*ind*_ = *β/*10. This demonstrated batch changes for infected sows following failed insemination in the *n −* 2 batch in the insemination sector broke the infectious dynamic within the batch.

The duration of infection was 56 days, which is longer than the period between entry into the insemination sector and the detection of insemination failure (42 days). Therefore, a retrograded infected sow could potentially infect other individuals, at least in the insemination sector, and potentially as far as the gestating sector (if a sow was infected a few days before being retrograded, it would arrive in the gestating sector of its new batch at 36 days, and could be infectious for about 20 days). For the scenarios without grouping, changes to nominal batch-rearing management did not have a notable impact on the transmission dynamics. For the scenarios with grouping (per 6) in the gestating sector, exceptions to batch rearing substantially impacted the infection dynamics, with transmissions occurring between non-adjacent pens in fully susceptible groups of sows.

For the scenarios with a value of *β*_*ind*_ of 0.24 (scenarios 1 and 2) (Figure 6), the first peak corresponded to the entry into the gestating sector. Due to the grouping by 6 of the sows in this sector, the batch reassignment strongly impacted the transmission dynamics, with higher peaks of incidence due to the potential introduction of infected sows in fully susceptible pens.

## 4 DISCUSSION

We developed a model to better understand the interplay between social and spatial organization of a pig production herd and infection dynamics. Indeed, several pathogens (e.g. PRRSv, PCV-2, influenza viruses, *Mycoplasma hypopneumoniae*) can yield to deviations in herd management due to clinical expressions in pigs, impairing the nominal management to be operated. The complex interactions between disease management strategies and their impact on herd organization, together with the dynamics of disease spread, have not yet been fully accounted for in the current model due to the complexity it introduces in the model design. To achieve this, we applied an artificial intelligence-based approach (Ezanno et al., 2020). We introduced an original multi-level agent-based design pattern to capture organizational features involved in the complexity of highly structured populations in time and space. This approach was associated with a dedicated modelling language to facilitate the specification of such organizations without writing computer code, and it was integrated into the EMULSION framework. Our approach facilitated the representation of the complex spatio-temporal herd structure, enabling us to conduct a comprehensive study of the system.

The integration of AI methodologies into epidemiological modelling, including simulation architecture and knowledge representation methods, extends the capabilities of epidemiological models. This approach, through MLABS enhanced with organizational concerns (OMLABS), allows for the representation of mechanisms previously unconsidered in the field. OMLABS allows the study of various scales in a single simulation, facilitating a detailed analysis of the impact of each level in the overall dynamic. This, in turn, allows for a focused identification of effective measures and provides specific recommendations for action.

A literature review highlighted the role of mathematical models as tools for improving the understanding of viral infection spread in pig production units (Andraud and Rose, 2020). Swine influenza virus, PCV-2, or hepatitis E virus were indeed fields of application for models with different paradigms, from individual-based approaches to population-based models, depending on the transmission characteristics of the pathogens (Reynolds et al., 2014; White et al., 2017; Andraud et al., 2008; Salines et al., 2020).

Multi-level Agent-based Systems provide solutions for unifying these modelling paradigms using a common framework, with high Readability, Reproducibility, and Flexibility (RRF). The EMULSION framework provides real readability (R) through its specific language (DSL) in a “no-code” approach and ensures, through its internal structure, reproducibility (R) (Picault et al., 2019). The add-on of our organizational design-pattern offers the opportunity to tackle population structures with fine-grained representations of their epidemiological consequences with great flexibility (F), keeping the above-mentioned (RRF) advantages (Sicard et al., 2021b). This was recently illustrated with a model representing the transmission of a swine influenza A virus in a pig herds with different spatial configurations (Sicard et al., 2022).

Previous models represented the impact of management practices on the transmission dynamics, to assess how husbandry practices could be modified to better handle health issues at the herd level (Cador et al., 2016; Suksamran et al., 2017) Clinical aspects and their potential consequences, in terms of management and infection dynamics, were nevertheless rarely considered. PRRS virus is recognized as a major economic burden for swine producers, inducing huge loss due to growth retardation, abortions or reproduction failures, birth of stillborn fetuses or weak-condition piglets (Le Coz, 2007; Renken et al., 2021; Charpin et al., 2012). Such clinical outcomes necessarily induce exceptions in the management of the herds, affecting, in turn, the transmission dynamics. We therefore developed a model, accounting for organizational features, to study the impact of such disease-related exceptions on the transmission of an infectious agent within a pig herd.

The model primarily focuses on representing the dynamics in the breeding sectors and their impact on the whole farm. Our Results highlight the roles played by various transmission routes, including batch change of sows after insemination failure, sow grouping in the gestating sector, and indirect contact rate (*β*_*ind*_), representing a potential airborne transmission route (Cador et al., 2016). Introduction of infectious sows in susceptible batches clearly increased the risk of persistence of the virus on farm.

Husbandry practices were highlighted as increasing the risk of transmission at group level and persistence at herd level for different infectious agents, from both field observations and synthetic data (Walachowski et al., 2014; Fablet et al., 2013; Cador et al., 2017). However the role of deviations in nominal batch management due to clinical expressions, such as reproductive issues in breeding sows or growth retardation in pigs, was not objectified. This could nevertheless favour the transmission between groups of pigs which could deserve further investigation to develop farm-specific control solutions. Our model identified the primary transmission routes and the impact of reinfection on the breeding population. These factors could lead to infection in maternity piglets and, ultimately, active immunity that could lead to virus extinction. While establishing effective immunity in piglets with maternal immunity can be challenging, our previous research highlighted its potential as a control measure for Influenza A (Sicard et al., 2022).

This study considered only deviations of management due to insemination failures, which could correspond to clinical consequences for sows infected by PRRSv during early gestation after introduction of the virus on farm. We did not consider the potential for adoption between litters, whether within the same batch or from different batches. However, this aspect could be a topic for future research. However, the consequences of PRRSv infections are dramatic for animals of all physiological stages. Late abortions would lead to mixing animals from different batches. Culling rate might also be increased, inducing the introduction of renewal gilts, feeling the pool of susceptible animals. In growing pigs, birth of piglets of weak condition, or growth retardation due to infections could favour mingling of animals from different litters in nursery pens. These impacts may have indirect consequences on herd management, leading to exceptions in nominal management, such as mixing animals at different physiological stages. Our study showed that exceptions due to batch reassignment for sows played an important role in infectious dynamics. Our proposed methodological solution, which used artificial intelligence and organizational design patterns for multi-level agent-based simulation, provides a concrete way to represent these phenomena. However, the model will need to be extended to account for the virus impact on the management of exceptions at whole farm scale.

## 5 ACKNOWLEDGEMENTS

This work is supported by a grant from INRAE (Animal Health Division) and the French Region Pays de la Loire. We are grateful to the INRAE MIGALE bioinformatics facility (MIGALE, INRAE, 2020. Migale bioinformatics Facility, DOI: 10.15454/1.5572390655343293E12) for providing computing resources.

