## Supplementary material for "Pig herd management and infection transmission dynamics: a challenge for modellers": SDRP_like.yaml

```
model_name: test-total

model_info:
  abstract: '<description>'

time_info:
  time_unit: 'days'
  delta_t: '1'
  total_duration: '1500'
  origin: 'January 1'


grouping:
  population:


    HSVPG: [health_state, vaccinated, pen]
    # HSVP: [health_state, vaccinated, pen]
    BF: [BFR, health_state, batch]
    HSB: [health_state, batch]
    ISB: [inseminationStatus, batch]
    PS: [physiologicalStep, animal_type, batch, culling, litters]
    ATB: [animal_type, batch]
    CB: [culling, batch]
    ATBL: [animal_type, batch, litters]
    GS: [gestating_status, batch]
    PI: [production_impact]
    MI: [maternal_immunity]
    VP: [vaccinated, health_state, sector]
    HSS: [health_state, sector]
    C: [culling]

processes:
  individuals:
    - animal_type: ATBL
    - health_state: HSVPG
    - BFR: BF
    - inseminationStatus: ISB
    - culling: CB
    - physiologicalStep: PS
    - gestating_status: GS
    - production_impact: PI
    - maternal_immunity: MI
    - vaccinated: VP


levels:
  population:
    desc: 'level of the population'
    aggregation_type: 'hybrid'
    default_prototype: default_pop
    contains:
      - individuals
  individuals:
    desc: 'level of the individuals'
    default_prototype: default_pig

state_machines:
  animal_type:
    desc: 'type of animals'
    states:
      - Sow:
          name: 'Sow'
          desc: 'Sow'
          fillcolor: orange
      - Piglet:
          name: 'Piglet'
          desc: 'Piglet'
          fillcolor: blue
          on_stay:
            - set_var: weight
              value: weight_growth

  maternal_immunity:
    desc: 'Maternal immunity status for piglets'
    states:
      - M:
          name: 'M'
          desc: "with maternal immunity"
          fillcolor: green
          duration: mda_duration
      - notM:
          name: 'notM'
          desc: 'no maternal immunity'
          fillcolor: blue
    transitions:
      - from: M
        to: notM
        proba: 1

  vaccinated:
    desc: 'vaccination status'
    states:
      - vac:
          name: 'vaccinated'
          desc: 'with vaccination'
          fillcolor: green
      - notVac:
          name: 'notVaccinated'
          desc: 'without vaccination'
          fillcolor: blue

  health_state:
    states:
      - S:
          name: "Susceptible non-vaccinated"
          desc: "Susceptible non vaccinated"
          fillcolor: green
          on_enter:
            - become: sowST
              proba: '(total_ST < initial_I_sow) * (time < 2)'
      - E:
          name: "Exposed non-vaccinated"
          desc: 'Exposed non-vaccinated'
          fillcolor: orange
          duration: exposed_duration
      - I:
          name: "Infected non-vaccinated"
          desc: "Infected non-vaccinated"
          fillcolor: red
          on_enter:
            - become: [infected_first, infected_second, infected_third]
              proba: [first_gest, second_gest, third_gest]
      - R:
          name: "Recovered non-vaccinated"
          desc: "Recovered non vaccinated"
          fillcolor: blue
      - ST:
          name: "transitory Sow S, becoming In to simulate introduction after a delay"
          desc: 'transitory Sow'
          duration: delay_introduction
      - D:
          name: "Deceased"
          desc: "Deceased"
          autoremove: yes
    transitions:
      - from: S
        to: E
        rate: 'force_of_infection'
        cond: withE
      - from: S
        to: I
        rate: 'force_of_infection'
        cond: 'NOT(withE)'
      - from: E
        to: I
        proba: 1
      - from: I
        to: R
        rate: 'gamma'
      - from: ST
        to: I
        proba: 1

  BFR:
    desc: 'alternate BFR status'
    states:
      - BFR0:
          name: 'BFR0'
          desc: 'BFR 0 for initial conditions'
          fillcolor: black
      - BFR1:
          name: 'BFR1'
          desc: 'BFR 1'
          fillcolor: blue
      - BFR2:
          name: 'BFR2'
          desc: 'BFR 2'
          fillcolor: green
      - BFR3:
          name: 'BFR3'
          desc: 'BFR 3'
          fillcolor: red
          next: BFR1

  gestating_status:
    desc: 'gestating state of sows. Divided into 3 periods, each period
    corresponds to different levels of impact of the disease on the
    transmission of maternal immunity and abortions.'
    states:
      - first_gestating_period:
          name: 'first gestating period'
          desc: 'first gestating period: impact only on abortion. Produce S'
          duration: first_gestating_duration
          on_enter:
            - set_var: farrowing_rank
              value: "farrowing_rank + 1"
      - second_gestating_period:
          name: 'second gestating period'
          desc: 'second gestating period: impact on abortion. Produce I'
          duration: second_gestating_duration
      - third_gestating_period:
          name: 'third gestating period'
          desc: 'third gestting period: impact on abortion. Produce M'
          duration: third_gestating_duration
          on_exit:
            - produce_offspring: [newborn_S, newborn_I, newborn_M]
              amount: 'fertility'
              proba: [prod_S, prod_I, prod_M]
      - nonGestating:
          name: 'non-gestating'
          desc: 'non-gestating'
          default: yes
    transitions:
      - from: first_gestating_period
        to: second_gestating_period
        proba: 1
      - from: second_gestating_period
        to: third_gestating_period
        proba: 1
      - from: third_gestating_period
        to: nonGestating
        proba: 1

  production_impact:
    desc: 'status of offspring production for sows, depending on the period of infection during gestation'
    states:
      - produce_S:
          name: 'produce S'
          desc: "produce S"
          fillcolor: blue
          default: yes
      - produce_I:
          name: 'produce I'
          desc: "produce I"
          fillcolor: green
      - produce_M:
          name: 'produce M'
          desc: "produce M"
          fillcolor: red
      - no_production:
          name: 'no_production'
          desc: 'no_production - for piglets...'
          fillcolor: grey

  physiologicalStep:
    desc: 'physiological step evolution'
    states:
      - Quarantine:
          name: "Quarantine"
          desc: 'state used for initilize the population: each batch
          exit at batch_interval day interval'
          duration: decay_exit
      - Insemination:
          name: 'Insemination'
          desc: 'Insemination period (only for sows)'
          fillcolor: green
          duration: insemination_duration
          on_enter:
            - become: gestatingSow
              proba: is_Sow
      - Gestating:
          name: 'Gestating'
          desc: 'Gestating period (only for sows)'
          fillcolor: brown
          duration: gestating_duration
          on_enter:
            - become: inseminateSow
              proba: 1
      - Farrowing:
          name: 'Farrowing'
          desc: 'Farrowing period (for sows and piglets)'
          fillcolor: blue
          duration: farrowing_duration
          on_enter:
            - become: to_cull_proto
              proba: proba_to_cull
      - PostWeaning:
          name: 'PostWeaning'
          desc: 'Post-weaning period (only for piglets)'
          fillcolor: purple
          duration: post_weaning_duration
      - Finishing:
          name: 'finishing'
          desc: 'Finishing period (only for piglets)'
          fillcolor: purple
          line_style: dotted
          duration: finishing_duration
      - Exit:
          name: 'Exit'
          desc: 'Exit the system (slaughterhouse, dead, etc.)'
          fillcolor: black
          autoremove: True
    transitions:
      - from: Quarantine
        to: Insemination
        proba: 1
      - from: Insemination
        to: Gestating
        proba: 1
      - from: Gestating
        to: Farrowing
        proba: 1
        on_cross:
          - become: 'next_BFR'
      - from: Farrowing
        to: Insemination
        proba: 1
        cond: is_Sow
        on_cross:
          - produce_offspring: new_Sow
            proba: proba_produce
      - from: Farrowing
        to: PostWeaning
        proba: 1
        cond: is_Piglet
      - from: PostWeaning
        to: Finishing
        proba: 1
      - from: Finishing
        to: Exit
        proba: 1

  inseminationStatus:
    desc: 'Insemination status: failure or succes'
    states:
      - inseminate:
          name: 'inseminate'
          desc: 'inselinate'
          duration: '(2*batch_interval) - insemination_duration'
      - failureIns:
          name: 'failureIns'
          desc: 'failed insemination'
          on_enter:
              - become: [failure_insemination_batch1, failure_insemination_batch2, failure_insemination_batch3, failure_insemination_batch4, failure_insemination_batch5, failure_insemination_batch6, failure_insemination_batch7]
                proba: [proba_apply_protoype_batch1, proba_apply_protoype_batch2, proba_apply_protoype_batch3, proba_apply_protoype_batch4, proba_apply_protoype_batch5, proba_apply_protoype_batch6, proba_apply_protoype_batch7]
      - successIns:
          name: 'successIns'
          desc: 'successfull insemination'
    transitions:
      - from: inseminate
        to: failureIns
        proba: 'proba_fail_insemination * is_redoub'
      - from: inseminate
        to: successIns
        proba: 'IfThenElse(is_redoub, 1-proba_fail_insemination, 1)'

  culling:
    desc: 'process to cull and renew sows'
    states:
      - to_cull:
          name: 'to_cull'
          desc: 'sows that are coming to cull'
          duration: 'farrowing_duration'
      - to_keep:
          name: 'to_keep'
          desc: 'sows that not coming to cull'
      - died:
          name: 'died'
          desc: 'culling'
          autoremove: True
    transitions:
      - from: to_cull
        to: died
        proba: 1

organizations:
  litter:
    name: 'litter'
    desc: "litter organization (social aspect)"
    execute_process:
      - allocate
    spaces:
      name: 'litters'
      description: 'litters'
      allocation:
        - if: 'is_Piglet'
          then: 'with_source()'
        - if: 'is_Sow'
          then: alternate_free(spaces=ALL, filter=[Sow, batches])
      nodes:
        litter01:
          informations:
            excret:
              value: 0
        litter02:
          informations:
            excret:
              value: 0
        litter03:
          informations:
            excret:
              value: 0
        litter04:
          informations:
            excret:
              value: 0
        litter05:
          informations:
            excret:
              value: 0
        litter06:
          informations:
            excret:
              value: 0
        litter07:
          informations:
            excret:
              value: 0
        litter08:
          informations:
            excret:
              value: 0
        litter09:
          informations:
            excret:
              value: 0
        litter10:
          informations:
            excret:
              value: 0
        litter11:
          informations:
            excret:
              value: 0
        litter12:
          informations:
            excret:
              value: 0
        litter13:
          informations:
            excret:
              value: 0
        litter14:
          informations:
            excret:
              value: 0
        litter15:
          informations:
            excret:
              value: 0
        litter16:
          informations:
            excret:
              value: 0
        litter17:
          informations:
            excret:
              value: 0
        litter18:
          informations:
            excret:
              value: 0
  pen_group:
    name: 'pen_group'
    desc: "pen_group organization (social aspect)"
    execute_process:
      - allocate
    spaces:
      name: 'pen_groups'
      description: 'pen_groups'
      allocation:
        - if: 'isin_litter01 + isin_litter02'
          then: pen_group1
        - if: 'isin_litter03 + isin_litter04'
          then: pen_group2
        - if: 'isin_litter05 + isin_litter06'
          then: pen_group3
        - if: 'isin_litter07 + isin_litter08'
          then: pen_group4
        - if: 'isin_litter09 + isin_litter10'
          then: pen_group5
        - if: 'isin_litter11 + isin_litter12'
          then: pen_group6
        - if: 'isin_litter13 + isin_litter14'
          then: pen_group7
        - if: 'isin_litter15 + isin_litter16'
          then: pen_group8
        - if: 'isin_litter17 + isin_litter18'
          then: pen_group9
      nodes:
        pen_group0:
          informations:
            excret:
              value: 0
        pen_group1:
          informations:
            excret:
              value: 0
        pen_group2:
          informations:
            excret:
              value: 0
        pen_group3:
          informations:
            excret:
              value: 0
        pen_group4:
          informations:
            excret:
              value: 0
        pen_group5:
          informations:
            excret:
              value: 0
        pen_group6:
          informations:
            excret:
              value: 0
        pen_group7:
          informations:
            excret:
              value: 0
        pen_group8:
          informations:
            excret:
              value: 0
        pen_group9:
          informations:
            excret:
              value: 0
  batches:
    name: 'batches'
    desc: "batch management"
    execute_process:
      - allocate
    spaces:
      name: 'batch'
      description: 'batch'
      allocation:
        - if: 'is_Piglet + is_Sow'
          then: 'with_source()'
          else: 'batch7'
      nodes:
        batch1:
          index: 1
          informations:
            excret:
              value: 0
        batch2:
          index: 2
          informations:
            excret:
              value: 0
        batch3:
          index: 3
          informations:
            excret:
              value: 0
        batch4:
          index: 4
          informations:
            excret:
              value: 0
        batch5:
          index: 5
          informations:
            excret:
              value: 0
        batch6:
          index: 6
          informations:
            excret:
              value: 0
        batch7:
          index: 7
          informations:
            excret:
              value: 0
    state_machines:
      - animal_type


  housing:
    name: 'housing'
    description: 'Housing in sector (i.e. buildings)'
    execute_process:
      - allocate
    spaces:
      name: 'sector'
      description: 'sector (buildings) corresponding to physiological step'
      allocation:
        - if: 'is_Quarantine'
          then: quarantineSector
        - if: 'is_Insemination'
          then: serviceSector
        - if: 'is_Gestating'
          then: gestatingSector
        - if: 'is_Farrowing'
          then: farrowingSector
        - if: 'is_PostWeaning'
          then: nurserySector
        - if: 'is_Finishing'
          then: finishingSector
      nodes:
        quarantineSector:
          reference: quarantineSector
        serviceSector:
          reference: serviceSector
        gestatingSector:
          reference: gestatingSector
        farrowingSector:
          reference: farrowingSector
        nurserySector:
          reference: nurserySector
        finishingSector:
          reference: finishingSector
      graph:
        - 'serviceSector ->(excret:1) gestatingSector'
        - 'gestatingSector ->(excret:1) farrowingSector'
        - 'farrowingSector ->(excret:1) nurserySector'
        - 'nurserySector ->(excret:1) finishingSector'
    state_machines:
      - physiologicalStep

  quarantineSector:
    name: 'quarantineSector'
    desc: 'used to initialize the population (see the associated state_machine)'
    execute_process:
      - allocate
    spaces:
      name: 'room'
      description: 'room of the quarantine sector (only one)'
      informations:
        excret:
          value: 0
      allocation:
        - quarantine1
      nodes:
        quarantine1:
          reference: quarantine1

  quarantine1:
    name: "quarantine1"
    desc: 'pens division of room quarantine1'
    execute_process:
      - allocate
    spaces:
      name: 'pen'
      description: 'pens of the quarantine1 room'
      informations:
        excret:
          value: 2
      allocation:
        - quarantine1_pen1
      nodes:
        quarantine1_pen1:
          informations:
            excret:
              value: 0
  serviceSector:
    name: 'serviceSector'
    desc: 'service sector'
    execute_process:
      - allocate
    spaces:
      name: 'room'
      description: 'sector in single room'
      allocation:
        - service1
      nodes:
        service1:
          reference: service1

  service1:
    name: 'service1'
    desc: 'service1'
    execute_process:
      - allocate
    spaces:
      name: 'pen'
      description: 'only one room'
      allocation:
        - service1_pen1
      nodes:
        service1_pen1:
          informations:
            excret:
              value: 0


  gestatingSector:
    name: 'gestatingSector'
    desc: 'gestating sector'
    execute_process:
      - allocate
    spaces:
      name: 'room'
      descritpion: 'sector is divided in 4 rooms'
      allocation:
        - 'alternate_by_group(group=batch, spaces=ALL)'
      nodes:
        gestating1:
          reference: gestating1
        gestating2:
          reference: gestating2
        gestating3:
          reference: gestating3
        gestating4:
          reference: gestating4

  gestating1:
    name: 'gestating 1'
    desc: 'gestating 1'
    processes:
      deposit_pen1:
        action: 'set_to(information=excret, value=total_I_notVac_gestating1_pen1, spaces=[gestating1_pen1])'
      deposit_pen2:
        action: 'set_to(information=excret, value=total_I_notVac_gestating1_pen2, spaces=[gestating1_pen2])'
      deposit_pen3:
        action: 'set_to(information=excret, value=total_I_notVac_gestating1_pen3, spaces=[gestating1_pen3])'
    execute_process:
      - allocate
      - deposit_pen1
      - deposit_pen2
      - deposit_pen3
      - propagate_information
    spaces:
      name: 'pen'
      description: '3 pens -> groups of 6 sows'
      allocation:
        - if: 'isin_pen_group1 + isin_pen_group2 + isin_pen_group3'
          then: gestating1_pen1
        - if: 'isin_pen_group4 + isin_pen_group5 + isin_pen_group6'
          then: gestating1_pen2
        - if: 'isin_pen_group7 + isin_pen_group8 + isin_pen_group9'
          then: gestating1_pen3
      nodes:
        gestating1_pen1:
          informations:
            excret:
              value: 0


        gestating1_pen2:
          informations:
            excret:
              value: 0


        gestating1_pen3:
          informations:
            excret:
              value: 0
      graph:
        - "gestating1_pen1 ->(excret: propag_inter) gestating1_pen2 -> (excret: propag_inter) gestating1_pen1"
        - "gestating1_pen1 ->(excret: propag_inter) gestating1_pen3 -> (excret: propag_inter) gestating1_pen1"
        - "gestating1_pen2 ->(excret: propag_inter) gestating1_pen3 -> (excret: propag_inter) gestating1_pen2"

  gestating2:
    name: 'gestating 2'
    desc: 'gestating 2'
    processes:
      deposit_pen1:
        action: 'set_to(information=excret, value=total_I_notVac_gestating2_pen1, spaces=[gestating2_pen1])'
      deposit_pen2:
        action: 'set_to(information=excret, value=total_I_notVac_gestating2_pen2, spaces=[gestating2_pen2])'
      deposit_pen3:
        action: 'set_to(information=excret, value=total_I_notVac_gestating2_pen3, spaces=[gestating2_pen3])'
    execute_process:
      - allocate
      - deposit_pen1
      - deposit_pen2
      - deposit_pen3
      - propagate_information
    spaces:
      name: 'pen'
      description: '3 pens -> groups of 6 sows'
      allocation:
        - if: 'isin_pen_group1 + isin_pen_group2 + isin_pen_group3'
          then: gestating2_pen1
        - if: 'isin_pen_group4 + isin_pen_group5 + isin_pen_group6'
          then: gestating2_pen2
        - if: 'isin_pen_group7 + isin_pen_group8 + isin_pen_group9'
          then: gestating2_pen3
      nodes:
        gestating2_pen1:
          informations:
            excret:
              value: 0


        gestating2_pen2:
          informations:
            excret:
              value: 0


        gestating2_pen3:
          informations:
            excret:
              value: 0
      graph:
        - "gestating2_pen1 ->(excret: propag_inter) gestating2_pen2 -> (excret: propag_inter) gestating2_pen1"
        - "gestating2_pen1 ->(excret: propag_inter) gestating2_pen3 -> (excret: propag_inter) gestating2_pen1"
        - "gestating2_pen2 ->(excret: propag_inter) gestating2_pen3 -> (excret: propag_inter) gestating2_pen2"

  gestating3:
    name: 'gestating 3'
    desc: 'gestating 3'
    processes:
      deposit_pen1:
        action: 'set_to(information=excret, value=total_I_notVac_gestating3_pen1, spaces=[gestating3_pen1])'
      deposit_pen2:
        action: 'set_to(information=excret, value=total_I_notVac_gestating3_pen2, spaces=[gestating3_pen2])'
      deposit_pen3:
        action: 'set_to(information=excret, value=total_I_notVac_gestating3_pen3, spaces=[gestating3_pen3])'
    execute_process:
      - allocate
      - deposit_pen1
      - deposit_pen2
      - deposit_pen3
      - propagate_information
    spaces:
      name: 'pen'
      description: '15 pens'
      allocation:
        - if: 'isin_pen_group1 + isin_pen_group2 + isin_pen_group3'
          then: gestating3_pen1
        - if: 'isin_pen_group4 + isin_pen_group5 + isin_pen_group6'
          then: gestating3_pen2
        - if: 'isin_pen_group7 + isin_pen_group8 + isin_pen_group9'
          then: gestating3_pen3
      nodes:
        gestating3_pen1:
          informations:
            excret:
              value: 0


        gestating3_pen2:
          informations:
            excret:
              value: 0


        gestating3_pen3:
          informations:
            excret:
              value: 0
      graph:
        - "gestating3_pen1 ->(excret: propag_inter) gestating3_pen2 -> (excret: propag_inter) gestating3_pen1"
        - "gestating3_pen1 ->(excret: propag_inter) gestating3_pen3 -> (excret: propag_inter) gestating3_pen1"
        - "gestating3_pen2 ->(excret: propag_inter) gestating3_pen3 -> (excret: propag_inter) gestating3_pen2"
  gestating4:
    name: 'gestating 1'
    desc: 'gestating 1'
    processes:
      deposit_pen1:
        action: 'set_to(information=excret, value=total_I_notVac_gestating4_pen1, spaces=[gestating4_pen1])'
      deposit_pen2:
        action: 'set_to(information=excret, value=total_I_notVac_gestating4_pen2, spaces=[gestating4_pen2])'
      deposit_pen3:
        action: 'set_to(information=excret, value=total_I_notVac_gestating4_pen3, spaces=[gestating4_pen3])'
    execute_process:
      - allocate
      - deposit_pen1
      - deposit_pen2
      - deposit_pen3
      - propagate_information
    spaces:
      name: 'pen'
      description: '15 pens'
      allocation:
        - if: 'isin_pen_group1 + isin_pen_group2 + isin_pen_group3'
          then: gestating4_pen1
        - if: 'isin_pen_group4 + isin_pen_group5 + isin_pen_group6'
          then: gestating4_pen2
        - if: 'isin_pen_group7 + isin_pen_group8 + isin_pen_group9'
          then: gestating4_pen3
      nodes:
        gestating4_pen1:
          informations:
            excret:
              value: 0


        gestating4_pen2:
          informations:
            excret:
              value: 0


        gestating4_pen3:
          informations:
            excret:
              value: 0
      graph:
        - "gestating4_pen1 ->(excret: propag_inter) gestating4_pen2 -> (excret: propag_inter) gestating4_pen1"
        - "gestating4_pen1 ->(excret: propag_inter) gestating4_pen3 -> (excret: propag_inter) gestating4_pen1"
        - "gestating4_pen2 ->(excret: propag_inter) gestating4_pen3 -> (excret: propag_inter) gestating4_pen2"

  farrowingSector:
    name: 'farrowingeSector'
    desc: 'farrowing sector'
    execute_process:
      - allocate
    spaces:
      name: 'room'
      descritpion: 'sector is divided in 2 rooms'
      allocation:
        - if: 'is_Piglet'
          then: 'with_source()'
          else: 'alternate_by_group(group=batch, spaces=ALL)'
      nodes:
        farrowing1:
          reference: farrowing1
        farrowing2:
          reference: farrowing2


  farrowing1:
    name: 'farrowing 1'
    desc: 'farrowing 1'
    processes:
      deposit_pen1:
        action: 'set_to(information=excret, value=total_I_notVac_farrowing1_pen1, spaces=[farrowing1_pen1])'
      deposit_pen2:
        action: 'set_to(information=excret, value=total_I_notVac_farrowing1_pen2, spaces=[farrowing1_pen2])'
      deposit_pen3:
        action: 'set_to(information=excret, value=total_I_notVac_farrowing1_pen3, spaces=[farrowing1_pen3])'
      deposit_pen4:
        action: 'set_to(information=excret, value=total_I_notVac_farrowing1_pen4, spaces=[farrowing1_pen4])'
      deposit_pen5:
        action: 'set_to(information=excret, value=total_I_notVac_farrowing1_pen5, spaces=[farrowing1_pen5])'
      deposit_pen6:
        action: 'set_to(information=excret, value=total_I_notVac_farrowing1_pen6, spaces=[farrowing1_pen6])'
      deposit_pen7:
        action: 'set_to(information=excret, value=total_I_notVac_farrowing1_pen7, spaces=[farrowing1_pen7])'
      deposit_pen8:
        action: 'set_to(information=excret, value=total_I_notVac_farrowing1_pen8, spaces=[farrowing1_pen8])'
      deposit_pen9:
        action: 'set_to(information=excret, value=total_I_notVac_farrowing1_pen9, spaces=[farrowing1_pen9])'
      deposit_pen10:
        action: 'set_to(information=excret, value=total_I_notVac_farrowing1_pen10, spaces=[farrowing1_pen10])'
      deposit_pen11:
        action: 'set_to(information=excret, value=total_I_notVac_farrowing1_pen11, spaces=[farrowing1_pen11])'
      deposit_pen12:
        action: 'set_to(information=excret, value=total_I_notVac_farrowing1_pen12, spaces=[farrowing1_pen12])'
      deposit_pen13:
        action: 'set_to(information=excret, value=total_I_notVac_farrowing1_pen13, spaces=[farrowing1_pen13])'
      deposit_pen14:
        action: 'set_to(information=excret, value=total_I_notVac_farrowing1_pen14, spaces=[farrowing1_pen14])'
      deposit_pen15:
        action: 'set_to(information=excret, value=total_I_notVac_farrowing1_pen15, spaces=[farrowing1_pen15])'
      deposit_pen16:
        action: 'set_to(information=excret, value=total_I_notVac_farrowing1_pen16, spaces=[farrowing1_pen16])'
      deposit_pen17:
        action: 'set_to(information=excret, value=total_I_notVac_farrowing1_pen17, spaces=[farrowing1_pen17])'
      deposit_pen18:
        action: 'set_to(information=excret, value=total_I_notVac_farrowing1_pen18, spaces=[farrowing1_pen18])'
    execute_process:
      - allocate
      - deposit_pen1
      - deposit_pen2
      - deposit_pen3
      - deposit_pen4
      - deposit_pen5
      - deposit_pen6
      - deposit_pen7
      - deposit_pen8
      - deposit_pen9
      - deposit_pen10
      - deposit_pen11
      - deposit_pen12
      - deposit_pen13
      - deposit_pen14
      - deposit_pen15
      - deposit_pen16
      - deposit_pen17
      - deposit_pen18
      - propagate_information
    spaces:
      name: 'pen'
      description: '15 pens'
      allocation:
        - if: 'isin_litter01'
          then: farrowing1_pen1
        - if: 'isin_litter02'
          then: farrowing1_pen2
        - if: 'isin_litter03'
          then: farrowing1_pen3
        - if: 'isin_litter04'
          then: farrowing1_pen4
        - if: 'isin_litter05'
          then: farrowing1_pen5
        - if: 'isin_litter06'
          then: farrowing1_pen6
        - if: 'isin_litter07'
          then: farrowing1_pen7
        - if: 'isin_litter08'
          then: farrowing1_pen8
        - if: 'isin_litter09'
          then: farrowing1_pen9
        - if: 'isin_litter10'
          then: farrowing1_pen10
        - if: 'isin_litter11'
          then: farrowing1_pen11
        - if: 'isin_litter12'
          then: farrowing1_pen12
        - if: 'isin_litter13'
          then: farrowing1_pen13
        - if: 'isin_litter14'
          then: farrowing1_pen14
        - if: 'isin_litter15'
          then: farrowing1_pen15
        - if: 'isin_litter16'
          then: farrowing1_pen16
        - if: 'isin_litter17'
          then: farrowing1_pen17
        - if: 'isin_litter18'
          then: farrowing1_pen18
      nodes:
        farrowing1_pen1:
          informations:
            excret:
              value: 0


        farrowing1_pen2:
          informations:
            excret:
              value: 0


        farrowing1_pen3:
          informations:
            excret:
              value: 0


        farrowing1_pen4:
          informations:
            excret:
              value: 0


        farrowing1_pen5:
          informations:
            excret:
              value: 0


        farrowing1_pen6:
          informations:
            excret:
              value: 0


        farrowing1_pen7:
          informations:
            excret:
              value: 0


        farrowing1_pen8:
          informations:
            excret:
              value: 0


        farrowing1_pen9:
          informations:
            excret:
              value: 0


        farrowing1_pen10:
          informations:
            excret:
              value: 0


        farrowing1_pen11:
          informations:
            excret:
              value: 0


        farrowing1_pen12:
          informations:
            excret:
              value: 0


        farrowing1_pen13:
          informations:
            excret:
              value: 0


        farrowing1_pen14:
          informations:
            excret:
              value: 0


        farrowing1_pen15:
          informations:
            excret:
              value: 0


        farrowing1_pen16:
          informations:
            excret:
              value: 0


        farrowing1_pen17:
          informations:
            excret:
              value: 0


        farrowing1_pen18:
          informations:
            excret:
              value: 0
      graph:
        - 'farrowing1_pen1 ->(excret: propag_inter) farrowing1_pen2 ->(excret: propag_inter) farrowing1_pen1'
        - 'farrowing1_pen1 ->(excret: propag_inter) farrowing1_pen3 ->(excret: propag_inter) farrowing1_pen1'
        - 'farrowing1_pen1 ->(excret: propag_inter) farrowing1_pen4 ->(excret: propag_inter) farrowing1_pen1'
        - 'farrowing1_pen1 ->(excret: propag_inter) farrowing1_pen5 ->(excret: propag_inter) farrowing1_pen1'
        - 'farrowing1_pen1 ->(excret: propag_inter) farrowing1_pen6 ->(excret: propag_inter) farrowing1_pen1'
        - 'farrowing1_pen1 ->(excret: propag_inter) farrowing1_pen7 ->(excret: propag_inter) farrowing1_pen1'
        - 'farrowing1_pen1 ->(excret: propag_inter) farrowing1_pen8 ->(excret: propag_inter) farrowing1_pen1'
        - 'farrowing1_pen1 ->(excret: propag_inter) farrowing1_pen9 ->(excret: propag_inter) farrowing1_pen1'
        - 'farrowing1_pen1 ->(excret: propag_inter) farrowing1_pen10 ->(excret: propag_inter) farrowing1_pen1'
        - 'farrowing1_pen1 ->(excret: propag_inter) farrowing1_pen11 ->(excret: propag_inter) farrowing1_pen1'
        - 'farrowing1_pen1 ->(excret: propag_inter) farrowing1_pen12 ->(excret: propag_inter) farrowing1_pen1'
        - 'farrowing1_pen1 ->(excret: propag_inter) farrowing1_pen13 ->(excret: propag_inter) farrowing1_pen1'
        - 'farrowing1_pen1 ->(excret: propag_inter) farrowing1_pen14 ->(excret: propag_inter) farrowing1_pen1'
        - 'farrowing1_pen1 ->(excret: propag_inter) farrowing1_pen15 ->(excret: propag_inter) farrowing1_pen1'
        - 'farrowing1_pen1 ->(excret: propag_inter) farrowing1_pen16 ->(excret: propag_inter) farrowing1_pen1'
        - 'farrowing1_pen1 ->(excret: propag_inter) farrowing1_pen17 ->(excret: propag_inter) farrowing1_pen1'
        - 'farrowing1_pen1 ->(excret: propag_inter) farrowing1_pen18 ->(excret: propag_inter) farrowing1_pen1'
        - 'farrowing1_pen2->(excret: propag_inter) farrowing1_pen3 ->(excret: propag_inter) farrowing1_pen2'
        - 'farrowing1_pen2->(excret: propag_inter) farrowing1_pen4 ->(excret: propag_inter) farrowing1_pen2'
        - 'farrowing1_pen2->(excret: propag_inter) farrowing1_pen5 ->(excret: propag_inter) farrowing1_pen2'
        - 'farrowing1_pen2->(excret: propag_inter) farrowing1_pen6 ->(excret: propag_inter) farrowing1_pen2'
        - 'farrowing1_pen2->(excret: propag_inter) farrowing1_pen7 ->(excret: propag_inter) farrowing1_pen2'
        - 'farrowing1_pen2->(excret: propag_inter) farrowing1_pen8 ->(excret: propag_inter) farrowing1_pen2'
        - 'farrowing1_pen2->(excret: propag_inter) farrowing1_pen9 ->(excret: propag_inter) farrowing1_pen2'
        - 'farrowing1_pen2->(excret: propag_inter) farrowing1_pen10 ->(excret: propag_inter) farrowing1_pen2'
        - 'farrowing1_pen2->(excret: propag_inter) farrowing1_pen11 ->(excret: propag_inter) farrowing1_pen2'
        - 'farrowing1_pen2->(excret: propag_inter) farrowing1_pen12 ->(excret: propag_inter) farrowing1_pen2'
        - 'farrowing1_pen2->(excret: propag_inter) farrowing1_pen13 ->(excret: propag_inter) farrowing1_pen2'
        - 'farrowing1_pen2->(excret: propag_inter) farrowing1_pen14 ->(excret: propag_inter) farrowing1_pen2'
        - 'farrowing1_pen2->(excret: propag_inter) farrowing1_pen15 ->(excret: propag_inter) farrowing1_pen2'
        - 'farrowing1_pen2->(excret: propag_inter) farrowing1_pen16 ->(excret: propag_inter) farrowing1_pen2'
        - 'farrowing1_pen2->(excret: propag_inter) farrowing1_pen17 ->(excret: propag_inter) farrowing1_pen2'
        - 'farrowing1_pen2->(excret: propag_inter) farrowing1_pen18 ->(excret: propag_inter) farrowing1_pen2'
        - 'farrowing1_pen3->(excret: propag_inter) farrowing1_pen4 ->(excret: propag_inter) farrowing1_pen3'
        - 'farrowing1_pen3->(excret: propag_inter) farrowing1_pen5 ->(excret: propag_inter) farrowing1_pen3'
        - 'farrowing1_pen3->(excret: propag_inter) farrowing1_pen6 ->(excret: propag_inter) farrowing1_pen3'
        - 'farrowing1_pen3->(excret: propag_inter) farrowing1_pen7 ->(excret: propag_inter) farrowing1_pen3'
        - 'farrowing1_pen3->(excret: propag_inter) farrowing1_pen8 ->(excret: propag_inter) farrowing1_pen3'
        - 'farrowing1_pen3->(excret: propag_inter) farrowing1_pen9 ->(excret: propag_inter) farrowing1_pen3'
        - 'farrowing1_pen3->(excret: propag_inter) farrowing1_pen10 ->(excret: propag_inter) farrowing1_pen3'
        - 'farrowing1_pen3->(excret: propag_inter) farrowing1_pen11 ->(excret: propag_inter) farrowing1_pen3'
        - 'farrowing1_pen3->(excret: propag_inter) farrowing1_pen12 ->(excret: propag_inter) farrowing1_pen3'
        - 'farrowing1_pen3->(excret: propag_inter) farrowing1_pen13 ->(excret: propag_inter) farrowing1_pen3'
        - 'farrowing1_pen3->(excret: propag_inter) farrowing1_pen14 ->(excret: propag_inter) farrowing1_pen3'
        - 'farrowing1_pen3->(excret: propag_inter) farrowing1_pen15 ->(excret: propag_inter) farrowing1_pen3'
        - 'farrowing1_pen3->(excret: propag_inter) farrowing1_pen16 ->(excret: propag_inter) farrowing1_pen3'
        - 'farrowing1_pen3->(excret: propag_inter) farrowing1_pen17 ->(excret: propag_inter) farrowing1_pen3'
        - 'farrowing1_pen3->(excret: propag_inter) farrowing1_pen18 ->(excret: propag_inter) farrowing1_pen3'
        - 'farrowing1_pen4->(excret: propag_inter) farrowing1_pen5 ->(excret: propag_inter) farrowing1_pen4'
        - 'farrowing1_pen4->(excret: propag_inter) farrowing1_pen6 ->(excret: propag_inter) farrowing1_pen4'
        - 'farrowing1_pen4->(excret: propag_inter) farrowing1_pen7 ->(excret: propag_inter) farrowing1_pen4'
        - 'farrowing1_pen4->(excret: propag_inter) farrowing1_pen8 ->(excret: propag_inter) farrowing1_pen4'
        - 'farrowing1_pen4->(excret: propag_inter) farrowing1_pen9 ->(excret: propag_inter) farrowing1_pen4'
        - 'farrowing1_pen4->(excret: propag_inter) farrowing1_pen10 ->(excret: propag_inter) farrowing1_pen4'
        - 'farrowing1_pen4->(excret: propag_inter) farrowing1_pen11 ->(excret: propag_inter) farrowing1_pen4'
        - 'farrowing1_pen4->(excret: propag_inter) farrowing1_pen12 ->(excret: propag_inter) farrowing1_pen4'
        - 'farrowing1_pen4->(excret: propag_inter) farrowing1_pen13 ->(excret: propag_inter) farrowing1_pen4'
        - 'farrowing1_pen4->(excret: propag_inter) farrowing1_pen14 ->(excret: propag_inter) farrowing1_pen4'
        - 'farrowing1_pen4->(excret: propag_inter) farrowing1_pen15 ->(excret: propag_inter) farrowing1_pen4'
        - 'farrowing1_pen4->(excret: propag_inter) farrowing1_pen16 ->(excret: propag_inter) farrowing1_pen4'
        - 'farrowing1_pen4->(excret: propag_inter) farrowing1_pen17 ->(excret: propag_inter) farrowing1_pen4'
        - 'farrowing1_pen4->(excret: propag_inter) farrowing1_pen18 ->(excret: propag_inter) farrowing1_pen4'
        - 'farrowing1_pen5->(excret: propag_inter) farrowing1_pen6 ->(excret: propag_inter) farrowing1_pen5'
        - 'farrowing1_pen5->(excret: propag_inter) farrowing1_pen7 ->(excret: propag_inter) farrowing1_pen5'
        - 'farrowing1_pen5->(excret: propag_inter) farrowing1_pen8 ->(excret: propag_inter) farrowing1_pen5'
        - 'farrowing1_pen5->(excret: propag_inter) farrowing1_pen9 ->(excret: propag_inter) farrowing1_pen5'
        - 'farrowing1_pen5->(excret: propag_inter) farrowing1_pen10 ->(excret: propag_inter) farrowing1_pen5'
        - 'farrowing1_pen5->(excret: propag_inter) farrowing1_pen11 ->(excret: propag_inter) farrowing1_pen5'
        - 'farrowing1_pen5->(excret: propag_inter) farrowing1_pen12 ->(excret: propag_inter) farrowing1_pen5'
        - 'farrowing1_pen5->(excret: propag_inter) farrowing1_pen13 ->(excret: propag_inter) farrowing1_pen5'
        - 'farrowing1_pen5->(excret: propag_inter) farrowing1_pen14 ->(excret: propag_inter) farrowing1_pen5'
        - 'farrowing1_pen5->(excret: propag_inter) farrowing1_pen15 ->(excret: propag_inter) farrowing1_pen5'
        - 'farrowing1_pen5->(excret: propag_inter) farrowing1_pen16 ->(excret: propag_inter) farrowing1_pen5'
        - 'farrowing1_pen5->(excret: propag_inter) farrowing1_pen17 ->(excret: propag_inter) farrowing1_pen5'
        - 'farrowing1_pen5->(excret: propag_inter) farrowing1_pen18 ->(excret: propag_inter) farrowing1_pen5'
        - 'farrowing1_pen6->(excret: propag_inter) farrowing1_pen7 ->(excret: propag_inter) farrowing1_pen6'
        - 'farrowing1_pen6->(excret: propag_inter) farrowing1_pen8 ->(excret: propag_inter) farrowing1_pen6'
        - 'farrowing1_pen6->(excret: propag_inter) farrowing1_pen9 ->(excret: propag_inter) farrowing1_pen6'
        - 'farrowing1_pen6->(excret: propag_inter) farrowing1_pen10 ->(excret: propag_inter) farrowing1_pen6'
        - 'farrowing1_pen6->(excret: propag_inter) farrowing1_pen11 ->(excret: propag_inter) farrowing1_pen6'
        - 'farrowing1_pen6->(excret: propag_inter) farrowing1_pen12 ->(excret: propag_inter) farrowing1_pen6'
        - 'farrowing1_pen6->(excret: propag_inter) farrowing1_pen13 ->(excret: propag_inter) farrowing1_pen6'
        - 'farrowing1_pen6->(excret: propag_inter) farrowing1_pen14 ->(excret: propag_inter) farrowing1_pen6'
        - 'farrowing1_pen6->(excret: propag_inter) farrowing1_pen15 ->(excret: propag_inter) farrowing1_pen6'
        - 'farrowing1_pen6->(excret: propag_inter) farrowing1_pen16 ->(excret: propag_inter) farrowing1_pen6'
        - 'farrowing1_pen6->(excret: propag_inter) farrowing1_pen17 ->(excret: propag_inter) farrowing1_pen6'
        - 'farrowing1_pen6->(excret: propag_inter) farrowing1_pen18 ->(excret: propag_inter) farrowing1_pen6'
        - 'farrowing1_pen7->(excret: propag_inter) farrowing1_pen8 ->(excret: propag_inter) farrowing1_pen7'
        - 'farrowing1_pen7->(excret: propag_inter) farrowing1_pen9 ->(excret: propag_inter) farrowing1_pen7'
        - 'farrowing1_pen7->(excret: propag_inter) farrowing1_pen10 ->(excret: propag_inter) farrowing1_pen7'
        - 'farrowing1_pen7->(excret: propag_inter) farrowing1_pen11 ->(excret: propag_inter) farrowing1_pen7'
        - 'farrowing1_pen7->(excret: propag_inter) farrowing1_pen12 ->(excret: propag_inter) farrowing1_pen7'
        - 'farrowing1_pen7->(excret: propag_inter) farrowing1_pen13 ->(excret: propag_inter) farrowing1_pen7'
        - 'farrowing1_pen7->(excret: propag_inter) farrowing1_pen14 ->(excret: propag_inter) farrowing1_pen7'
        - 'farrowing1_pen7->(excret: propag_inter) farrowing1_pen15 ->(excret: propag_inter) farrowing1_pen7'
        - 'farrowing1_pen7->(excret: propag_inter) farrowing1_pen16 ->(excret: propag_inter) farrowing1_pen7'
        - 'farrowing1_pen7->(excret: propag_inter) farrowing1_pen17 ->(excret: propag_inter) farrowing1_pen7'
        - 'farrowing1_pen7->(excret: propag_inter) farrowing1_pen18 ->(excret: propag_inter) farrowing1_pen7'
        - 'farrowing1_pen8->(excret: propag_inter) farrowing1_pen9 ->(excret: propag_inter) farrowing1_pen8'
        - 'farrowing1_pen8->(excret: propag_inter) farrowing1_pen10 ->(excret: propag_inter) farrowing1_pen8'
        - 'farrowing1_pen8->(excret: propag_inter) farrowing1_pen11 ->(excret: propag_inter) farrowing1_pen8'
        - 'farrowing1_pen8->(excret: propag_inter) farrowing1_pen12 ->(excret: propag_inter) farrowing1_pen8'
        - 'farrowing1_pen8->(excret: propag_inter) farrowing1_pen13 ->(excret: propag_inter) farrowing1_pen8'
        - 'farrowing1_pen8->(excret: propag_inter) farrowing1_pen14 ->(excret: propag_inter) farrowing1_pen8'
        - 'farrowing1_pen8->(excret: propag_inter) farrowing1_pen15 ->(excret: propag_inter) farrowing1_pen8'
        - 'farrowing1_pen8->(excret: propag_inter) farrowing1_pen16 ->(excret: propag_inter) farrowing1_pen8'
        - 'farrowing1_pen8->(excret: propag_inter) farrowing1_pen17 ->(excret: propag_inter) farrowing1_pen8'
        - 'farrowing1_pen8->(excret: propag_inter) farrowing1_pen18 ->(excret: propag_inter) farrowing1_pen8'
        - 'farrowing1_pen9->(excret: propag_inter) farrowing1_pen10 ->(excret: propag_inter) farrowing1_pen9'
        - 'farrowing1_pen9->(excret: propag_inter) farrowing1_pen11 ->(excret: propag_inter) farrowing1_pen9'
        - 'farrowing1_pen9->(excret: propag_inter) farrowing1_pen12 ->(excret: propag_inter) farrowing1_pen9'
        - 'farrowing1_pen9->(excret: propag_inter) farrowing1_pen13 ->(excret: propag_inter) farrowing1_pen9'
        - 'farrowing1_pen9->(excret: propag_inter) farrowing1_pen14 ->(excret: propag_inter) farrowing1_pen9'
        - 'farrowing1_pen9->(excret: propag_inter) farrowing1_pen15 ->(excret: propag_inter) farrowing1_pen9'
        - 'farrowing1_pen9->(excret: propag_inter) farrowing1_pen16 ->(excret: propag_inter) farrowing1_pen9'
        - 'farrowing1_pen9->(excret: propag_inter) farrowing1_pen17 ->(excret: propag_inter) farrowing1_pen9'
        - 'farrowing1_pen9->(excret: propag_inter) farrowing1_pen18 ->(excret: propag_inter) farrowing1_pen9'
        - 'farrowing1_pen10->(excret: propag_inter) farrowing1_pen11 ->(excret: propag_inter) farrowing1_pen10'
        - 'farrowing1_pen10->(excret: propag_inter) farrowing1_pen12 ->(excret: propag_inter) farrowing1_pen10'
        - 'farrowing1_pen10->(excret: propag_inter) farrowing1_pen13 ->(excret: propag_inter) farrowing1_pen10'
        - 'farrowing1_pen10->(excret: propag_inter) farrowing1_pen14 ->(excret: propag_inter) farrowing1_pen10'
        - 'farrowing1_pen10->(excret: propag_inter) farrowing1_pen15 ->(excret: propag_inter) farrowing1_pen10'
        - 'farrowing1_pen10->(excret: propag_inter) farrowing1_pen16 ->(excret: propag_inter) farrowing1_pen10'
        - 'farrowing1_pen10->(excret: propag_inter) farrowing1_pen17 ->(excret: propag_inter) farrowing1_pen10'
        - 'farrowing1_pen10->(excret: propag_inter) farrowing1_pen18 ->(excret: propag_inter) farrowing1_pen10'
        - 'farrowing1_pen11->(excret: propag_inter) farrowing1_pen12 ->(excret: propag_inter) farrowing1_pen11'
        - 'farrowing1_pen11->(excret: propag_inter) farrowing1_pen13 ->(excret: propag_inter) farrowing1_pen11'
        - 'farrowing1_pen11->(excret: propag_inter) farrowing1_pen14 ->(excret: propag_inter) farrowing1_pen11'
        - 'farrowing1_pen11->(excret: propag_inter) farrowing1_pen15 ->(excret: propag_inter) farrowing1_pen11'
        - 'farrowing1_pen11->(excret: propag_inter) farrowing1_pen16 ->(excret: propag_inter) farrowing1_pen11'
        - 'farrowing1_pen11->(excret: propag_inter) farrowing1_pen17 ->(excret: propag_inter) farrowing1_pen11'
        - 'farrowing1_pen11->(excret: propag_inter) farrowing1_pen18 ->(excret: propag_inter) farrowing1_pen11'
        - 'farrowing1_pen12->(excret: propag_inter) farrowing1_pen13 ->(excret: propag_inter) farrowing1_pen12'
        - 'farrowing1_pen12->(excret: propag_inter) farrowing1_pen14 ->(excret: propag_inter) farrowing1_pen12'
        - 'farrowing1_pen12->(excret: propag_inter) farrowing1_pen15 ->(excret: propag_inter) farrowing1_pen12'
        - 'farrowing1_pen12->(excret: propag_inter) farrowing1_pen16 ->(excret: propag_inter) farrowing1_pen12'
        - 'farrowing1_pen12->(excret: propag_inter) farrowing1_pen17 ->(excret: propag_inter) farrowing1_pen12'
        - 'farrowing1_pen12->(excret: propag_inter) farrowing1_pen18 ->(excret: propag_inter) farrowing1_pen12'
        - 'farrowing1_pen13->(excret: propag_inter) farrowing1_pen14 ->(excret: propag_inter) farrowing1_pen13'
        - 'farrowing1_pen13->(excret: propag_inter) farrowing1_pen15 ->(excret: propag_inter) farrowing1_pen13'
        - 'farrowing1_pen13->(excret: propag_inter) farrowing1_pen16 ->(excret: propag_inter) farrowing1_pen13'
        - 'farrowing1_pen13->(excret: propag_inter) farrowing1_pen17 ->(excret: propag_inter) farrowing1_pen13'
        - 'farrowing1_pen13->(excret: propag_inter) farrowing1_pen18 ->(excret: propag_inter) farrowing1_pen13'
        - 'farrowing1_pen14->(excret: propag_inter) farrowing1_pen15 ->(excret: propag_inter) farrowing1_pen14'
        - 'farrowing1_pen14->(excret: propag_inter) farrowing1_pen16 ->(excret: propag_inter) farrowing1_pen14'
        - 'farrowing1_pen14->(excret: propag_inter) farrowing1_pen17 ->(excret: propag_inter) farrowing1_pen14'
        - 'farrowing1_pen14->(excret: propag_inter) farrowing1_pen18 ->(excret: propag_inter) farrowing1_pen14'
        - 'farrowing1_pen15->(excret: propag_inter) farrowing1_pen16 ->(excret: propag_inter) farrowing1_pen15'
        - 'farrowing1_pen15->(excret: propag_inter) farrowing1_pen17 ->(excret: propag_inter) farrowing1_pen15'
        - 'farrowing1_pen15->(excret: propag_inter) farrowing1_pen18 ->(excret: propag_inter) farrowing1_pen15'
        - 'farrowing1_pen16->(excret: propag_inter) farrowing1_pen17 ->(excret: propag_inter) farrowing1_pen16'
        - 'farrowing1_pen16->(excret: propag_inter) farrowing1_pen18 ->(excret: propag_inter) farrowing1_pen16'
        - 'farrowing1_pen17->(excret: propag_inter) farrowing1_pen18 ->(excret: propag_inter) farrowing1_pen17'


  farrowing2:
    name: 'farrowing 1'
    desc: 'farrowing 1'
    processes:
      deposit_pen1:
        action: 'set_to(information=excret, value=total_I_notVac_farrowing2_pen1, spaces=[farrowing2_pen1])'
      deposit_pen2:
        action: 'set_to(information=excret, value=total_I_notVac_farrowing2_pen2, spaces=[farrowing2_pen2])'
      deposit_pen3:
        action: 'set_to(information=excret, value=total_I_notVac_farrowing2_pen3, spaces=[farrowing2_pen3])'
      deposit_pen4:
        action: 'set_to(information=excret, value=total_I_notVac_farrowing2_pen4, spaces=[farrowing2_pen4])'
      deposit_pen5:
        action: 'set_to(information=excret, value=total_I_notVac_farrowing2_pen5, spaces=[farrowing2_pen5])'
      deposit_pen6:
        action: 'set_to(information=excret, value=total_I_notVac_farrowing2_pen6, spaces=[farrowing2_pen6])'
      deposit_pen7:
        action: 'set_to(information=excret, value=total_I_notVac_farrowing2_pen7, spaces=[farrowing2_pen7])'
      deposit_pen8:
        action: 'set_to(information=excret, value=total_I_notVac_farrowing2_pen8, spaces=[farrowing2_pen8])'
      deposit_pen9:
        action: 'set_to(information=excret, value=total_I_notVac_farrowing2_pen9, spaces=[farrowing2_pen9])'
      deposit_pen10:
        action: 'set_to(information=excret, value=total_I_notVac_farrowing2_pen10, spaces=[farrowing2_pen10])'
      deposit_pen11:
        action: 'set_to(information=excret, value=total_I_notVac_farrowing2_pen11, spaces=[farrowing2_pen11])'
      deposit_pen12:
        action: 'set_to(information=excret, value=total_I_notVac_farrowing2_pen12, spaces=[farrowing2_pen12])'
      deposit_pen13:
        action: 'set_to(information=excret, value=total_I_notVac_farrowing2_pen13, spaces=[farrowing2_pen13])'
      deposit_pen14:
        action: 'set_to(information=excret, value=total_I_notVac_farrowing2_pen14, spaces=[farrowing2_pen14])'
      deposit_pen15:
        action: 'set_to(information=excret, value=total_I_notVac_farrowing2_pen15, spaces=[farrowing2_pen15])'
      deposit_pen16:
        action: 'set_to(information=excret, value=total_I_notVac_farrowing2_pen16, spaces=[farrowing2_pen16])'
      deposit_pen17:
        action: 'set_to(information=excret, value=total_I_notVac_farrowing2_pen17, spaces=[farrowing2_pen17])'
      deposit_pen18:
        action: 'set_to(information=excret, value=total_I_notVac_farrowing2_pen18, spaces=[farrowing2_pen18])'
    execute_process:
      - allocate
      - deposit_pen1
      - deposit_pen2
      - deposit_pen3
      - deposit_pen4
      - deposit_pen5
      - deposit_pen6
      - deposit_pen7
      - deposit_pen8
      - deposit_pen9
      - deposit_pen10
      - deposit_pen11
      - deposit_pen12
      - deposit_pen13
      - deposit_pen14
      - deposit_pen15
      - deposit_pen16
      - deposit_pen17
      - deposit_pen18
      - propagate_information
    spaces:
      name: 'pen'
      description: '15 pens'
      allocation:
        - if: 'isin_litter01'
          then: farrowing2_pen1
        - if: 'isin_litter02'
          then: farrowing2_pen2
        - if: 'isin_litter03'
          then: farrowing2_pen3
        - if: 'isin_litter04'
          then: farrowing2_pen4
        - if: 'isin_litter05'
          then: farrowing2_pen5
        - if: 'isin_litter06'
          then: farrowing2_pen6
        - if: 'isin_litter07'
          then: farrowing2_pen7
        - if: 'isin_litter08'
          then: farrowing2_pen8
        - if: 'isin_litter09'
          then: farrowing2_pen9
        - if: 'isin_litter10'
          then: farrowing2_pen10
        - if: 'isin_litter11'
          then: farrowing2_pen11
        - if: 'isin_litter12'
          then: farrowing2_pen12
        - if: 'isin_litter13'
          then: farrowing2_pen13
        - if: 'isin_litter14'
          then: farrowing2_pen14
        - if: 'isin_litter15'
          then: farrowing2_pen15
        - if: 'isin_litter16'
          then: farrowing2_pen16
        - if: 'isin_litter17'
          then: farrowing2_pen17
        - if: 'isin_litter18'
          then: farrowing2_pen18
      nodes:
        farrowing2_pen1:
          informations:
            excret:
              value: 0


        farrowing2_pen2:
          informations:
            excret:
              value: 0


        farrowing2_pen3:
          informations:
            excret:
              value: 0


        farrowing2_pen4:
          informations:
            excret:
              value: 0


        farrowing2_pen5:
          informations:
            excret:
              value: 0


        farrowing2_pen6:
          informations:
            excret:
              value: 0


        farrowing2_pen7:
          informations:
            excret:
              value: 0


        farrowing2_pen8:
          informations:
            excret:
              value: 0


        farrowing2_pen9:
          informations:
            excret:
              value: 0


        farrowing2_pen10:
          informations:
            excret:
              value: 0


        farrowing2_pen11:
          informations:
            excret:
              value: 0


        farrowing2_pen12:
          informations:
            excret:
              value: 0


        farrowing2_pen13:
          informations:
            excret:
              value: 0


        farrowing2_pen14:
          informations:
            excret:
              value: 0


        farrowing2_pen15:
          informations:
            excret:
              value: 0


        farrowing2_pen16:
          informations:
            excret:
              value: 0


        farrowing2_pen17:
          informations:
            excret:
              value: 0


        farrowing2_pen18:
          informations:
            excret:
              value: 0
      graph:
        - 'farrowing2_pen1 ->(excret: propag_inter) farrowing2_pen2 ->(excret: propag_inter) farrowing2_pen1'
        - 'farrowing2_pen1 ->(excret: propag_inter) farrowing2_pen3 ->(excret: propag_inter) farrowing2_pen1'
        - 'farrowing2_pen1 ->(excret: propag_inter) farrowing2_pen4 ->(excret: propag_inter) farrowing2_pen1'
        - 'farrowing2_pen1 ->(excret: propag_inter) farrowing2_pen5 ->(excret: propag_inter) farrowing2_pen1'
        - 'farrowing2_pen1 ->(excret: propag_inter) farrowing2_pen6 ->(excret: propag_inter) farrowing2_pen1'
        - 'farrowing2_pen1 ->(excret: propag_inter) farrowing2_pen7 ->(excret: propag_inter) farrowing2_pen1'
        - 'farrowing2_pen1 ->(excret: propag_inter) farrowing2_pen8 ->(excret: propag_inter) farrowing2_pen1'
        - 'farrowing2_pen1 ->(excret: propag_inter) farrowing2_pen9 ->(excret: propag_inter) farrowing2_pen1'
        - 'farrowing2_pen1 ->(excret: propag_inter) farrowing2_pen10 ->(excret: propag_inter) farrowing2_pen1'
        - 'farrowing2_pen1 ->(excret: propag_inter) farrowing2_pen11 ->(excret: propag_inter) farrowing2_pen1'
        - 'farrowing2_pen1 ->(excret: propag_inter) farrowing2_pen12 ->(excret: propag_inter) farrowing2_pen1'
        - 'farrowing2_pen1 ->(excret: propag_inter) farrowing2_pen13 ->(excret: propag_inter) farrowing2_pen1'
        - 'farrowing2_pen1 ->(excret: propag_inter) farrowing2_pen14 ->(excret: propag_inter) farrowing2_pen1'
        - 'farrowing2_pen1 ->(excret: propag_inter) farrowing2_pen15 ->(excret: propag_inter) farrowing2_pen1'
        - 'farrowing2_pen1 ->(excret: propag_inter) farrowing2_pen16 ->(excret: propag_inter) farrowing2_pen1'
        - 'farrowing2_pen1 ->(excret: propag_inter) farrowing2_pen17 ->(excret: propag_inter) farrowing2_pen1'
        - 'farrowing2_pen1 ->(excret: propag_inter) farrowing2_pen18 ->(excret: propag_inter) farrowing2_pen1'
        - 'farrowing2_pen2->(excret: propag_inter) farrowing2_pen3 ->(excret: propag_inter) farrowing2_pen2'
        - 'farrowing2_pen2->(excret: propag_inter) farrowing2_pen4 ->(excret: propag_inter) farrowing2_pen2'
        - 'farrowing2_pen2->(excret: propag_inter) farrowing2_pen5 ->(excret: propag_inter) farrowing2_pen2'
        - 'farrowing2_pen2->(excret: propag_inter) farrowing2_pen6 ->(excret: propag_inter) farrowing2_pen2'
        - 'farrowing2_pen2->(excret: propag_inter) farrowing2_pen7 ->(excret: propag_inter) farrowing2_pen2'
        - 'farrowing2_pen2->(excret: propag_inter) farrowing2_pen8 ->(excret: propag_inter) farrowing2_pen2'
        - 'farrowing2_pen2->(excret: propag_inter) farrowing2_pen9 ->(excret: propag_inter) farrowing2_pen2'
        - 'farrowing2_pen2->(excret: propag_inter) farrowing2_pen10 ->(excret: propag_inter) farrowing2_pen2'
        - 'farrowing2_pen2->(excret: propag_inter) farrowing2_pen11 ->(excret: propag_inter) farrowing2_pen2'
        - 'farrowing2_pen2->(excret: propag_inter) farrowing2_pen12 ->(excret: propag_inter) farrowing2_pen2'
        - 'farrowing2_pen2->(excret: propag_inter) farrowing2_pen13 ->(excret: propag_inter) farrowing2_pen2'
        - 'farrowing2_pen2->(excret: propag_inter) farrowing2_pen14 ->(excret: propag_inter) farrowing2_pen2'
        - 'farrowing2_pen2->(excret: propag_inter) farrowing2_pen15 ->(excret: propag_inter) farrowing2_pen2'
        - 'farrowing2_pen2->(excret: propag_inter) farrowing2_pen16 ->(excret: propag_inter) farrowing2_pen2'
        - 'farrowing2_pen2->(excret: propag_inter) farrowing2_pen17 ->(excret: propag_inter) farrowing2_pen2'
        - 'farrowing2_pen2->(excret: propag_inter) farrowing2_pen18 ->(excret: propag_inter) farrowing2_pen2'
        - 'farrowing2_pen3->(excret: propag_inter) farrowing2_pen4 ->(excret: propag_inter) farrowing2_pen3'
        - 'farrowing2_pen3->(excret: propag_inter) farrowing2_pen5 ->(excret: propag_inter) farrowing2_pen3'
        - 'farrowing2_pen3->(excret: propag_inter) farrowing2_pen6 ->(excret: propag_inter) farrowing2_pen3'
        - 'farrowing2_pen3->(excret: propag_inter) farrowing2_pen7 ->(excret: propag_inter) farrowing2_pen3'
        - 'farrowing2_pen3->(excret: propag_inter) farrowing2_pen8 ->(excret: propag_inter) farrowing2_pen3'
        - 'farrowing2_pen3->(excret: propag_inter) farrowing2_pen9 ->(excret: propag_inter) farrowing2_pen3'
        - 'farrowing2_pen3->(excret: propag_inter) farrowing2_pen10 ->(excret: propag_inter) farrowing2_pen3'
        - 'farrowing2_pen3->(excret: propag_inter) farrowing2_pen11 ->(excret: propag_inter) farrowing2_pen3'
        - 'farrowing2_pen3->(excret: propag_inter) farrowing2_pen12 ->(excret: propag_inter) farrowing2_pen3'
        - 'farrowing2_pen3->(excret: propag_inter) farrowing2_pen13 ->(excret: propag_inter) farrowing2_pen3'
        - 'farrowing2_pen3->(excret: propag_inter) farrowing2_pen14 ->(excret: propag_inter) farrowing2_pen3'
        - 'farrowing2_pen3->(excret: propag_inter) farrowing2_pen15 ->(excret: propag_inter) farrowing2_pen3'
        - 'farrowing2_pen3->(excret: propag_inter) farrowing2_pen16 ->(excret: propag_inter) farrowing2_pen3'
        - 'farrowing2_pen3->(excret: propag_inter) farrowing2_pen17 ->(excret: propag_inter) farrowing2_pen3'
        - 'farrowing2_pen3->(excret: propag_inter) farrowing2_pen18 ->(excret: propag_inter) farrowing2_pen3'
        - 'farrowing2_pen4->(excret: propag_inter) farrowing2_pen5 ->(excret: propag_inter) farrowing2_pen4'
        - 'farrowing2_pen4->(excret: propag_inter) farrowing2_pen6 ->(excret: propag_inter) farrowing2_pen4'
        - 'farrowing2_pen4->(excret: propag_inter) farrowing2_pen7 ->(excret: propag_inter) farrowing2_pen4'
        - 'farrowing2_pen4->(excret: propag_inter) farrowing2_pen8 ->(excret: propag_inter) farrowing2_pen4'
        - 'farrowing2_pen4->(excret: propag_inter) farrowing2_pen9 ->(excret: propag_inter) farrowing2_pen4'
        - 'farrowing2_pen4->(excret: propag_inter) farrowing2_pen10 ->(excret: propag_inter) farrowing2_pen4'
        - 'farrowing2_pen4->(excret: propag_inter) farrowing2_pen11 ->(excret: propag_inter) farrowing2_pen4'
        - 'farrowing2_pen4->(excret: propag_inter) farrowing2_pen12 ->(excret: propag_inter) farrowing2_pen4'
        - 'farrowing2_pen4->(excret: propag_inter) farrowing2_pen13 ->(excret: propag_inter) farrowing2_pen4'
        - 'farrowing2_pen4->(excret: propag_inter) farrowing2_pen14 ->(excret: propag_inter) farrowing2_pen4'
        - 'farrowing2_pen4->(excret: propag_inter) farrowing2_pen15 ->(excret: propag_inter) farrowing2_pen4'
        - 'farrowing2_pen4->(excret: propag_inter) farrowing2_pen16 ->(excret: propag_inter) farrowing2_pen4'
        - 'farrowing2_pen4->(excret: propag_inter) farrowing2_pen17 ->(excret: propag_inter) farrowing2_pen4'
        - 'farrowing2_pen4->(excret: propag_inter) farrowing2_pen18 ->(excret: propag_inter) farrowing2_pen4'
        - 'farrowing2_pen5->(excret: propag_inter) farrowing2_pen6 ->(excret: propag_inter) farrowing2_pen5'
        - 'farrowing2_pen5->(excret: propag_inter) farrowing2_pen7 ->(excret: propag_inter) farrowing2_pen5'
        - 'farrowing2_pen5->(excret: propag_inter) farrowing2_pen8 ->(excret: propag_inter) farrowing2_pen5'
        - 'farrowing2_pen5->(excret: propag_inter) farrowing2_pen9 ->(excret: propag_inter) farrowing2_pen5'
        - 'farrowing2_pen5->(excret: propag_inter) farrowing2_pen10 ->(excret: propag_inter) farrowing2_pen5'
        - 'farrowing2_pen5->(excret: propag_inter) farrowing2_pen11 ->(excret: propag_inter) farrowing2_pen5'
        - 'farrowing2_pen5->(excret: propag_inter) farrowing2_pen12 ->(excret: propag_inter) farrowing2_pen5'
        - 'farrowing2_pen5->(excret: propag_inter) farrowing2_pen13 ->(excret: propag_inter) farrowing2_pen5'
        - 'farrowing2_pen5->(excret: propag_inter) farrowing2_pen14 ->(excret: propag_inter) farrowing2_pen5'
        - 'farrowing2_pen5->(excret: propag_inter) farrowing2_pen15 ->(excret: propag_inter) farrowing2_pen5'
        - 'farrowing2_pen5->(excret: propag_inter) farrowing2_pen16 ->(excret: propag_inter) farrowing2_pen5'
        - 'farrowing2_pen5->(excret: propag_inter) farrowing2_pen17 ->(excret: propag_inter) farrowing2_pen5'
        - 'farrowing2_pen5->(excret: propag_inter) farrowing2_pen18 ->(excret: propag_inter) farrowing2_pen5'
        - 'farrowing2_pen6->(excret: propag_inter) farrowing2_pen7 ->(excret: propag_inter) farrowing2_pen6'
        - 'farrowing2_pen6->(excret: propag_inter) farrowing2_pen8 ->(excret: propag_inter) farrowing2_pen6'
        - 'farrowing2_pen6->(excret: propag_inter) farrowing2_pen9 ->(excret: propag_inter) farrowing2_pen6'
        - 'farrowing2_pen6->(excret: propag_inter) farrowing2_pen10 ->(excret: propag_inter) farrowing2_pen6'
        - 'farrowing2_pen6->(excret: propag_inter) farrowing2_pen11 ->(excret: propag_inter) farrowing2_pen6'
        - 'farrowing2_pen6->(excret: propag_inter) farrowing2_pen12 ->(excret: propag_inter) farrowing2_pen6'
        - 'farrowing2_pen6->(excret: propag_inter) farrowing2_pen13 ->(excret: propag_inter) farrowing2_pen6'
        - 'farrowing2_pen6->(excret: propag_inter) farrowing2_pen14 ->(excret: propag_inter) farrowing2_pen6'
        - 'farrowing2_pen6->(excret: propag_inter) farrowing2_pen15 ->(excret: propag_inter) farrowing2_pen6'
        - 'farrowing2_pen6->(excret: propag_inter) farrowing2_pen16 ->(excret: propag_inter) farrowing2_pen6'
        - 'farrowing2_pen6->(excret: propag_inter) farrowing2_pen17 ->(excret: propag_inter) farrowing2_pen6'
        - 'farrowing2_pen6->(excret: propag_inter) farrowing2_pen18 ->(excret: propag_inter) farrowing2_pen6'
        - 'farrowing2_pen7->(excret: propag_inter) farrowing2_pen8 ->(excret: propag_inter) farrowing2_pen7'
        - 'farrowing2_pen7->(excret: propag_inter) farrowing2_pen9 ->(excret: propag_inter) farrowing2_pen7'
        - 'farrowing2_pen7->(excret: propag_inter) farrowing2_pen10 ->(excret: propag_inter) farrowing2_pen7'
        - 'farrowing2_pen7->(excret: propag_inter) farrowing2_pen11 ->(excret: propag_inter) farrowing2_pen7'
        - 'farrowing2_pen7->(excret: propag_inter) farrowing2_pen12 ->(excret: propag_inter) farrowing2_pen7'
        - 'farrowing2_pen7->(excret: propag_inter) farrowing2_pen13 ->(excret: propag_inter) farrowing2_pen7'
        - 'farrowing2_pen7->(excret: propag_inter) farrowing2_pen14 ->(excret: propag_inter) farrowing2_pen7'
        - 'farrowing2_pen7->(excret: propag_inter) farrowing2_pen15 ->(excret: propag_inter) farrowing2_pen7'
        - 'farrowing2_pen7->(excret: propag_inter) farrowing2_pen16 ->(excret: propag_inter) farrowing2_pen7'
        - 'farrowing2_pen7->(excret: propag_inter) farrowing2_pen17 ->(excret: propag_inter) farrowing2_pen7'
        - 'farrowing2_pen7->(excret: propag_inter) farrowing2_pen18 ->(excret: propag_inter) farrowing2_pen7'
        - 'farrowing2_pen8->(excret: propag_inter) farrowing2_pen9 ->(excret: propag_inter) farrowing2_pen8'
        - 'farrowing2_pen8->(excret: propag_inter) farrowing2_pen10 ->(excret: propag_inter) farrowing2_pen8'
        - 'farrowing2_pen8->(excret: propag_inter) farrowing2_pen11 ->(excret: propag_inter) farrowing2_pen8'
        - 'farrowing2_pen8->(excret: propag_inter) farrowing2_pen12 ->(excret: propag_inter) farrowing2_pen8'
        - 'farrowing2_pen8->(excret: propag_inter) farrowing2_pen13 ->(excret: propag_inter) farrowing2_pen8'
        - 'farrowing2_pen8->(excret: propag_inter) farrowing2_pen14 ->(excret: propag_inter) farrowing2_pen8'
        - 'farrowing2_pen8->(excret: propag_inter) farrowing2_pen15 ->(excret: propag_inter) farrowing2_pen8'
        - 'farrowing2_pen8->(excret: propag_inter) farrowing2_pen16 ->(excret: propag_inter) farrowing2_pen8'
        - 'farrowing2_pen8->(excret: propag_inter) farrowing2_pen17 ->(excret: propag_inter) farrowing2_pen8'
        - 'farrowing2_pen8->(excret: propag_inter) farrowing2_pen18 ->(excret: propag_inter) farrowing2_pen8'
        - 'farrowing2_pen9->(excret: propag_inter) farrowing2_pen10 ->(excret: propag_inter) farrowing2_pen9'
        - 'farrowing2_pen9->(excret: propag_inter) farrowing2_pen11 ->(excret: propag_inter) farrowing2_pen9'
        - 'farrowing2_pen9->(excret: propag_inter) farrowing2_pen12 ->(excret: propag_inter) farrowing2_pen9'
        - 'farrowing2_pen9->(excret: propag_inter) farrowing2_pen13 ->(excret: propag_inter) farrowing2_pen9'
        - 'farrowing2_pen9->(excret: propag_inter) farrowing2_pen14 ->(excret: propag_inter) farrowing2_pen9'
        - 'farrowing2_pen9->(excret: propag_inter) farrowing2_pen15 ->(excret: propag_inter) farrowing2_pen9'
        - 'farrowing2_pen9->(excret: propag_inter) farrowing2_pen16 ->(excret: propag_inter) farrowing2_pen9'
        - 'farrowing2_pen9->(excret: propag_inter) farrowing2_pen17 ->(excret: propag_inter) farrowing2_pen9'
        - 'farrowing2_pen9->(excret: propag_inter) farrowing2_pen18 ->(excret: propag_inter) farrowing2_pen9'
        - 'farrowing2_pen10->(excret: propag_inter) farrowing2_pen11 ->(excret: propag_inter) farrowing2_pen10'
        - 'farrowing2_pen10->(excret: propag_inter) farrowing2_pen12 ->(excret: propag_inter) farrowing2_pen10'
        - 'farrowing2_pen10->(excret: propag_inter) farrowing2_pen13 ->(excret: propag_inter) farrowing2_pen10'
        - 'farrowing2_pen10->(excret: propag_inter) farrowing2_pen14 ->(excret: propag_inter) farrowing2_pen10'
        - 'farrowing2_pen10->(excret: propag_inter) farrowing2_pen15 ->(excret: propag_inter) farrowing2_pen10'
        - 'farrowing2_pen10->(excret: propag_inter) farrowing2_pen16 ->(excret: propag_inter) farrowing2_pen10'
        - 'farrowing2_pen10->(excret: propag_inter) farrowing2_pen17 ->(excret: propag_inter) farrowing2_pen10'
        - 'farrowing2_pen10->(excret: propag_inter) farrowing2_pen18 ->(excret: propag_inter) farrowing2_pen10'
        - 'farrowing2_pen11->(excret: propag_inter) farrowing2_pen12 ->(excret: propag_inter) farrowing2_pen11'
        - 'farrowing2_pen11->(excret: propag_inter) farrowing2_pen13 ->(excret: propag_inter) farrowing2_pen11'
        - 'farrowing2_pen11->(excret: propag_inter) farrowing2_pen14 ->(excret: propag_inter) farrowing2_pen11'
        - 'farrowing2_pen11->(excret: propag_inter) farrowing2_pen15 ->(excret: propag_inter) farrowing2_pen11'
        - 'farrowing2_pen11->(excret: propag_inter) farrowing2_pen16 ->(excret: propag_inter) farrowing2_pen11'
        - 'farrowing2_pen11->(excret: propag_inter) farrowing2_pen17 ->(excret: propag_inter) farrowing2_pen11'
        - 'farrowing2_pen11->(excret: propag_inter) farrowing2_pen18 ->(excret: propag_inter) farrowing2_pen11'
        - 'farrowing2_pen12->(excret: propag_inter) farrowing2_pen13 ->(excret: propag_inter) farrowing2_pen12'
        - 'farrowing2_pen12->(excret: propag_inter) farrowing2_pen14 ->(excret: propag_inter) farrowing2_pen12'
        - 'farrowing2_pen12->(excret: propag_inter) farrowing2_pen15 ->(excret: propag_inter) farrowing2_pen12'
        - 'farrowing2_pen12->(excret: propag_inter) farrowing2_pen16 ->(excret: propag_inter) farrowing2_pen12'
        - 'farrowing2_pen12->(excret: propag_inter) farrowing2_pen17 ->(excret: propag_inter) farrowing2_pen12'
        - 'farrowing2_pen12->(excret: propag_inter) farrowing2_pen18 ->(excret: propag_inter) farrowing2_pen12'
        - 'farrowing2_pen13->(excret: propag_inter) farrowing2_pen14 ->(excret: propag_inter) farrowing2_pen13'
        - 'farrowing2_pen13->(excret: propag_inter) farrowing2_pen15 ->(excret: propag_inter) farrowing2_pen13'
        - 'farrowing2_pen13->(excret: propag_inter) farrowing2_pen16 ->(excret: propag_inter) farrowing2_pen13'
        - 'farrowing2_pen13->(excret: propag_inter) farrowing2_pen17 ->(excret: propag_inter) farrowing2_pen13'
        - 'farrowing2_pen13->(excret: propag_inter) farrowing2_pen18 ->(excret: propag_inter) farrowing2_pen13'
        - 'farrowing2_pen14->(excret: propag_inter) farrowing2_pen15 ->(excret: propag_inter) farrowing2_pen14'
        - 'farrowing2_pen14->(excret: propag_inter) farrowing2_pen16 ->(excret: propag_inter) farrowing2_pen14'
        - 'farrowing2_pen14->(excret: propag_inter) farrowing2_pen17 ->(excret: propag_inter) farrowing2_pen14'
        - 'farrowing2_pen14->(excret: propag_inter) farrowing2_pen18 ->(excret: propag_inter) farrowing2_pen14'
        - 'farrowing2_pen15->(excret: propag_inter) farrowing2_pen16 ->(excret: propag_inter) farrowing2_pen15'
        - 'farrowing2_pen15->(excret: propag_inter) farrowing2_pen17 ->(excret: propag_inter) farrowing2_pen15'
        - 'farrowing2_pen15->(excret: propag_inter) farrowing2_pen18 ->(excret: propag_inter) farrowing2_pen15'
        - 'farrowing2_pen16->(excret: propag_inter) farrowing2_pen17 ->(excret: propag_inter) farrowing2_pen16'
        - 'farrowing2_pen16->(excret: propag_inter) farrowing2_pen18 ->(excret: propag_inter) farrowing2_pen16'
        - 'farrowing2_pen17->(excret: propag_inter) farrowing2_pen18 ->(excret: propag_inter) farrowing2_pen17'

  nurserySector:
    name: 'nurserySector'
    desc: 'nursery sector'
    execute_process:
      - allocate
    spaces:
      name: 'room'
      descritpion: 'sector is divided in 4 rooms'
      allocation:
        - 'alternate_by_group(group=batch, spaces=ALL)'
      nodes:
        nursery1:
          reference: nursery1
        nursery2:
          reference: nursery2
        nursery3:
          reference: nursery3


  nursery1:
    name: 'nursery 1'
    desc: 'nursery 1'
    processes:
      deposit_pen1:
        action: 'set_to(information=excret, value=total_I_notVac_nursery1_pen1, spaces=[nursery1_pen1])'
      deposit_pen2:
        action: 'set_to(information=excret, value=total_I_notVac_nursery1_pen2, spaces=[nursery1_pen2])'
      deposit_pen3:
        action: 'set_to(information=excret, value=total_I_notVac_nursery1_pen3, spaces=[nursery1_pen3])'
      deposit_pen4:
        action: 'set_to(information=excret, value=total_I_notVac_nursery1_pen4, spaces=[nursery1_pen4])'
      deposit_pen5:
        action: 'set_to(information=excret, value=total_I_notVac_nursery1_pen5, spaces=[nursery1_pen5])'
      deposit_pen6:
        action: 'set_to(information=excret, value=total_I_notVac_nursery1_pen6, spaces=[nursery1_pen6])'
      deposit_pen7:
        action: 'set_to(information=excret, value=total_I_notVac_nursery1_pen7, spaces=[nursery1_pen7])'
      deposit_pen8:
        action: 'set_to(information=excret, value=total_I_notVac_nursery1_pen8, spaces=[nursery1_pen8])'
    execute_process:
      - allocate
      - deposit_pen1
      - deposit_pen2
      - deposit_pen3
      - deposit_pen4
      - deposit_pen5
      - deposit_pen6
      - deposit_pen7
      - deposit_pen8
      - propagate_information
    spaces:
      name: 'pen'
      description: '15 pens'
      allocation:
        - if: 'isin_pen_group1'
          then: nursery1_pen1
        - if: 'isin_pen_group2'
          then: nursery1_pen2
        - if: 'isin_pen_group3'
          then: nursery1_pen3
        - if: 'isin_pen_group4'
          then: nursery1_pen4
        - if: 'isin_pen_group5'
          then: nursery1_pen5
        - if: 'isin_pen_group6'
          then: nursery1_pen6
        - if: 'isin_pen_group7'
          then: nursery1_pen7
        - if: 'isin_pen_group8'
          then: nursery1_pen8
        - if: 'isin_pen_group9'
          then: nursery1_pen9
      nodes:
        nursery1_pen1:
          informations:
            excret:
              value: 0


        nursery1_pen2:
          informations:
            excret:
              value: 0


        nursery1_pen3:
          informations:
            excret:
              value: 0


        nursery1_pen4:
          informations:
            excret:
              value: 0


        nursery1_pen5:
          informations:
            excret:
              value: 0


        nursery1_pen6:
          informations:
            excret:
              value: 0


        nursery1_pen7:
          informations:
            excret:
              value: 0


        nursery1_pen8:
          informations:
            excret:
              value: 0


        nursery1_pen9:
          informations:
            excret:
              value: 0
      graph:
        - "nursery1_pen1 ->(excret: propag_intra) nursery1_pen2 -> (excret: propag_intra) nursery1_pen1"
        - "nursery1_pen3 ->(excret: propag_intra) nursery1_pen4 -> (excret: propag_intra) nursery1_pen3"
        - "nursery1_pen5 ->(excret: propag_intra) nursery1_pen6 -> (excret: propag_intra) nursery1_pen5"
        - "nursery1_pen7 ->(excret: propag_intra) nursery1_pen8 -> (excret: propag_intra) nursery1_pen7"
        - "nursery1_pen1 ->(excret: propag_inter) nursery1_pen3 -> (excret: propag_inter) nursery1_pen1"
        - "nursery1_pen1 ->(excret: propag_inter) nursery1_pen4 -> (excret: propag_inter) nursery1_pen1"
        - "nursery1_pen1 ->(excret: propag_inter) nursery1_pen5 -> (excret: propag_inter) nursery1_pen1"
        - "nursery1_pen1 ->(excret: propag_inter) nursery1_pen6 -> (excret: propag_inter) nursery1_pen1"
        - "nursery1_pen1 ->(excret: propag_inter) nursery1_pen7 -> (excret: propag_inter) nursery1_pen1"
        - "nursery1_pen1 ->(excret: propag_inter) nursery1_pen8 -> (excret: propag_inter) nursery1_pen1"
        - "nursery1_pen1 ->(excret: propag_inter) nursery1_pen9 -> (excret: propag_inter) nursery1_pen1"
        - "nursery1_pen2 ->(excret: propag_inter) nursery1_pen3 -> (excret: propag_inter) nursery1_pen2"
        - "nursery1_pen2 ->(excret: propag_inter) nursery1_pen4 -> (excret: propag_inter) nursery1_pen2"
        - "nursery1_pen2 ->(excret: propag_inter) nursery1_pen5 -> (excret: propag_inter) nursery1_pen2"
        - "nursery1_pen2 ->(excret: propag_inter) nursery1_pen6 -> (excret: propag_inter) nursery1_pen2"
        - "nursery1_pen2 ->(excret: propag_inter) nursery1_pen7 -> (excret: propag_inter) nursery1_pen2"
        - "nursery1_pen2 ->(excret: propag_inter) nursery1_pen8 -> (excret: propag_inter) nursery1_pen2"
        - "nursery1_pen2 ->(excret: propag_inter) nursery1_pen9 -> (excret: propag_inter) nursery1_pen2"
        - "nursery1_pen3 ->(excret: propag_inter) nursery1_pen5 -> (excret: propag_inter) nursery1_pen3"
        - "nursery1_pen3 ->(excret: propag_inter) nursery1_pen6 -> (excret: propag_inter) nursery1_pen3"
        - "nursery1_pen3 ->(excret: propag_inter) nursery1_pen7 -> (excret: propag_inter) nursery1_pen3"
        - "nursery1_pen3 ->(excret: propag_inter) nursery1_pen8 -> (excret: propag_inter) nursery1_pen3"
        - "nursery1_pen3 ->(excret: propag_inter) nursery1_pen9 -> (excret: propag_inter) nursery1_pen3"
        - "nursery1_pen4 ->(excret: propag_inter) nursery1_pen5 -> (excret: propag_inter) nursery1_pen4"
        - "nursery1_pen4 ->(excret: propag_inter) nursery1_pen6 -> (excret: propag_inter) nursery1_pen4"
        - "nursery1_pen4 ->(excret: propag_inter) nursery1_pen7 -> (excret: propag_inter) nursery1_pen4"
        - "nursery1_pen4 ->(excret: propag_inter) nursery1_pen8 -> (excret: propag_inter) nursery1_pen4"
        - "nursery1_pen4 ->(excret: propag_inter) nursery1_pen9 -> (excret: propag_inter) nursery1_pen4"
        - "nursery1_pen5 ->(excret: propag_inter) nursery1_pen7 -> (excret: propag_inter) nursery1_pen5"
        - "nursery1_pen5 ->(excret: propag_inter) nursery1_pen8 -> (excret: propag_inter) nursery1_pen5"
        - "nursery1_pen5 ->(excret: propag_inter) nursery1_pen9 -> (excret: propag_inter) nursery1_pen5"
        - "nursery1_pen6 ->(excret: propag_inter) nursery1_pen7 -> (excret: propag_inter) nursery1_pen6"
        - "nursery1_pen6 ->(excret: propag_inter) nursery1_pen8 -> (excret: propag_inter) nursery1_pen6"
        - "nursery1_pen6 ->(excret: propag_inter) nursery1_pen9 -> (excret: propag_inter) nursery1_pen6"
        - "nursery1_pen7 ->(excret: propag_inter) nursery1_pen9 -> (excret: propag_inter) nursery1_pen7"
        - "nursery1_pen8 ->(excret: propag_inter) nursery1_pen9 -> (excret: propag_inter) nursery1_pen8"

  nursery2:
    name: 'nursery 2'
    desc: 'nursery 2'
    processes:
      deposit_pen1:
        action: 'set_to(information=excret, value=total_I_notVac_nursery2_pen1, spaces=[nursery2_pen1])'
      deposit_pen2:
        action: 'set_to(information=excret, value=total_I_notVac_nursery2_pen2, spaces=[nursery2_pen2])'
      deposit_pen3:
        action: 'set_to(information=excret, value=total_I_notVac_nursery2_pen3, spaces=[nursery2_pen3])'
      deposit_pen4:
        action: 'set_to(information=excret, value=total_I_notVac_nursery2_pen4, spaces=[nursery2_pen4])'
      deposit_pen5:
        action: 'set_to(information=excret, value=total_I_notVac_nursery2_pen5, spaces=[nursery2_pen5])'
      deposit_pen6:
        action: 'set_to(information=excret, value=total_I_notVac_nursery2_pen6, spaces=[nursery2_pen6])'
      deposit_pen7:
        action: 'set_to(information=excret, value=total_I_notVac_nursery2_pen7, spaces=[nursery2_pen7])'
      deposit_pen8:
        action: 'set_to(information=excret, value=total_I_notVac_nursery2_pen8, spaces=[nursery2_pen8])'
    execute_process:
      - allocate
      - deposit_pen1
      - deposit_pen2
      - deposit_pen3
      - deposit_pen4
      - deposit_pen5
      - deposit_pen6
      - deposit_pen7
      - deposit_pen8
      - propagate_information
    spaces:
      name: 'pen'
      description: '15 pens'
      allocation:
        - if: 'isin_pen_group1'
          then: nursery2_pen1
        - if: 'isin_pen_group2'
          then: nursery2_pen2
        - if: 'isin_pen_group3'
          then: nursery2_pen3
        - if: 'isin_pen_group4'
          then: nursery2_pen4
        - if: 'isin_pen_group5'
          then: nursery2_pen5
        - if: 'isin_pen_group6'
          then: nursery2_pen6
        - if: 'isin_pen_group7'
          then: nursery2_pen7
        - if: 'isin_pen_group8'
          then: nursery2_pen8
        - if: 'isin_pen_group9'
          then: nursery2_pen9
      nodes:
        nursery2_pen1:
          informations:
            excret:
              value: 0


        nursery2_pen2:
          informations:
            excret:
              value: 0


        nursery2_pen3:
          informations:
            excret:
              value: 0


        nursery2_pen4:
          informations:
            excret:
              value: 0


        nursery2_pen5:
          informations:
            excret:
              value: 0


        nursery2_pen6:
          informations:
            excret:
              value: 0


        nursery2_pen7:
          informations:
            excret:
              value: 0


        nursery2_pen8:
          informations:
            excret:
              value: 0


        nursery2_pen9:
          informations:
            excret:
              value: 0
      graph:
        - "nursery2_pen1 ->(excret: propag_intra) nursery2_pen2 -> (excret: propag_intra) nursery2_pen1"
        - "nursery2_pen3 ->(excret: propag_intra) nursery2_pen4 -> (excret: propag_intra) nursery2_pen3"
        - "nursery2_pen5 ->(excret: propag_intra) nursery2_pen6 -> (excret: propag_intra) nursery2_pen5"
        - "nursery2_pen7 ->(excret: propag_intra) nursery2_pen8 -> (excret: propag_intra) nursery2_pen7"
        - "nursery2_pen1 ->(excret: propag_inter) nursery2_pen3 -> (excret: propag_inter) nursery2_pen1"
        - "nursery2_pen1 ->(excret: propag_inter) nursery2_pen4 -> (excret: propag_inter) nursery2_pen1"
        - "nursery2_pen1 ->(excret: propag_inter) nursery2_pen5 -> (excret: propag_inter) nursery2_pen1"
        - "nursery2_pen1 ->(excret: propag_inter) nursery2_pen6 -> (excret: propag_inter) nursery2_pen1"
        - "nursery2_pen1 ->(excret: propag_inter) nursery2_pen7 -> (excret: propag_inter) nursery2_pen1"
        - "nursery2_pen1 ->(excret: propag_inter) nursery2_pen8 -> (excret: propag_inter) nursery2_pen1"
        - "nursery2_pen1 ->(excret: propag_inter) nursery2_pen9 -> (excret: propag_inter) nursery2_pen1"
        - "nursery2_pen2 ->(excret: propag_inter) nursery2_pen3 -> (excret: propag_inter) nursery2_pen2"
        - "nursery2_pen2 ->(excret: propag_inter) nursery2_pen4 -> (excret: propag_inter) nursery2_pen2"
        - "nursery2_pen2 ->(excret: propag_inter) nursery2_pen5 -> (excret: propag_inter) nursery2_pen2"
        - "nursery2_pen2 ->(excret: propag_inter) nursery2_pen6 -> (excret: propag_inter) nursery2_pen2"
        - "nursery2_pen2 ->(excret: propag_inter) nursery2_pen7 -> (excret: propag_inter) nursery2_pen2"
        - "nursery2_pen2 ->(excret: propag_inter) nursery2_pen8 -> (excret: propag_inter) nursery2_pen2"
        - "nursery2_pen2 ->(excret: propag_inter) nursery2_pen9 -> (excret: propag_inter) nursery2_pen2"
        - "nursery2_pen3 ->(excret: propag_inter) nursery2_pen5 -> (excret: propag_inter) nursery2_pen3"
        - "nursery2_pen3 ->(excret: propag_inter) nursery2_pen6 -> (excret: propag_inter) nursery2_pen3"
        - "nursery2_pen3 ->(excret: propag_inter) nursery2_pen7 -> (excret: propag_inter) nursery2_pen3"
        - "nursery2_pen3 ->(excret: propag_inter) nursery2_pen8 -> (excret: propag_inter) nursery2_pen3"
        - "nursery2_pen3 ->(excret: propag_inter) nursery2_pen9 -> (excret: propag_inter) nursery2_pen3"
        - "nursery2_pen4 ->(excret: propag_inter) nursery2_pen5 -> (excret: propag_inter) nursery2_pen4"
        - "nursery2_pen4 ->(excret: propag_inter) nursery2_pen6 -> (excret: propag_inter) nursery2_pen4"
        - "nursery2_pen4 ->(excret: propag_inter) nursery2_pen7 -> (excret: propag_inter) nursery2_pen4"
        - "nursery2_pen4 ->(excret: propag_inter) nursery2_pen8 -> (excret: propag_inter) nursery2_pen4"
        - "nursery2_pen4 ->(excret: propag_inter) nursery2_pen9 -> (excret: propag_inter) nursery2_pen4"
        - "nursery2_pen5 ->(excret: propag_inter) nursery2_pen7 -> (excret: propag_inter) nursery2_pen5"
        - "nursery2_pen5 ->(excret: propag_inter) nursery2_pen8 -> (excret: propag_inter) nursery2_pen5"
        - "nursery2_pen5 ->(excret: propag_inter) nursery2_pen9 -> (excret: propag_inter) nursery2_pen5"
        - "nursery2_pen6 ->(excret: propag_inter) nursery2_pen7 -> (excret: propag_inter) nursery2_pen6"
        - "nursery2_pen6 ->(excret: propag_inter) nursery2_pen8 -> (excret: propag_inter) nursery2_pen6"
        - "nursery2_pen6 ->(excret: propag_inter) nursery2_pen9 -> (excret: propag_inter) nursery2_pen6"
        - "nursery2_pen7 ->(excret: propag_inter) nursery2_pen9 -> (excret: propag_inter) nursery2_pen7"
        - "nursery2_pen8 ->(excret: propag_inter) nursery2_pen9 -> (excret: propag_inter) nursery2_pen8"

  nursery3:
    name: 'nursery 3'
    desc: 'nursery 3'
    processes:
      deposit_pen1:
        action: 'set_to(information=excret, value=total_I_notVac_nursery3_pen1, spaces=[nursery3_pen1])'
      deposit_pen2:
        action: 'set_to(information=excret, value=total_I_notVac_nursery3_pen2, spaces=[nursery3_pen2])'
      deposit_pen3:
        action: 'set_to(information=excret, value=total_I_notVac_nursery3_pen3, spaces=[nursery3_pen3])'
      deposit_pen4:
        action: 'set_to(information=excret, value=total_I_notVac_nursery3_pen4, spaces=[nursery3_pen4])'
      deposit_pen5:
        action: 'set_to(information=excret, value=total_I_notVac_nursery3_pen5, spaces=[nursery3_pen5])'
      deposit_pen6:
        action: 'set_to(information=excret, value=total_I_notVac_nursery3_pen6, spaces=[nursery3_pen6])'
      deposit_pen7:
        action: 'set_to(information=excret, value=total_I_notVac_nursery3_pen7, spaces=[nursery3_pen7])'
      deposit_pen8:
        action: 'set_to(information=excret, value=total_I_notVac_nursery3_pen8, spaces=[nursery3_pen8])'
    execute_process:
      - allocate
      - deposit_pen1
      - deposit_pen2
      - deposit_pen3
      - deposit_pen4
      - deposit_pen5
      - deposit_pen6
      - deposit_pen7
      - deposit_pen8
      - propagate_information
    spaces:
      name: 'pen'
      description: '15 pens'
      allocation:
        - if: 'isin_pen_group1'
          then: nursery3_pen1
        - if: 'isin_pen_group2'
          then: nursery3_pen2
        - if: 'isin_pen_group3'
          then: nursery3_pen3
        - if: 'isin_pen_group4'
          then: nursery3_pen4
        - if: 'isin_pen_group5'
          then: nursery3_pen5
        - if: 'isin_pen_group6'
          then: nursery3_pen6
        - if: 'isin_pen_group7'
          then: nursery3_pen7
        - if: 'isin_pen_group8'
          then: nursery3_pen8
        - if: 'isin_pen_group9'
          then: nursery3_pen9
      nodes:
        nursery3_pen1:
          informations:
            excret:
              value: 0


        nursery3_pen2:
          informations:
            excret:
              value: 0


        nursery3_pen3:
          informations:
            excret:
              value: 0


        nursery3_pen4:
          informations:
            excret:
              value: 0


        nursery3_pen5:
          informations:
            excret:
              value: 0


        nursery3_pen6:
          informations:
            excret:
              value: 0


        nursery3_pen7:
          informations:
            excret:
              value: 0


        nursery3_pen8:
          informations:
            excret:
              value: 0


        nursery3_pen9:
          informations:
            excret:
              value: 0
      graph:
        - "nursery3_pen1 ->(excret: propag_intra) nursery3_pen2 -> (excret: propag_intra) nursery3_pen1"
        - "nursery3_pen3 ->(excret: propag_intra) nursery3_pen4 -> (excret: propag_intra) nursery3_pen3"
        - "nursery3_pen5 ->(excret: propag_intra) nursery3_pen6 -> (excret: propag_intra) nursery3_pen5"
        - "nursery3_pen7 ->(excret: propag_intra) nursery3_pen8 -> (excret: propag_intra) nursery3_pen7"
        - "nursery3_pen1 ->(excret: propag_inter) nursery3_pen3 -> (excret: propag_inter) nursery3_pen1"
        - "nursery3_pen1 ->(excret: propag_inter) nursery3_pen4 -> (excret: propag_inter) nursery3_pen1"
        - "nursery3_pen1 ->(excret: propag_inter) nursery3_pen5 -> (excret: propag_inter) nursery3_pen1"
        - "nursery3_pen1 ->(excret: propag_inter) nursery3_pen6 -> (excret: propag_inter) nursery3_pen1"
        - "nursery3_pen1 ->(excret: propag_inter) nursery3_pen7 -> (excret: propag_inter) nursery3_pen1"
        - "nursery3_pen1 ->(excret: propag_inter) nursery3_pen8 -> (excret: propag_inter) nursery3_pen1"
        - "nursery3_pen1 ->(excret: propag_inter) nursery3_pen9 -> (excret: propag_inter) nursery3_pen1"
        - "nursery3_pen2 ->(excret: propag_inter) nursery3_pen3 -> (excret: propag_inter) nursery3_pen2"
        - "nursery3_pen2 ->(excret: propag_inter) nursery3_pen4 -> (excret: propag_inter) nursery3_pen2"
        - "nursery3_pen2 ->(excret: propag_inter) nursery3_pen5 -> (excret: propag_inter) nursery3_pen2"
        - "nursery3_pen2 ->(excret: propag_inter) nursery3_pen6 -> (excret: propag_inter) nursery3_pen2"
        - "nursery3_pen2 ->(excret: propag_inter) nursery3_pen7 -> (excret: propag_inter) nursery3_pen2"
        - "nursery3_pen2 ->(excret: propag_inter) nursery3_pen8 -> (excret: propag_inter) nursery3_pen2"
        - "nursery3_pen2 ->(excret: propag_inter) nursery3_pen9 -> (excret: propag_inter) nursery3_pen2"
        - "nursery3_pen3 ->(excret: propag_inter) nursery3_pen5 -> (excret: propag_inter) nursery3_pen3"
        - "nursery3_pen3 ->(excret: propag_inter) nursery3_pen6 -> (excret: propag_inter) nursery3_pen3"
        - "nursery3_pen3 ->(excret: propag_inter) nursery3_pen7 -> (excret: propag_inter) nursery3_pen3"
        - "nursery3_pen3 ->(excret: propag_inter) nursery3_pen8 -> (excret: propag_inter) nursery3_pen3"
        - "nursery3_pen3 ->(excret: propag_inter) nursery3_pen9 -> (excret: propag_inter) nursery3_pen3"
        - "nursery3_pen4 ->(excret: propag_inter) nursery3_pen5 -> (excret: propag_inter) nursery3_pen4"
        - "nursery3_pen4 ->(excret: propag_inter) nursery3_pen6 -> (excret: propag_inter) nursery3_pen4"
        - "nursery3_pen4 ->(excret: propag_inter) nursery3_pen7 -> (excret: propag_inter) nursery3_pen4"
        - "nursery3_pen4 ->(excret: propag_inter) nursery3_pen8 -> (excret: propag_inter) nursery3_pen4"
        - "nursery3_pen4 ->(excret: propag_inter) nursery3_pen9 -> (excret: propag_inter) nursery3_pen4"
        - "nursery3_pen5 ->(excret: propag_inter) nursery3_pen7 -> (excret: propag_inter) nursery3_pen5"
        - "nursery3_pen5 ->(excret: propag_inter) nursery3_pen8 -> (excret: propag_inter) nursery3_pen5"
        - "nursery3_pen5 ->(excret: propag_inter) nursery3_pen9 -> (excret: propag_inter) nursery3_pen5"
        - "nursery3_pen6 ->(excret: propag_inter) nursery3_pen7 -> (excret: propag_inter) nursery3_pen6"
        - "nursery3_pen6 ->(excret: propag_inter) nursery3_pen8 -> (excret: propag_inter) nursery3_pen6"
        - "nursery3_pen6 ->(excret: propag_inter) nursery3_pen9 -> (excret: propag_inter) nursery3_pen6"
        - "nursery3_pen7 ->(excret: propag_inter) nursery3_pen9 -> (excret: propag_inter) nursery3_pen7"
        - "nursery3_pen8 ->(excret: propag_inter) nursery3_pen9 -> (excret: propag_inter) nursery3_pen8"

  finishingSector:
    name: 'finishingSector'
    desc: 'finishing sector'
    execute_process:
      - allocate
    spaces:
      name: 'room'
      descritpion: 'sector is divided in 6 rooms'
      allocation:
        - 'alternate_by_group(group=batch, spaces=ALL)'
      nodes:
        finishing1:
          reference: finishing1
        finishing2:
          reference: finishing2
        finishing3:
          reference: finishing3
        finishing4:
          reference: finishing4
        finishing5:
          reference: finishing5
        finishing6:
          reference: finishing6


  finishing1:
    name: 'finishing 1'
    desc: 'finishing 1'
    processes:
      deposit_pen1:
        action: 'set_to(information=excret, value=total_I_notVac_finishing1_pen1, spaces=[finishing1_pen1])'
      deposit_pen2:
        action: 'set_to(information=excret, value=total_I_notVac_finishing1_pen2, spaces=[finishing1_pen2])'
      deposit_pen3:
        action: 'set_to(information=excret, value=total_I_notVac_finishing1_pen3, spaces=[finishing1_pen3])'
      deposit_pen4:
        action: 'set_to(information=excret, value=total_I_notVac_finishing1_pen4, spaces=[finishing1_pen4])'
      deposit_pen5:
        action: 'set_to(information=excret, value=total_I_notVac_finishing1_pen5, spaces=[finishing1_pen5])'
      deposit_pen6:
        action: 'set_to(information=excret, value=total_I_notVac_finishing1_pen6, spaces=[finishing1_pen6])'
      deposit_pen7:
        action: 'set_to(information=excret, value=total_I_notVac_finishing1_pen7, spaces=[finishing1_pen7])'
      deposit_pen8:
        action: 'set_to(information=excret, value=total_I_notVac_finishing1_pen8, spaces=[finishing1_pen8])'
    execute_process:
      - allocate
      - deposit_pen1
      - deposit_pen2
      - deposit_pen3
      - deposit_pen4
      - deposit_pen5
      - deposit_pen6
      - deposit_pen7
      - deposit_pen8
      - propagate_information
    spaces:
      name: 'pen'
      description: '15 pens'
      allocation:
        - if: 'isin_pen_group1'
          then: finishing1_pen1
        - if: 'isin_pen_group2'
          then: finishing1_pen2
        - if: 'isin_pen_group3'
          then: finishing1_pen3
        - if: 'isin_pen_group4'
          then: finishing1_pen4
        - if: 'isin_pen_group5'
          then: finishing1_pen5
        - if: 'isin_pen_group6'
          then: finishing1_pen6
        - if: 'isin_pen_group7'
          then: finishing1_pen7
        - if: 'isin_pen_group8'
          then: finishing1_pen8
        - if: 'isin_pen_group9'
          then: finishing1_pen9
      nodes:
        finishing1_pen1:
          informations:
            excret:
              value: 0


        finishing1_pen2:
          informations:
            excret:
              value: 0


        finishing1_pen3:
          informations:
            excret:
              value: 0


        finishing1_pen4:
          informations:
            excret:
              value: 0


        finishing1_pen5:
          informations:
            excret:
              value: 0


        finishing1_pen6:
          informations:
            excret:
              value: 0


        finishing1_pen7:
          informations:
            excret:
              value: 0


        finishing1_pen8:
          informations:
            excret:
              value: 0


        finishing1_pen9:
          informations:
            excret:
              value: 0
      graph:
        - "finishing1_pen1 ->(excret: propag_intra) finishing1_pen2 -> (excret: propag_intra) finishing1_pen1"
        - "finishing1_pen3 ->(excret: propag_intra) finishing1_pen4 -> (excret: propag_intra) finishing1_pen3"
        - "finishing1_pen5 ->(excret: propag_intra) finishing1_pen6 -> (excret: propag_intra) finishing1_pen5"
        - "finishing1_pen7 ->(excret: propag_intra) finishing1_pen8 -> (excret: propag_intra) finishing1_pen7"
        - "finishing1_pen1 ->(excret: propag_inter) finishing1_pen3 -> (excret: propag_inter) finishing1_pen1"
        - "finishing1_pen1 ->(excret: propag_inter) finishing1_pen4 -> (excret: propag_inter) finishing1_pen1"
        - "finishing1_pen1 ->(excret: propag_inter) finishing1_pen5 -> (excret: propag_inter) finishing1_pen1"
        - "finishing1_pen1 ->(excret: propag_inter) finishing1_pen6 -> (excret: propag_inter) finishing1_pen1"
        - "finishing1_pen1 ->(excret: propag_inter) finishing1_pen7 -> (excret: propag_inter) finishing1_pen1"
        - "finishing1_pen1 ->(excret: propag_inter) finishing1_pen8 -> (excret: propag_inter) finishing1_pen1"
        - "finishing1_pen1 ->(excret: propag_inter) finishing1_pen9 -> (excret: propag_inter) finishing1_pen1"
        - "finishing1_pen2 ->(excret: propag_inter) finishing1_pen3 -> (excret: propag_inter) finishing1_pen2"
        - "finishing1_pen2 ->(excret: propag_inter) finishing1_pen4 -> (excret: propag_inter) finishing1_pen2"
        - "finishing1_pen2 ->(excret: propag_inter) finishing1_pen5 -> (excret: propag_inter) finishing1_pen2"
        - "finishing1_pen2 ->(excret: propag_inter) finishing1_pen6 -> (excret: propag_inter) finishing1_pen2"
        - "finishing1_pen2 ->(excret: propag_inter) finishing1_pen7 -> (excret: propag_inter) finishing1_pen2"
        - "finishing1_pen2 ->(excret: propag_inter) finishing1_pen8 -> (excret: propag_inter) finishing1_pen2"
        - "finishing1_pen2 ->(excret: propag_inter) finishing1_pen9 -> (excret: propag_inter) finishing1_pen2"
        - "finishing1_pen3 ->(excret: propag_inter) finishing1_pen5 -> (excret: propag_inter) finishing1_pen3"
        - "finishing1_pen3 ->(excret: propag_inter) finishing1_pen6 -> (excret: propag_inter) finishing1_pen3"
        - "finishing1_pen3 ->(excret: propag_inter) finishing1_pen7 -> (excret: propag_inter) finishing1_pen3"
        - "finishing1_pen3 ->(excret: propag_inter) finishing1_pen8 -> (excret: propag_inter) finishing1_pen3"
        - "finishing1_pen3 ->(excret: propag_inter) finishing1_pen9 -> (excret: propag_inter) finishing1_pen3"
        - "finishing1_pen4 ->(excret: propag_inter) finishing1_pen5 -> (excret: propag_inter) finishing1_pen4"
        - "finishing1_pen4 ->(excret: propag_inter) finishing1_pen6 -> (excret: propag_inter) finishing1_pen4"
        - "finishing1_pen4 ->(excret: propag_inter) finishing1_pen7 -> (excret: propag_inter) finishing1_pen4"
        - "finishing1_pen4 ->(excret: propag_inter) finishing1_pen8 -> (excret: propag_inter) finishing1_pen4"
        - "finishing1_pen4 ->(excret: propag_inter) finishing1_pen9 -> (excret: propag_inter) finishing1_pen4"
        - "finishing1_pen5 ->(excret: propag_inter) finishing1_pen7 -> (excret: propag_inter) finishing1_pen5"
        - "finishing1_pen5 ->(excret: propag_inter) finishing1_pen8 -> (excret: propag_inter) finishing1_pen5"
        - "finishing1_pen5 ->(excret: propag_inter) finishing1_pen9 -> (excret: propag_inter) finishing1_pen5"
        - "finishing1_pen6 ->(excret: propag_inter) finishing1_pen7 -> (excret: propag_inter) finishing1_pen6"
        - "finishing1_pen6 ->(excret: propag_inter) finishing1_pen8 -> (excret: propag_inter) finishing1_pen6"
        - "finishing1_pen6 ->(excret: propag_inter) finishing1_pen9 -> (excret: propag_inter) finishing1_pen6"
        - "finishing1_pen7 ->(excret: propag_inter) finishing1_pen9 -> (excret: propag_inter) finishing1_pen7"
        - "finishing1_pen8 ->(excret: propag_inter) finishing1_pen9 -> (excret: propag_inter) finishing1_pen8"

  finishing2:
    name: 'finishing 1'
    desc: 'finishing 1'
    processes:
      deposit_pen1:
        action: 'set_to(information=excret, value=total_I_notVac_finishing2_pen1, spaces=[finishing2_pen1])'
      deposit_pen2:
        action: 'set_to(information=excret, value=total_I_notVac_finishing2_pen2, spaces=[finishing2_pen2])'
      deposit_pen3:
        action: 'set_to(information=excret, value=total_I_notVac_finishing2_pen3, spaces=[finishing2_pen3])'
      deposit_pen4:
        action: 'set_to(information=excret, value=total_I_notVac_finishing2_pen4, spaces=[finishing2_pen4])'
      deposit_pen5:
        action: 'set_to(information=excret, value=total_I_notVac_finishing2_pen5, spaces=[finishing2_pen5])'
      deposit_pen6:
        action: 'set_to(information=excret, value=total_I_notVac_finishing2_pen6, spaces=[finishing2_pen6])'
      deposit_pen7:
        action: 'set_to(information=excret, value=total_I_notVac_finishing2_pen7, spaces=[finishing2_pen7])'
      deposit_pen8:
        action: 'set_to(information=excret, value=total_I_notVac_finishing2_pen8, spaces=[finishing2_pen8])'
    execute_process:
      - allocate
      - deposit_pen1
      - deposit_pen2
      - deposit_pen3
      - deposit_pen4
      - deposit_pen5
      - deposit_pen6
      - deposit_pen7
      - deposit_pen8
      - propagate_information
    spaces:
      name: 'pen'
      description: '15 pens'
      allocation:
        - if: 'isin_pen_group1'
          then: finishing2_pen1
        - if: 'isin_pen_group2'
          then: finishing2_pen2
        - if: 'isin_pen_group3'
          then: finishing2_pen3
        - if: 'isin_pen_group4'
          then: finishing2_pen4
        - if: 'isin_pen_group5'
          then: finishing2_pen5
        - if: 'isin_pen_group6'
          then: finishing2_pen6
        - if: 'isin_pen_group7'
          then: finishing2_pen7
        - if: 'isin_pen_group8'
          then: finishing2_pen8
        - if: 'isin_pen_group9'
          then: finishing2_pen9
      nodes:
        finishing2_pen1:
          informations:
            excret:
              value: 0


        finishing2_pen2:
          informations:
            excret:
              value: 0


        finishing2_pen3:
          informations:
            excret:
              value: 0


        finishing2_pen4:
          informations:
            excret:
              value: 0


        finishing2_pen5:
          informations:
            excret:
              value: 0


        finishing2_pen6:
          informations:
            excret:
              value: 0


        finishing2_pen7:
          informations:
            excret:
              value: 0


        finishing2_pen8:
          informations:
            excret:
              value: 0


        finishing2_pen9:
          informations:
            excret:
              value: 0
      graph:
        - "finishing2_pen1 ->(excret: propag_intra) finishing2_pen2 -> (excret: propag_intra) finishing2_pen1"
        - "finishing2_pen3 ->(excret: propag_intra) finishing2_pen4 -> (excret: propag_intra) finishing2_pen3"
        - "finishing2_pen5 ->(excret: propag_intra) finishing2_pen6 -> (excret: propag_intra) finishing2_pen5"
        - "finishing2_pen7 ->(excret: propag_intra) finishing2_pen8 -> (excret: propag_intra) finishing2_pen7"
        - "finishing2_pen1 ->(excret: propag_inter) finishing2_pen3 -> (excret: propag_inter) finishing2_pen1"
        - "finishing2_pen1 ->(excret: propag_inter) finishing2_pen4 -> (excret: propag_inter) finishing2_pen1"
        - "finishing2_pen1 ->(excret: propag_inter) finishing2_pen5 -> (excret: propag_inter) finishing2_pen1"
        - "finishing2_pen1 ->(excret: propag_inter) finishing2_pen6 -> (excret: propag_inter) finishing2_pen1"
        - "finishing2_pen1 ->(excret: propag_inter) finishing2_pen7 -> (excret: propag_inter) finishing2_pen1"
        - "finishing2_pen1 ->(excret: propag_inter) finishing2_pen8 -> (excret: propag_inter) finishing2_pen1"
        - "finishing2_pen1 ->(excret: propag_inter) finishing2_pen9 -> (excret: propag_inter) finishing2_pen1"
        - "finishing2_pen2 ->(excret: propag_inter) finishing2_pen3 -> (excret: propag_inter) finishing2_pen2"
        - "finishing2_pen2 ->(excret: propag_inter) finishing2_pen4 -> (excret: propag_inter) finishing2_pen2"
        - "finishing2_pen2 ->(excret: propag_inter) finishing2_pen5 -> (excret: propag_inter) finishing2_pen2"
        - "finishing2_pen2 ->(excret: propag_inter) finishing2_pen6 -> (excret: propag_inter) finishing2_pen2"
        - "finishing2_pen2 ->(excret: propag_inter) finishing2_pen7 -> (excret: propag_inter) finishing2_pen2"
        - "finishing2_pen2 ->(excret: propag_inter) finishing2_pen8 -> (excret: propag_inter) finishing2_pen2"
        - "finishing2_pen2 ->(excret: propag_inter) finishing2_pen9 -> (excret: propag_inter) finishing2_pen2"
        - "finishing2_pen3 ->(excret: propag_inter) finishing2_pen5 -> (excret: propag_inter) finishing2_pen3"
        - "finishing2_pen3 ->(excret: propag_inter) finishing2_pen6 -> (excret: propag_inter) finishing2_pen3"
        - "finishing2_pen3 ->(excret: propag_inter) finishing2_pen7 -> (excret: propag_inter) finishing2_pen3"
        - "finishing2_pen3 ->(excret: propag_inter) finishing2_pen8 -> (excret: propag_inter) finishing2_pen3"
        - "finishing2_pen3 ->(excret: propag_inter) finishing2_pen9 -> (excret: propag_inter) finishing2_pen3"
        - "finishing2_pen4 ->(excret: propag_inter) finishing2_pen5 -> (excret: propag_inter) finishing2_pen4"
        - "finishing2_pen4 ->(excret: propag_inter) finishing2_pen6 -> (excret: propag_inter) finishing2_pen4"
        - "finishing2_pen4 ->(excret: propag_inter) finishing2_pen7 -> (excret: propag_inter) finishing2_pen4"
        - "finishing2_pen4 ->(excret: propag_inter) finishing2_pen8 -> (excret: propag_inter) finishing2_pen4"
        - "finishing2_pen4 ->(excret: propag_inter) finishing2_pen9 -> (excret: propag_inter) finishing2_pen4"
        - "finishing2_pen5 ->(excret: propag_inter) finishing2_pen7 -> (excret: propag_inter) finishing2_pen5"
        - "finishing2_pen5 ->(excret: propag_inter) finishing2_pen8 -> (excret: propag_inter) finishing2_pen5"
        - "finishing2_pen5 ->(excret: propag_inter) finishing2_pen9 -> (excret: propag_inter) finishing2_pen5"
        - "finishing2_pen6 ->(excret: propag_inter) finishing2_pen7 -> (excret: propag_inter) finishing2_pen6"
        - "finishing2_pen6 ->(excret: propag_inter) finishing2_pen8 -> (excret: propag_inter) finishing2_pen6"
        - "finishing2_pen6 ->(excret: propag_inter) finishing2_pen9 -> (excret: propag_inter) finishing2_pen6"
        - "finishing2_pen7 ->(excret: propag_inter) finishing2_pen9 -> (excret: propag_inter) finishing2_pen7"
        - "finishing2_pen8 ->(excret: propag_inter) finishing2_pen9 -> (excret: propag_inter) finishing2_pen8"

  finishing3:
    name: 'finishing 1'
    desc: 'finishing 1'
    processes:
      deposit_pen1:
        action: 'set_to(information=excret, value=total_I_notVac_finishing3_pen1, spaces=[finishing3_pen1])'
      deposit_pen2:
        action: 'set_to(information=excret, value=total_I_notVac_finishing3_pen2, spaces=[finishing3_pen2])'
      deposit_pen3:
        action: 'set_to(information=excret, value=total_I_notVac_finishing3_pen3, spaces=[finishing3_pen3])'
      deposit_pen4:
        action: 'set_to(information=excret, value=total_I_notVac_finishing3_pen4, spaces=[finishing3_pen4])'
      deposit_pen5:
        action: 'set_to(information=excret, value=total_I_notVac_finishing3_pen5, spaces=[finishing3_pen5])'
      deposit_pen6:
        action: 'set_to(information=excret, value=total_I_notVac_finishing3_pen6, spaces=[finishing3_pen6])'
      deposit_pen7:
        action: 'set_to(information=excret, value=total_I_notVac_finishing3_pen7, spaces=[finishing3_pen7])'
      deposit_pen8:
        action: 'set_to(information=excret, value=total_I_notVac_finishing3_pen8, spaces=[finishing3_pen8])'
    execute_process:
      - allocate
      - deposit_pen1
      - deposit_pen2
      - deposit_pen3
      - deposit_pen4
      - deposit_pen5
      - deposit_pen6
      - deposit_pen7
      - deposit_pen8
      - propagate_information
    spaces:
      name: 'pen'
      description: '15 pens'
      allocation:
        - if: 'isin_pen_group1'
          then: finishing3_pen1
        - if: 'isin_pen_group2'
          then: finishing3_pen2
        - if: 'isin_pen_group3'
          then: finishing3_pen3
        - if: 'isin_pen_group4'
          then: finishing3_pen4
        - if: 'isin_pen_group5'
          then: finishing3_pen5
        - if: 'isin_pen_group6'
          then: finishing3_pen6
        - if: 'isin_pen_group7'
          then: finishing3_pen7
        - if: 'isin_pen_group8'
          then: finishing3_pen8
        - if: 'isin_pen_group9'
          then: finishing3_pen9
      nodes:
        finishing3_pen1:
          informations:
            excret:
              value: 0


        finishing3_pen2:
          informations:
            excret:
              value: 0


        finishing3_pen3:
          informations:
            excret:
              value: 0


        finishing3_pen4:
          informations:
            excret:
              value: 0


        finishing3_pen5:
          informations:
            excret:
              value: 0


        finishing3_pen6:
          informations:
            excret:
              value: 0


        finishing3_pen7:
          informations:
            excret:
              value: 0


        finishing3_pen8:
          informations:
            excret:
              value: 0


        finishing3_pen9:
          informations:
            excret:
              value: 0
      graph:
        - "finishing3_pen1 ->(excret: propag_intra) finishing3_pen2 -> (excret: propag_intra) finishing3_pen1"
        - "finishing3_pen3 ->(excret: propag_intra) finishing3_pen4 -> (excret: propag_intra) finishing3_pen3"
        - "finishing3_pen5 ->(excret: propag_intra) finishing3_pen6 -> (excret: propag_intra) finishing3_pen5"
        - "finishing3_pen7 ->(excret: propag_intra) finishing3_pen8 -> (excret: propag_intra) finishing3_pen7"
        - "finishing3_pen1 ->(excret: propag_inter) finishing3_pen3 -> (excret: propag_inter) finishing3_pen1"
        - "finishing3_pen1 ->(excret: propag_inter) finishing3_pen4 -> (excret: propag_inter) finishing3_pen1"
        - "finishing3_pen1 ->(excret: propag_inter) finishing3_pen5 -> (excret: propag_inter) finishing3_pen1"
        - "finishing3_pen1 ->(excret: propag_inter) finishing3_pen6 -> (excret: propag_inter) finishing3_pen1"
        - "finishing3_pen1 ->(excret: propag_inter) finishing3_pen7 -> (excret: propag_inter) finishing3_pen1"
        - "finishing3_pen1 ->(excret: propag_inter) finishing3_pen8 -> (excret: propag_inter) finishing3_pen1"
        - "finishing3_pen1 ->(excret: propag_inter) finishing3_pen9 -> (excret: propag_inter) finishing3_pen1"
        - "finishing3_pen2 ->(excret: propag_inter) finishing3_pen3 -> (excret: propag_inter) finishing3_pen2"
        - "finishing3_pen2 ->(excret: propag_inter) finishing3_pen4 -> (excret: propag_inter) finishing3_pen2"
        - "finishing3_pen2 ->(excret: propag_inter) finishing3_pen5 -> (excret: propag_inter) finishing3_pen2"
        - "finishing3_pen2 ->(excret: propag_inter) finishing3_pen6 -> (excret: propag_inter) finishing3_pen2"
        - "finishing3_pen2 ->(excret: propag_inter) finishing3_pen7 -> (excret: propag_inter) finishing3_pen2"
        - "finishing3_pen2 ->(excret: propag_inter) finishing3_pen8 -> (excret: propag_inter) finishing3_pen2"
        - "finishing3_pen2 ->(excret: propag_inter) finishing3_pen9 -> (excret: propag_inter) finishing3_pen2"
        - "finishing3_pen3 ->(excret: propag_inter) finishing3_pen5 -> (excret: propag_inter) finishing3_pen3"
        - "finishing3_pen3 ->(excret: propag_inter) finishing3_pen6 -> (excret: propag_inter) finishing3_pen3"
        - "finishing3_pen3 ->(excret: propag_inter) finishing3_pen7 -> (excret: propag_inter) finishing3_pen3"
        - "finishing3_pen3 ->(excret: propag_inter) finishing3_pen8 -> (excret: propag_inter) finishing3_pen3"
        - "finishing3_pen3 ->(excret: propag_inter) finishing3_pen9 -> (excret: propag_inter) finishing3_pen3"
        - "finishing3_pen4 ->(excret: propag_inter) finishing3_pen5 -> (excret: propag_inter) finishing3_pen4"
        - "finishing3_pen4 ->(excret: propag_inter) finishing3_pen6 -> (excret: propag_inter) finishing3_pen4"
        - "finishing3_pen4 ->(excret: propag_inter) finishing3_pen7 -> (excret: propag_inter) finishing3_pen4"
        - "finishing3_pen4 ->(excret: propag_inter) finishing3_pen8 -> (excret: propag_inter) finishing3_pen4"
        - "finishing3_pen4 ->(excret: propag_inter) finishing3_pen9 -> (excret: propag_inter) finishing3_pen4"
        - "finishing3_pen5 ->(excret: propag_inter) finishing3_pen7 -> (excret: propag_inter) finishing3_pen5"
        - "finishing3_pen5 ->(excret: propag_inter) finishing3_pen8 -> (excret: propag_inter) finishing3_pen5"
        - "finishing3_pen5 ->(excret: propag_inter) finishing3_pen9 -> (excret: propag_inter) finishing3_pen5"
        - "finishing3_pen6 ->(excret: propag_inter) finishing3_pen7 -> (excret: propag_inter) finishing3_pen6"
        - "finishing3_pen6 ->(excret: propag_inter) finishing3_pen8 -> (excret: propag_inter) finishing3_pen6"
        - "finishing3_pen6 ->(excret: propag_inter) finishing3_pen9 -> (excret: propag_inter) finishing3_pen6"
        - "finishing3_pen7 ->(excret: propag_inter) finishing3_pen9 -> (excret: propag_inter) finishing3_pen7"
        - "finishing3_pen8 ->(excret: propag_inter) finishing3_pen9 -> (excret: propag_inter) finishing3_pen8"

  finishing4:
    name: 'finishing 1'
    desc: 'finishing 1'
    processes:
      deposit_pen1:
        action: 'set_to(information=excret, value=total_I_notVac_finishing4_pen1, spaces=[finishing4_pen1])'
      deposit_pen2:
        action: 'set_to(information=excret, value=total_I_notVac_finishing4_pen2, spaces=[finishing4_pen2])'
      deposit_pen3:
        action: 'set_to(information=excret, value=total_I_notVac_finishing4_pen3, spaces=[finishing4_pen3])'
      deposit_pen4:
        action: 'set_to(information=excret, value=total_I_notVac_finishing4_pen4, spaces=[finishing4_pen4])'
      deposit_pen5:
        action: 'set_to(information=excret, value=total_I_notVac_finishing4_pen5, spaces=[finishing4_pen5])'
      deposit_pen6:
        action: 'set_to(information=excret, value=total_I_notVac_finishing4_pen6, spaces=[finishing4_pen6])'
      deposit_pen7:
        action: 'set_to(information=excret, value=total_I_notVac_finishing4_pen7, spaces=[finishing4_pen7])'
      deposit_pen8:
        action: 'set_to(information=excret, value=total_I_notVac_finishing4_pen8, spaces=[finishing4_pen8])'
    execute_process:
      - allocate
      - deposit_pen1
      - deposit_pen2
      - deposit_pen3
      - deposit_pen4
      - deposit_pen5
      - deposit_pen6
      - deposit_pen7
      - deposit_pen8
      - propagate_information
    spaces:
      name: 'pen'
      description: '15 pens'
      allocation:
        - if: 'isin_pen_group1'
          then: finishing4_pen1
        - if: 'isin_pen_group2'
          then: finishing4_pen2
        - if: 'isin_pen_group3'
          then: finishing4_pen3
        - if: 'isin_pen_group4'
          then: finishing4_pen4
        - if: 'isin_pen_group5'
          then: finishing4_pen5
        - if: 'isin_pen_group6'
          then: finishing4_pen6
        - if: 'isin_pen_group7'
          then: finishing4_pen7
        - if: 'isin_pen_group8'
          then: finishing4_pen8
        - if: 'isin_pen_group9'
          then: finishing4_pen9
      nodes:
        finishing4_pen1:
          informations:
            excret:
              value: 0


        finishing4_pen2:
          informations:
            excret:
              value: 0


        finishing4_pen3:
          informations:
            excret:
              value: 0


        finishing4_pen4:
          informations:
            excret:
              value: 0


        finishing4_pen5:
          informations:
            excret:
              value: 0


        finishing4_pen6:
          informations:
            excret:
              value: 0


        finishing4_pen7:
          informations:
            excret:
              value: 0


        finishing4_pen8:
          informations:
            excret:
              value: 0


        finishing4_pen9:
          informations:
            excret:
              value: 0
      graph:
        - "finishing4_pen1 ->(excret: propag_intra) finishing4_pen2 -> (excret: propag_intra) finishing4_pen1"
        - "finishing4_pen3 ->(excret: propag_intra) finishing4_pen4 -> (excret: propag_intra) finishing4_pen3"
        - "finishing4_pen5 ->(excret: propag_intra) finishing4_pen6 -> (excret: propag_intra) finishing4_pen5"
        - "finishing4_pen7 ->(excret: propag_intra) finishing4_pen8 -> (excret: propag_intra) finishing4_pen7"
        - "finishing4_pen1 ->(excret: propag_inter) finishing4_pen3 -> (excret: propag_inter) finishing4_pen1"
        - "finishing4_pen1 ->(excret: propag_inter) finishing4_pen4 -> (excret: propag_inter) finishing4_pen1"
        - "finishing4_pen1 ->(excret: propag_inter) finishing4_pen5 -> (excret: propag_inter) finishing4_pen1"
        - "finishing4_pen1 ->(excret: propag_inter) finishing4_pen6 -> (excret: propag_inter) finishing4_pen1"
        - "finishing4_pen1 ->(excret: propag_inter) finishing4_pen7 -> (excret: propag_inter) finishing4_pen1"
        - "finishing4_pen1 ->(excret: propag_inter) finishing4_pen8 -> (excret: propag_inter) finishing4_pen1"
        - "finishing4_pen1 ->(excret: propag_inter) finishing4_pen9 -> (excret: propag_inter) finishing4_pen1"
        - "finishing4_pen2 ->(excret: propag_inter) finishing4_pen3 -> (excret: propag_inter) finishing4_pen2"
        - "finishing4_pen2 ->(excret: propag_inter) finishing4_pen4 -> (excret: propag_inter) finishing4_pen2"
        - "finishing4_pen2 ->(excret: propag_inter) finishing4_pen5 -> (excret: propag_inter) finishing4_pen2"
        - "finishing4_pen2 ->(excret: propag_inter) finishing4_pen6 -> (excret: propag_inter) finishing4_pen2"
        - "finishing4_pen2 ->(excret: propag_inter) finishing4_pen7 -> (excret: propag_inter) finishing4_pen2"
        - "finishing4_pen2 ->(excret: propag_inter) finishing4_pen8 -> (excret: propag_inter) finishing4_pen2"
        - "finishing4_pen2 ->(excret: propag_inter) finishing4_pen9 -> (excret: propag_inter) finishing4_pen2"
        - "finishing4_pen3 ->(excret: propag_inter) finishing4_pen5 -> (excret: propag_inter) finishing4_pen3"
        - "finishing4_pen3 ->(excret: propag_inter) finishing4_pen6 -> (excret: propag_inter) finishing4_pen3"
        - "finishing4_pen3 ->(excret: propag_inter) finishing4_pen7 -> (excret: propag_inter) finishing4_pen3"
        - "finishing4_pen3 ->(excret: propag_inter) finishing4_pen8 -> (excret: propag_inter) finishing4_pen3"
        - "finishing4_pen3 ->(excret: propag_inter) finishing4_pen9 -> (excret: propag_inter) finishing4_pen3"
        - "finishing4_pen4 ->(excret: propag_inter) finishing4_pen5 -> (excret: propag_inter) finishing4_pen4"
        - "finishing4_pen4 ->(excret: propag_inter) finishing4_pen6 -> (excret: propag_inter) finishing4_pen4"
        - "finishing4_pen4 ->(excret: propag_inter) finishing4_pen7 -> (excret: propag_inter) finishing4_pen4"
        - "finishing4_pen4 ->(excret: propag_inter) finishing4_pen8 -> (excret: propag_inter) finishing4_pen4"
        - "finishing4_pen4 ->(excret: propag_inter) finishing4_pen9 -> (excret: propag_inter) finishing4_pen4"
        - "finishing4_pen5 ->(excret: propag_inter) finishing4_pen7 -> (excret: propag_inter) finishing4_pen5"
        - "finishing4_pen5 ->(excret: propag_inter) finishing4_pen8 -> (excret: propag_inter) finishing4_pen5"
        - "finishing4_pen5 ->(excret: propag_inter) finishing4_pen9 -> (excret: propag_inter) finishing4_pen5"
        - "finishing4_pen6 ->(excret: propag_inter) finishing4_pen7 -> (excret: propag_inter) finishing4_pen6"
        - "finishing4_pen6 ->(excret: propag_inter) finishing4_pen8 -> (excret: propag_inter) finishing4_pen6"
        - "finishing4_pen6 ->(excret: propag_inter) finishing4_pen9 -> (excret: propag_inter) finishing4_pen6"
        - "finishing4_pen7 ->(excret: propag_inter) finishing4_pen9 -> (excret: propag_inter) finishing4_pen7"
        - "finishing4_pen8 ->(excret: propag_inter) finishing4_pen9 -> (excret: propag_inter) finishing4_pen8"


  finishing5:
    name: 'finishing 1'
    desc: 'finishing 1'
    processes:
      deposit_pen1:
        action: 'set_to(information=excret, value=total_I_notVac_finishing5_pen1, spaces=[finishing5_pen1])'
      deposit_pen2:
        action: 'set_to(information=excret, value=total_I_notVac_finishing5_pen2, spaces=[finishing5_pen2])'
      deposit_pen3:
        action: 'set_to(information=excret, value=total_I_notVac_finishing5_pen3, spaces=[finishing5_pen3])'
      deposit_pen4:
        action: 'set_to(information=excret, value=total_I_notVac_finishing5_pen4, spaces=[finishing5_pen4])'
      deposit_pen5:
        action: 'set_to(information=excret, value=total_I_notVac_finishing5_pen5, spaces=[finishing5_pen5])'
      deposit_pen6:
        action: 'set_to(information=excret, value=total_I_notVac_finishing5_pen6, spaces=[finishing5_pen6])'
      deposit_pen7:
        action: 'set_to(information=excret, value=total_I_notVac_finishing5_pen7, spaces=[finishing5_pen7])'
      deposit_pen8:
        action: 'set_to(information=excret, value=total_I_notVac_finishing5_pen8, spaces=[finishing5_pen8])'
    execute_process:
      - allocate
      - deposit_pen1
      - deposit_pen2
      - deposit_pen3
      - deposit_pen4
      - deposit_pen5
      - deposit_pen6
      - deposit_pen7
      - deposit_pen8
      - propagate_information
    spaces:
      name: 'pen'
      description: '15 pens'
      allocation:
        - if: 'isin_pen_group1'
          then: finishing5_pen1
        - if: 'isin_pen_group2'
          then: finishing5_pen2
        - if: 'isin_pen_group3'
          then: finishing5_pen3
        - if: 'isin_pen_group4'
          then: finishing5_pen4
        - if: 'isin_pen_group5'
          then: finishing5_pen5
        - if: 'isin_pen_group6'
          then: finishing5_pen6
        - if: 'isin_pen_group7'
          then: finishing5_pen7
        - if: 'isin_pen_group8'
          then: finishing5_pen8
        - if: 'isin_pen_group9'
          then: finishing5_pen9
      nodes:
        finishing5_pen1:
          informations:
            excret:
              value: 0


        finishing5_pen2:
          informations:
            excret:
              value: 0


        finishing5_pen3:
          informations:
            excret:
              value: 0


        finishing5_pen4:
          informations:
            excret:
              value: 0


        finishing5_pen5:
          informations:
            excret:
              value: 0


        finishing5_pen6:
          informations:
            excret:
              value: 0


        finishing5_pen7:
          informations:
            excret:
              value: 0


        finishing5_pen8:
          informations:
            excret:
              value: 0


        finishing5_pen9:
          informations:
            excret:
              value: 0
      graph:
        - "finishing5_pen1 ->(excret: propag_intra) finishing5_pen2 -> (excret: propag_intra) finishing5_pen1"
        - "finishing5_pen3 ->(excret: propag_intra) finishing5_pen4 -> (excret: propag_intra) finishing5_pen3"
        - "finishing5_pen5 ->(excret: propag_intra) finishing5_pen6 -> (excret: propag_intra) finishing5_pen5"
        - "finishing5_pen7 ->(excret: propag_intra) finishing5_pen8 -> (excret: propag_intra) finishing5_pen7"
        - "finishing5_pen1 ->(excret: propag_inter) finishing5_pen3 -> (excret: propag_inter) finishing5_pen1"
        - "finishing5_pen1 ->(excret: propag_inter) finishing5_pen4 -> (excret: propag_inter) finishing5_pen1"
        - "finishing5_pen1 ->(excret: propag_inter) finishing5_pen5 -> (excret: propag_inter) finishing5_pen1"
        - "finishing5_pen1 ->(excret: propag_inter) finishing5_pen6 -> (excret: propag_inter) finishing5_pen1"
        - "finishing5_pen1 ->(excret: propag_inter) finishing5_pen7 -> (excret: propag_inter) finishing5_pen1"
        - "finishing5_pen1 ->(excret: propag_inter) finishing5_pen8 -> (excret: propag_inter) finishing5_pen1"
        - "finishing5_pen1 ->(excret: propag_inter) finishing5_pen9 -> (excret: propag_inter) finishing5_pen1"
        - "finishing5_pen2 ->(excret: propag_inter) finishing5_pen3 -> (excret: propag_inter) finishing5_pen2"
        - "finishing5_pen2 ->(excret: propag_inter) finishing5_pen4 -> (excret: propag_inter) finishing5_pen2"
        - "finishing5_pen2 ->(excret: propag_inter) finishing5_pen5 -> (excret: propag_inter) finishing5_pen2"
        - "finishing5_pen2 ->(excret: propag_inter) finishing5_pen6 -> (excret: propag_inter) finishing5_pen2"
        - "finishing5_pen2 ->(excret: propag_inter) finishing5_pen7 -> (excret: propag_inter) finishing5_pen2"
        - "finishing5_pen2 ->(excret: propag_inter) finishing5_pen8 -> (excret: propag_inter) finishing5_pen2"
        - "finishing5_pen2 ->(excret: propag_inter) finishing5_pen9 -> (excret: propag_inter) finishing5_pen2"
        - "finishing5_pen3 ->(excret: propag_inter) finishing5_pen5 -> (excret: propag_inter) finishing5_pen3"
        - "finishing5_pen3 ->(excret: propag_inter) finishing5_pen6 -> (excret: propag_inter) finishing5_pen3"
        - "finishing5_pen3 ->(excret: propag_inter) finishing5_pen7 -> (excret: propag_inter) finishing5_pen3"
        - "finishing5_pen3 ->(excret: propag_inter) finishing5_pen8 -> (excret: propag_inter) finishing5_pen3"
        - "finishing5_pen3 ->(excret: propag_inter) finishing5_pen9 -> (excret: propag_inter) finishing5_pen3"
        - "finishing5_pen4 ->(excret: propag_inter) finishing5_pen5 -> (excret: propag_inter) finishing5_pen4"
        - "finishing5_pen4 ->(excret: propag_inter) finishing5_pen6 -> (excret: propag_inter) finishing5_pen4"
        - "finishing5_pen4 ->(excret: propag_inter) finishing5_pen7 -> (excret: propag_inter) finishing5_pen4"
        - "finishing5_pen4 ->(excret: propag_inter) finishing5_pen8 -> (excret: propag_inter) finishing5_pen4"
        - "finishing5_pen4 ->(excret: propag_inter) finishing5_pen9 -> (excret: propag_inter) finishing5_pen4"
        - "finishing5_pen5 ->(excret: propag_inter) finishing5_pen7 -> (excret: propag_inter) finishing5_pen5"
        - "finishing5_pen5 ->(excret: propag_inter) finishing5_pen8 -> (excret: propag_inter) finishing5_pen5"
        - "finishing5_pen5 ->(excret: propag_inter) finishing5_pen9 -> (excret: propag_inter) finishing5_pen5"
        - "finishing5_pen6 ->(excret: propag_inter) finishing5_pen7 -> (excret: propag_inter) finishing5_pen6"
        - "finishing5_pen6 ->(excret: propag_inter) finishing5_pen8 -> (excret: propag_inter) finishing5_pen6"
        - "finishing5_pen6 ->(excret: propag_inter) finishing5_pen9 -> (excret: propag_inter) finishing5_pen6"
        - "finishing5_pen7 ->(excret: propag_inter) finishing5_pen9 -> (excret: propag_inter) finishing5_pen7"
        - "finishing5_pen8 ->(excret: propag_inter) finishing5_pen9 -> (excret: propag_inter) finishing5_pen8"


  finishing6:
    name: 'finishing 1'
    desc: 'finishing 1'
    processes:
      deposit_pen1:
        action: 'set_to(information=excret, value=total_I_notVac_finishing6_pen1, spaces=[finishing6_pen1])'
      deposit_pen2:
        action: 'set_to(information=excret, value=total_I_notVac_finishing6_pen2, spaces=[finishing6_pen2])'
      deposit_pen3:
        action: 'set_to(information=excret, value=total_I_notVac_finishing6_pen3, spaces=[finishing6_pen3])'
      deposit_pen4:
        action: 'set_to(information=excret, value=total_I_notVac_finishing6_pen4, spaces=[finishing6_pen4])'
      deposit_pen5:
        action: 'set_to(information=excret, value=total_I_notVac_finishing6_pen5, spaces=[finishing6_pen5])'
      deposit_pen6:
        action: 'set_to(information=excret, value=total_I_notVac_finishing6_pen6, spaces=[finishing6_pen6])'
      deposit_pen7:
        action: 'set_to(information=excret, value=total_I_notVac_finishing6_pen7, spaces=[finishing6_pen7])'
      deposit_pen8:
        action: 'set_to(information=excret, value=total_I_notVac_finishing6_pen8, spaces=[finishing6_pen8])'
    execute_process:
      - allocate
      - deposit_pen1
      - deposit_pen2
      - deposit_pen3
      - deposit_pen4
      - deposit_pen5
      - deposit_pen6
      - deposit_pen7
      - deposit_pen8
      - propagate_information
    spaces:
      name: 'pen'
      description: '15 pens'
      allocation:
        - if: 'isin_pen_group1'
          then: finishing6_pen1
        - if: 'isin_pen_group2'
          then: finishing6_pen2
        - if: 'isin_pen_group3'
          then: finishing6_pen3
        - if: 'isin_pen_group4'
          then: finishing6_pen4
        - if: 'isin_pen_group5'
          then: finishing6_pen5
        - if: 'isin_pen_group6'
          then: finishing6_pen6
        - if: 'isin_pen_group7'
          then: finishing6_pen7
        - if: 'isin_pen_group8'
          then: finishing6_pen8
        - if: 'isin_pen_group9'
          then: finishing6_pen9
      nodes:
        finishing6_pen1:
          informations:
            excret:
              value: 0


        finishing6_pen2:
          informations:
            excret:
              value: 0


        finishing6_pen3:
          informations:
            excret:
              value: 0


        finishing6_pen4:
          informations:
            excret:
              value: 0


        finishing6_pen5:
          informations:
            excret:
              value: 0


        finishing6_pen6:
          informations:
            excret:
              value: 0


        finishing6_pen7:
          informations:
            excret:
              value: 0


        finishing6_pen8:
          informations:
            excret:
              value: 0


        finishing6_pen9:
          informations:
            excret:
              value: 0
      graph:
        - "finishing6_pen1 ->(excret: propag_intra) finishing6_pen2 -> (excret: propag_intra) finishing6_pen1"
        - "finishing6_pen3 ->(excret: propag_intra) finishing6_pen4 -> (excret: propag_intra) finishing6_pen3"
        - "finishing6_pen5 ->(excret: propag_intra) finishing6_pen6 -> (excret: propag_intra) finishing6_pen5"
        - "finishing6_pen7 ->(excret: propag_intra) finishing6_pen8 -> (excret: propag_intra) finishing6_pen7"
        - "finishing6_pen1 ->(excret: propag_inter) finishing6_pen3 -> (excret: propag_inter) finishing6_pen1"
        - "finishing6_pen1 ->(excret: propag_inter) finishing6_pen4 -> (excret: propag_inter) finishing6_pen1"
        - "finishing6_pen1 ->(excret: propag_inter) finishing6_pen5 -> (excret: propag_inter) finishing6_pen1"
        - "finishing6_pen1 ->(excret: propag_inter) finishing6_pen6 -> (excret: propag_inter) finishing6_pen1"
        - "finishing6_pen1 ->(excret: propag_inter) finishing6_pen7 -> (excret: propag_inter) finishing6_pen1"
        - "finishing6_pen1 ->(excret: propag_inter) finishing6_pen8 -> (excret: propag_inter) finishing6_pen1"
        - "finishing6_pen1 ->(excret: propag_inter) finishing6_pen9 -> (excret: propag_inter) finishing6_pen1"
        - "finishing6_pen2 ->(excret: propag_inter) finishing6_pen3 -> (excret: propag_inter) finishing6_pen2"
        - "finishing6_pen2 ->(excret: propag_inter) finishing6_pen4 -> (excret: propag_inter) finishing6_pen2"
        - "finishing6_pen2 ->(excret: propag_inter) finishing6_pen5 -> (excret: propag_inter) finishing6_pen2"
        - "finishing6_pen2 ->(excret: propag_inter) finishing6_pen6 -> (excret: propag_inter) finishing6_pen2"
        - "finishing6_pen2 ->(excret: propag_inter) finishing6_pen7 -> (excret: propag_inter) finishing6_pen2"
        - "finishing6_pen2 ->(excret: propag_inter) finishing6_pen8 -> (excret: propag_inter) finishing6_pen2"
        - "finishing6_pen2 ->(excret: propag_inter) finishing6_pen9 -> (excret: propag_inter) finishing6_pen2"
        - "finishing6_pen3 ->(excret: propag_inter) finishing6_pen5 -> (excret: propag_inter) finishing6_pen3"
        - "finishing6_pen3 ->(excret: propag_inter) finishing6_pen6 -> (excret: propag_inter) finishing6_pen3"
        - "finishing6_pen3 ->(excret: propag_inter) finishing6_pen7 -> (excret: propag_inter) finishing6_pen3"
        - "finishing6_pen3 ->(excret: propag_inter) finishing6_pen8 -> (excret: propag_inter) finishing6_pen3"
        - "finishing6_pen3 ->(excret: propag_inter) finishing6_pen9 -> (excret: propag_inter) finishing6_pen3"
        - "finishing6_pen4 ->(excret: propag_inter) finishing6_pen5 -> (excret: propag_inter) finishing6_pen4"
        - "finishing6_pen4 ->(excret: propag_inter) finishing6_pen6 -> (excret: propag_inter) finishing6_pen4"
        - "finishing6_pen4 ->(excret: propag_inter) finishing6_pen7 -> (excret: propag_inter) finishing6_pen4"
        - "finishing6_pen4 ->(excret: propag_inter) finishing6_pen8 -> (excret: propag_inter) finishing6_pen4"
        - "finishing6_pen4 ->(excret: propag_inter) finishing6_pen9 -> (excret: propag_inter) finishing6_pen4"
        - "finishing6_pen5 ->(excret: propag_inter) finishing6_pen7 -> (excret: propag_inter) finishing6_pen5"
        - "finishing6_pen5 ->(excret: propag_inter) finishing6_pen8 -> (excret: propag_inter) finishing6_pen5"
        - "finishing6_pen5 ->(excret: propag_inter) finishing6_pen9 -> (excret: propag_inter) finishing6_pen5"
        - "finishing6_pen6 ->(excret: propag_inter) finishing6_pen7 -> (excret: propag_inter) finishing6_pen6"
        - "finishing6_pen6 ->(excret: propag_inter) finishing6_pen8 -> (excret: propag_inter) finishing6_pen6"
        - "finishing6_pen6 ->(excret: propag_inter) finishing6_pen9 -> (excret: propag_inter) finishing6_pen6"
        - "finishing6_pen7 ->(excret: propag_inter) finishing6_pen9 -> (excret: propag_inter) finishing6_pen7"
        - "finishing6_pen8 ->(excret: propag_inter) finishing6_pen9 -> (excret: propag_inter) finishing6_pen8"


parameters:
  withE:
    desc: 'to swith between considering E (1) or not (0)'
    value: 0
  is_redoub:
    desc: 'redoubling if insemination failes'
    value: 'false'
  propag_intra:
    desc: 'rate for intra pen spread'
    value: 1
  propag_inter:
    desc: 'rate for inter pen spread'
    value: '0'


  initial_size_sow:
    desc: 'number of sows in each batch'
    value: 14
  initial_I_sow:
    desc: 'number of infected sows'
    value: 1

  fertility:
    desc: 'number of piglets per sow'
    value: 12

  nb_renew:
    desc: 'number of sows to be renew'
    value: 2
  nb_rank_to_cull:
    desc: 'rank number beyond which animals can be culled'
    value: 5
  nb_failed_to_cull:
    desc: 'number of failed insemination beyond animals can be culled'
    value: 2
  max_failed_ins:
    desc: 'maximum failed insemination'
    value: 2

  delay_introduction:
    desc: 'delay before introduce an infected sow'
    value: '(insemination_duration+gestating_duration+farrowing_duration) * 2'

  insemination_duration:
    desc: 'state duration'
    value: 34
  gestating_duration:
    desc: 'state duration'
    value: 78
  farrowing_duration:
    desc: 'state duration'
    value: "IfThenElse(is_Sow, 35, weaning_age)"
  post_weaning_duration:
    desc: 'state duration'
    value: "40"
  finishing_duration:
    desc: 'state duration'
    value: "114"

  weaning_age:
    desc: 'age of weaning'
    value: 28

  gestation_period:
    desc: 'duration of the gestation'
    value: 'insemination_duration + gestating_duration'

  decay_exit:
    desc: 'decay to exit'
    value: 'isin_batch1*0 +
    isin_batch2*(batch_interval*1) +
    isin_batch3*(batch_interval*2) +
    isin_batch4*(batch_interval*3) +
    isin_batch5*(batch_interval*4) +
    isin_batch6*(batch_interval*5) +
    isin_batch7*(batch_interval*6) '

  real_gest_duration:
    desc: 'Actual gestation period (considering an insemination at seven days, and the weaning age)'
    value: '(insemination_duration-7)+gestating_duration+(farrowing_duration-weaning_age)'
  first_gestating_duration:
    desc: 'duration of the first period of gestating (add 7 days, as insemination occurs 7 days after entering in insemination sector)'
    value: '7 + (real_gest_duration // 3)'
  second_gestating_duration:
    desc: 'duration of the second period of gestating (adjust time duration if real_gest_duration is not divisble by 3)'
    value: '(real_gest_duration // 3) + IfThenElse((real_gest_duration % 3) > 0,1,0)'
  third_gestating_duration:
    desc: 'duration of the third period of gestating'
    value: '(real_gest_duration // 3)'

  first_gest:
    desc: ''
    value: 'is_first_gestating_period * is_Sow'
  second_gest:
    desc: ''
    value: 'is_second_gestating_period * is_Sow'
  third_gest:
    desc: ''
    value: 'is_third_gestating_period * is_Sow'

  prod_S:
    desc: ''
    value: 'is_produce_S'
  prod_I:
    desc: ''
    value: 'is_produce_I'
  prod_M:
    desc: ''
    value: 'is_produce_M'

  batch_interval:
    desc: 'interval between two successive batches of sows and pigs (days)'
    value: 21


  condition_to_cull:
    desc: 'condition to cull'
    value: 'OR(farrowing_rank >= nb_rank_to_cull, nb_failed_insemination >= nb_failed_to_cull)'

  proba_to_cull:
    desc: 'Only sows are culled. ST cannot be culled of course'
    value: '(1 - is_ST) * is_Sow * condition_to_cull * (
        isin_batch1 * (total_to_cull_batch1 < MAX(0, nb_renew - MAX(0, (initial_size_sow - total_Sow_batch1)))) +
        isin_batch2 * (total_to_cull_batch2 < MAX(0, nb_renew - MAX(0, (initial_size_sow - total_Sow_batch2)))) +
        isin_batch3 * (total_to_cull_batch3 < MAX(0, nb_renew - MAX(0, (initial_size_sow - total_Sow_batch3)))) +
        isin_batch4 * (total_to_cull_batch4 < MAX(0, nb_renew - MAX(0, (initial_size_sow - total_Sow_batch4)))) +
        isin_batch5 * (total_to_cull_batch5 < MAX(0, nb_renew - MAX(0, (initial_size_sow - total_Sow_batch5)))) +
        isin_batch6 * (total_to_cull_batch6 < MAX(0, nb_renew - MAX(0, (initial_size_sow - total_Sow_batch6)))) +
        isin_batch7 * (total_to_cull_batch7 < MAX(0, nb_renew - MAX(0, (initial_size_sow - total_Sow_batch7))))
      )'

  proba_produce:
    desc: 'probabilty to produce new Sow for turnover'
    value: 'is_Sow * (
              isin_batch1 * (total_Sow_batch1 < initial_size_sow) +
              isin_batch2 * (total_Sow_batch2 < initial_size_sow) +
              isin_batch3 * (total_Sow_batch3 < initial_size_sow) +
              isin_batch4 * (total_Sow_batch4 < initial_size_sow) +
              isin_batch5 * (total_Sow_batch5 < initial_size_sow) +
              isin_batch6 * (total_Sow_batch6 < initial_size_sow) +
              isin_batch7 * (total_Sow_batch7 < initial_size_sow)
            )'

  limit_failed_ins:
    desc: 'limit to failed insemination'
    value: '((isin_batch1 * total_failureIns_batch3) +
            (isin_batch2 * total_failureIns_batch4) +
            (isin_batch3 * total_failureIns_batch5) +
            (isin_batch4 * total_failureIns_batch6) +
            (isin_batch5 * total_failureIns_batch7) +
            (isin_batch6 * total_failureIns_batch1) +
            (isin_batch7 * total_failureIns_batch2)) < max_failed_ins'

  proba_fail_insemination:
    desc: 'probability insemination fails. ST cannot failed'
    value: '(1-is_ST) * (is_I + ((1-is_I)* (1/(initial_size_sow))))'

  proba_apply_protoype_batch1:
    desc: 'to limit failure to two per batch. The test "isin" reponds to the old state of the individual, so it is compared to the value of the relevant batches in the completed state'
    value: 'isin_batch1 * ((total_failureIns_batch3) < max_failed_ins )'
  proba_apply_protoype_batch2:
    desc: ''
    value: 'isin_batch2 * ((total_failureIns_batch4) < max_failed_ins )'
  proba_apply_protoype_batch3:
    desc: ''
    value: 'isin_batch3 * ((total_failureIns_batch5) < max_failed_ins )'
  proba_apply_protoype_batch4:
    desc: ''
    value: 'isin_batch4 * ((total_failureIns_batch6) < max_failed_ins )'
  proba_apply_protoype_batch5:
    desc: ''
    value: 'isin_batch5 * ((total_failureIns_batch7) < max_failed_ins )'
  proba_apply_protoype_batch6:
    desc: ''
    value: 'isin_batch6 * ((total_failureIns_batch1) < max_failed_ins )'
  proba_apply_protoype_batch7:
    desc: ''
    value: 'isin_batch7 * ((total_failureIns_batch2) < max_failed_ins )'

  newborn_weight:
    desc: 'Birth weight'
    value: 'random_truncated_normal(1.4,0.37, 0.55, 2.25)'

  GF_maternity:
    desc: 'Growth factor in farrowing (maternity)'
    value: '0.146 - (1.4 - weight) * 0.02'

  GF_pw:
    desc: 'Growth factor in post-weaning and finishing'
    value: '0.0105 - (8 - weight) * 0.00015'

  DWG:
    desc: 'Daily weight gain in post-weaning and finishing'
    value: 'GF_pw * weight * log(270/weight)'

  weight_growth:
    desc: ''
    value: 'IfThenElse(isin_farrowingSector,
    (weight + GF_maternity + (0.041 * weight)),
    (weight + DWG))'

  t0:
    desc: 'time period between vaccination and vaccine-induced immunity'
    value: 56

  exposed_duration:
    desc: 'duration in exposed status for piglets M'
    value: 2

  epsilon_s:
    desc: 'vaccine-induced sterilizing immunity -> reduced transmission through lower host susceptibility'
    value: 0
  epsilon_i:
    desc: 'vaccine-induced reduction pathogen shedding -> reduced transmission through lower host infectivity'
    value: 0
  epsilon_gamma:
    desc: 'vaccine-induced increase in recovery rate -> speed up recovery rate of infected host'
    value: 'IfThenElse(time < t0,0,0.85)'
  d:
    desc: 'death rate due to the infection'
    value: 0.001
  mu:
    desc: 'death rate'
    value:  0
  p:
    desc: 'vaccination coverage'
    value: 0


  beta_nn:
    desc: 'transmission between non-vaccinated (Rose vaccine 2015)'
    value: 0.24
  beta_vn:
    desc: 'transmission between vaccinated and non-vaccinated'
    value: '(1 - epsilon_s) * beta_nn'
  beta_vv:
    desc: 'transmission between vaccinated'
    value: '(1 - epsilon_i) * beta_vn'
  beta_inter_pen:
    desc: 'transmission between pens (Andraud, vet_res 2008)'
    value: 'beta_nn / 10'

  force_of_infection_n:
    desc: 'force of infection for non-vaccinated'
    value: '((beta_nn * info_excret_my_pen)) * IfThenElse((isin_serviceSector), propag_inter, propag_intra)'
  force_of_infection_v:
    desc: 'force of infection for vaccinated'
    value: '((beta_vn * total_I_notVac_my_pen) + (beta_vv * total_I_vac_my_pen))'
  force_of_infection:
    desc: 'force of infection'
    value: '(IfThenElse(is_notVac, force_of_infection_n, force_of_infection_v))/total_my_pen'

  gamma_n:
    desc: 'recovery rate for non-vaccinated (Nodelijk mais 1/22 days Rose et al.)'
    value: 0.01785
  gamma_v:
    desc: 'recovery rate for vaccinated'
    value: 'gamma_n / (1 - epsilon_gamma)'
  gamma:
    desc: 'recovery rate'
    value: 'IfThenElse(is_notVac, gamma_n, gamma_v)'

  shape:
    desc: 'shape of gamma low'
    value: 3.6
  scale:
    desc: 'scale of gamma low'
    value: 1.35
  mda_duration:
    desc: 'maternal immunity duration'
    value: 'random_gamma(shape, scale)'

  lambda_n:
    desc: 'new born rate for non-vaccinated'
    value: 0.0017
  lambda_v:
    desc: 'new born rate for vaccinated'
    value: '(1 - p) * mu'
  lambda:
    desc: 'new born rate'
    value: 'IfThenElse(is_notVac, lambda_v, lambda_n)'

  proportion_I:
    desc: 'proportion of infected'
    value: 'DIV(total_I, total_population)'

weight:
  desc: 'weight of the animal'

tmp:
  desc: ''

dur_in_Ren:
  desc: ''

farrowing_rank:
  desc: 'rank farrowing for culling (if FR > 4)'

nb_failed_insemination:
  desc: 'number of failed insemination. if > 2 then cull'

age:
  desc: ''

agid:
  desc: ''


prototypes:
  population:
    - default_pop:
        desc: ''
  individuals:
    - default_pig:
        desc: ''
        culling: to_keep


    - next_BFR:
        desc: ''
        BFR: next_state

    - gestatingSow:
        desc: 'sow become in its last gestating step'
        gestating_status: first_gestating_period

    - inseminateSow:
        desc: 'determine if insemination succeed or failed'
        inseminationStatus: inseminate

    - newborn_S:
        dur_in_Ren: 0
        desc: 'new born animal S'
        animal_type: Piglet
        physiologicalStep: Farrowing
        gestating_status: nonGestating
        weight: newborn_weight
        production_impact: no_production
        health_state: S
        maternal_immunity: notM
        vaccinated: notVac
    - newborn_I:
        dur_in_Ren: 0
        desc: 'new born animal I'
        animal_type: Piglet
        physiologicalStep: Farrowing
        gestating_status: nonGestating
        weight: newborn_weight
        production_impact: no_production
        health_state: I
        maternal_immunity: notM
        vaccinated: notVac
    - newborn_M:
        dur_in_Ren: 0
        desc: 'new born animal M'
        animal_type: Piglet
        physiologicalStep: Farrowing
        gestating_status: nonGestating
        weight: newborn_weight
        production_impact: no_production
        health_state: S
        maternal_immunity: M
        vaccinated: notVac

    - infected_first:
        desc: 'Sows infected during first period of gestation'
        production_impact: produce_S
    - infected_second:
        desc: 'Sows infected during second period of gestation'
        production_impact: produce_I
    - infected_third:
        desc: 'Sows infected during third period of gestation'
        production_impact: produce_M

    - new_Sow:
        desc: 'new sow for turnover'
        health_state: S
        physiologicalStep: Insemination
        inseminationStatus: successIns
        gestating_status: first_gestating_period
        nb_failed_insemination: 0
        farrowing_rank: 0

    - to_cull_proto:
        desc: ''
        culling: to_cull

    - failure_insemination_batch1:
        desc: 'for failure insemination'
        gestating_status: first_gestating_period
        physiologicalStep: Insemination
        organization: [batches, litter, pen_group, housing]
        init_location:
          batches: 'batch3'
    - failure_insemination_batch2:
        desc: 'for failure insemination'
        gestating_status: first_gestating_period
        physiologicalStep: Insemination
        organization: [batches, litter, pen_group, housing]
        init_location:
          batches: 'batch4'
    - failure_insemination_batch3:
        desc: 'for failure insemination'
        gestating_status: first_gestating_period
        physiologicalStep: Insemination
        organization: [batches, litter, pen_group, housing]
        init_location:
          batches: 'batch5'
    - failure_insemination_batch4:
        desc: 'for failure insemination'
        gestating_status: first_gestating_period
        physiologicalStep: Insemination
        organization: [batches, litter, pen_group, housing]
        init_location:
          batches: 'batch6'
    - failure_insemination_batch5:
        desc: 'for failure insemination'
        physiologicalStep: Insemination
        gestating_status: first_gestating_period
        organization: [batches, litter, pen_group, housing]
        init_location:
          batches: 'batch7'
    - failure_insemination_batch6:
        desc: 'for failure insemination'
        gestating_status: first_gestating_period
        physiologicalStep: Insemination
        organization: [batches, litter, pen_group, housing]
        init_location:
          batches: 'batch1'
    - failure_insemination_batch7:
        desc: 'for failure insemination'
        gestating_status: first_gestating_period
        physiologicalStep: Insemination
        organization: [batches, litter, pen_group, housing]
        init_location:
          batches: 'batch2'

    - new_sow:
        desc: ''
        nb_failed_insemination: 0
        farrowing_rank: 0
        physiologicalStep: Insemination
        health_state: S


    - sow1:
        inseminationStatus: successIns
        farrowing_rank: "random_integers(0,5)"
        nb_failed_insemination: 0
        desc: 'sow'
        animal_type: Sow
        physiologicalStep: Insemination
        gestating_status: first_gestating_period
        health_state: S
        BFR: BFR1
        production_impact: produce_S
        maternal_immunity: notM
        vaccinated: notVac
        organization: [batches, litter, pen_group, housing]
        init_location:
          batches: 'batch1'

    - sowST:
        desc: 'sow ST'
        health_state: ST
        farrowing_rank: 0
        nb_failed_insemination: 0

    - sow2:
        inseminationStatus: successIns
        farrowing_rank: "random_integers(0,5)"
        nb_failed_insemination: 0
        desc: 'sow'
        animal_type: Sow
        physiologicalStep: Quarantine
        gestating_status: nonGestating
        health_state: S
        BFR: BFR0
        production_impact: produce_S
        maternal_immunity: notM
        vaccinated: notVac
        organization: [batches, litter, pen_group, housing]
        init_location:
          batches: 'batch2'

    - sow3:
        inseminationStatus: successIns
        farrowing_rank: "random_integers(0,5)"
        nb_failed_insemination: 0
        desc: 'sow'
        animal_type: Sow
        physiologicalStep: Quarantine
        gestating_status: nonGestating
        health_state: S
        BFR: BFR0
        production_impact: produce_S
        maternal_immunity: notM
        vaccinated: notVac
        organization: [batches, litter, pen_group, housing]
        init_location:
          batches: 'batch3'

    - sow4:
        inseminationStatus: successIns
        farrowing_rank: "random_integers(0,5)"
        nb_failed_insemination: 0
        desc: 'sow'
        animal_type: Sow
        physiologicalStep: Quarantine
        gestating_status: nonGestating
        health_state: S
        BFR: BFR0
        production_impact: produce_S
        maternal_immunity: notM
        vaccinated: notVac
        organization: [batches, litter, pen_group, housing]
        init_location:
          batches: 'batch4'

    - sow5:
        inseminationStatus: successIns
        farrowing_rank: "random_integers(0,5)"
        nb_failed_insemination: 0
        desc: 'sow'
        animal_type: Sow
        physiologicalStep: Quarantine
        gestating_status: nonGestating
        health_state: S
        BFR: BFR0
        production_impact: produce_S
        maternal_immunity: notM
        vaccinated: notVac
        organization: [batches, litter, pen_group, housing]
        init_location:
          batches: 'batch5'

    - sow6:
        inseminationStatus: successIns
        farrowing_rank: "random_integers(0,5)"
        nb_failed_insemination: 0
        desc: 'sow'
        animal_type: Sow
        physiologicalStep: Quarantine
        gestating_status: nonGestating
        health_state: S
        BFR: BFR0
        production_impact: produce_S
        maternal_immunity: notM
        vaccinated: notVac
        organization: [batches, litter, pen_group, housing]
        init_location:
          batches: 'batch6'

    - sow7:
        inseminationStatus: successIns
        farrowing_rank: "random_integers(0,5)"
        nb_failed_insemination: 0
        desc: 'sow'
        animal_type: Sow
        physiologicalStep: Quarantine
        gestating_status: nonGestating
        health_state: S
        BFR: BFR0
        production_impact: produce_S
        maternal_immunity: notM
        vaccinated: notVac
        organization: [batches, litter, pen_group, housing]
        init_location:
          batches: 'batch7'


initial_conditions:
  population:
    - prototype: sow1
      amount: 'initial_size_sow'
    - prototype: sow2
      amount: 'initial_size_sow'
    - prototype: sow3
      amount: 'initial_size_sow'
    - prototype: sow4
      amount: 'initial_size_sow'
    - prototype: sow5
      amount: 'initial_size_sow'
    - prototype: sow6
      amount: 'initial_size_sow'
    - prototype: sow7
      amount: 'initial_size_sow'


outputs:
  type: csv
  population:
    period: 1
    extra_vars:
      - total_I_batch1
      - total_I_batch2
      - total_I_batch3
      - total_I_batch4
      - total_I_batch5
      - total_I_batch6
      - total_I_batch7

      - total_I_serviceSector
      - total_I_gestatingSector
      - total_I_farrowingSector
      - total_I_nurserySector
      - total_I_finishingSector
...
```
