## Supplementary material for "Pig herd management and infection transmission dynamics: a challenge for modellers": supplementary.pdf

### Additional file

The following file (to be renamed as "SDRP_like.yaml") is meant to be run with the EMULSION framework (version 1.3beta1) which can be installed according to the instructions provided on the software webpage: https://sourcesup.renater.fr/www/emulsion-public/. The syntax of the EMULSION modelling language is also fully described there.

The installation of EMULSION, requires the following command:

```
sudo pip3 install emulsion==1.3beta1
```

Then for running a scenario, e.g. 'sdrp_like_output' for 1 stochastic repetition, requires the following command:

```
emulsion run SDRP_like.yaml -r 1 --output-dir sdrp_like_output
```

Simulation results are stored in directory 'sdrp_like_output' in a CSV file named 'counts.csv'.
